# RealTrace: Uncovering biological dynamics hidden under measurement noise in time-lapse microscopy data

**DOI:** 10.1101/2025.09.12.675772

**Authors:** Bjoern Kscheschinski, Athos Fiori, Dany Chauvin, Benjamin Martin, David Suter, Benjamin Towbin, Thomas Julou, Erik van Nimwegen

## Abstract

One of the most powerful approaches for identifying the mechanisms underlying complex biological phenomena is not just to measure bulk populations, or even take single-cell snapshots, but to directly track the behaviors of single cells *in time*. Indeed, in recent years there has been a steep rise in the use of time-lapse microscopy with fluorescent reporters to quantify single-cell time dynamics in many different biological systems.

However, there is a major challenge with the analysis of such data. Since biological changes are inherently small on short time scales, much of the true biological dynamics is hidden under unavoidable measurement noise, but we do not know what functional form the time dynamics may take, and smoothing over neighboring time points simply replaces the true dynamics with a time average.

Here, we present RealTrace, a Bayesian method that rigorously solves this challenge using only the assumption that, while true biological fluctuations are correlated on short time scales, measurement errors are uncorrelated. We show that this assumption can be formalized in a Bayesian model using maximum entropy process priors and that this model can be accurately solved by recursively approximating the non-linear dynamics over short time intervals. Given raw input cell size and fluorescence measurements, RealTrace calculates the estimated true sizes, total GFP contents, instantaneous growth rates, and volumic GFP production rates across all cells and time, together with rigorous error bars on all these quantities.

We demonstrate the broad applicability of RealTrace by analyzing time-lapse microscopy datasets from *E. coli* bacteria, mouse embryonic stem cells, and entire *C. elegans* embryos. To exemplify the subtle dynamical features RealTrace can uncover, we perform an in-depth study of *E. coli* cells carrying fluorescent reporters for constitutive and ribosomal promoters, across multiple growth conditions. We find growth rates of single cells substantially fluctuate in time and, as the average growth rate decreases, these fluctuations systematically increase in amplitude and duration. Both growth rate and gene expression vary systematically across the cell cycle, but while the pattern of growth rate variation is condition-independent, the systematic variation in gene expression across the cell cycle is both condition and promoter-dependent. Finally, under a sudden change in nutrients, single cells exhibit highly consistent transient changes in growth rate and gene expression. While gene expression rates settle into a new steady-state surprisingly rapidly, the growth rate shows a dramatic drop followed by an overshoot that takes several cell cycles to settle down. All these observations give fundamental new insights into the single-cell physiology of bacteria.

RealTrace is a highly versatile method that accurately removes measurement noise from any fluorescence time-lapse microscopy data, enabling the identification and quantification of subtle features in the biological dynamics that would otherwise be hidden under measurement noise.

## 1 Introduction

Due to technical limitations, most measurements of biological systems have traditionally concerned snapshots of bulk populations. However, when single-cell measurement technologies became available, it became clear that bulk measurements can easily give a highly misleading picture, and that single-cell measurements are far more powerful for uncovering underlying biological measurements. For example, while bulk measurements could suggest that the expression of a gene increases smoothly upon induction, single-cell measurements might reveal that individual cells have either low or high expression, and that only the *fraction* of cells in the high expression state increases with induction ^1,2^.

Next, when methods became available for tracking the dynamics of single cells, e.g., time-lapse microscopy with fluorescent reporters, it became clear that even single-cell snapshots can give a misleading picture of the dynamics at the single-cell level. To continue the example, while single-cell snapshots may suggest two subpopulations of cells with either low or high expression, time-lapse data may unveil that, in actuality, all cells are stochastically switching between a low expression state, and transient pulses of high expression, and that upon induction, these pulses become more frequent^3,4^. This observation yet again completely changes the interpretation of the underlying regulatory mechanisms ^5–7^.

And thus, not surprisingly, there has been a steep rise in the use of technologies that allow tracking of the behavior of single cells through time. This has led to numerous novel insights into regulatory mechanisms, including the dynamics of the cell cycle in bacteria ^8–10^ and eukaryotes ^11,12^, control of body size ^13^, cross-regulation between gene expression and growth ^14^, and the discovery that regulation is often implemented through stochastic pulsing or oscillations in regulator activity ^5,7,15–19^.

The current frontier in the tracking of the behavior of biological objects such as single cells (or nuclei, organs, compartments, or even whole embryos) is to make these observations quantitatively precise, so that they can be meaningfully compared with mathematical models that predict quantitative features such as: how fast are the responses to a change in condition, what is the amplitude of the response, how much do response times and amplitudes vary across cells, what is the extent and time scale of stochastic fluctuations within a single condition, and so on. For example, for time-lapse fluorescence microscopy data, this requires precise quantification of how much the size and fluorescence of the biological object change from one time point to the next.

Unfortunately, we cannot simply take the measured changes in size and fluorescence to estimate the true dynamics because there is substantial measurement noise that needs to be taken into account. Crucially, since the true state of biological systems varies continuously in time, on sufficiently short time scales, the true biological changes become smaller than the size of the measurement noise. Moreover, there are inherent physical limits on measurement accuracy, e.g., in fluorescence measurements, there is unavoidable shot noise in photon counts. Consequently, in most time-lapse fluorescence microscopy data, the true biological changes on short time scales are hidden under measurement noise, which is a very challenging problem.

In many recent works, measurement noise is dealt with by some sort of smoothing or averaging of measurements across local windows ^13,14,20–23^. But this fundamentally distorts the true biological dynamics, i.e., it replaces the real dynamics with a time average, which removes changes on short time scales and systematically attenuates fluctuations. We also cannot fit the measurements to some definite mathematical function because we do not *a priori* know what kind of form the dynamics might take. The challenge is to recover the true biological dynamics under the measurement noise without making specific assumptions about the functional form that the time dynamics takes.

Here, we show that this problem can be rigorously solved by exploiting that, while the true biological changes are correlated on short time scales, measurement errors are all independent of each other. The assumption that the underlying biological dynamics is smooth can be formalized using maximum entropy process priors, which generalize the maximum entropy method to time-dependent processes ^24,25^. The maximum entropy process prior characterizes the time dynamics of biological quantities such as growth only by their means, variances, and correlation time scales. We show how these maximum entropy process priors can be combined with statistical models of the measurement errors in a Bayesian procedure that infers the true underlying biological dynamics, correcting for the measurement errors, and providing rigorous error bars on all the estimates.

In particular, we here focus on time-lapse fluorescence microscopy data where the total size and fluorescence of a biological object such as a cell, a nucleus, or even a whole organism, is tracked in time. In most applications, the fluorescence either reports on the expression of a gene of interest, or some other cellular activity. We then parametrize the dynamics of size and fluorescence by unknown time-dependent rates of growth and volumic fluorescence production (i.e., the production of fluorescence per unit time and volume), so that in the simplest scenario of constant rates, the biological objects grow perfectly exponentially with time and the fluorescence per unit volume remains constant. From the measured time traces of size and total fluorescence, we then infer the time-dependent growth and volumic production rates across all cells and times. We show that, although the time traces of measured size and total fluorescence depend in a complex, non-linear manner on the fluctuating growth and volumic fluorescence production rates, the model can nonetheless be effectively solved by linearizing the non-linear dynamics over the short time intervals between measurements. In particular, we derive a recursive algorithm analogous to the forward/backward algorithm of hidden Markov models to calculate the likelihood and posterior distributions.

We implemented these methods in a tool, called RealTrace, that takes raw measurements of size and total fluorescence across a lineage tree of cells (or nuclei, or any other biological objects) and first finds the parameters of the maximum entropy process prior that maximizes the likelihood, and then infers posterior distributions of size, total fluorescence, growth rate, and volumic fluorescence production for each object and time point.

We demonstrate RealTrace’s wide applicability by applying it to traces of *C. elegans* larvae, mouse embryonic stem cells, and *E. coli* cells, and show that RealTrace uncovers many quantitative details of the dynamics that were hidden under measurement noise and cannot be recovered by simple averaging approaches. To exemplify the kind of dynamical features that RealTrace can uncover, we perform an in-depth analysis of datasets from *E. coli* cells growing in a microfluidic device^26,27^ in different growth conditions, and carrying fluorescent reporters of both ‘unregulated’ constitutive promoters and promoters of highly regulated ribosomal genes.

We find that growth rates vary substantially in time and from cell to cell and that these fluctuations can last for multiple cell cycles. Moreover, we find that these growth rate fluctuations become larger and more long-lasting in slower growth conditions. Second, we find that hidden under the large cell-to-cell variations, there are modest but consistent variations in growth rate and volumic GFP production rate across the cell cycle. Independent of growth condition, growth rate exhibits a minimum near the start of the cell cycle and a maximum near the end of the cell cycle. In contrast, the pattern of volumic GFP production across the cell cycle varies both across growth conditions and across promoters. Strikingly, we find that the variation in volumic GFP production across the cell cycle is different for different constitutive promoters, even though these promoters are not thought to contain regulatory sites beyond the binding site of the RNA polymerase. Finally, we find that upon a sudden change in nutrients, the growth and volumic GFP production rates show highly consistent changes across single cells. In particular, growth rates show an almost instantaneous dramatic drop, which is followed by an overshoot that lasts several cell cycles to relax to the final average growth rate. In contrast, the volumic GFP production quickly settles to a new equilibrium frequency for all promoters.

These observations provide important novel insights into the physiology and regulatory dynamics of single bacterial cells that, as far as we are aware, are not explained by any current models of growth and gene expression.

## 2 Results

### 2.1 Maximum entropy process priors formalize the assumption of smoothly varying biological states

We focus on datasets consisting of time courses of the measured size and total fluorescence of biological objects such as cells, nuclei, or even whole organisms. To simplify terminology, we will assume in the presentation of the methods that the data consist of time courses of the sizes and fluorescence of single cells. We start by formalizing the idea that the true size and total fluorescence of each cell vary smoothly with time.

We first need to choose a parametrization of the time dynamics of cell size and fluorescence. It is well known that for many biological objects, the increase in size per unit time is approximately proportional to their current size, so that size increases approximately exponentially with time. This approximately exponential growth applies to bacteria ^14,26,28–31^, budding yeast ^23,32^, mammalian cells ^33,34^, and even *C. elegans* larvae^13^. We thus choose a parametrization, where, in the simplest ‘constant’ scenario, the growth is perfectly exponential. That is, we parametrize the dynamics of the cell size *s*(*t*) by a growth-rate *λ*(*t*)

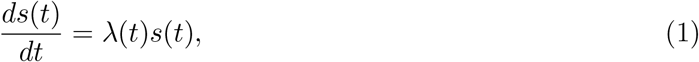

such that when *λ*(*t*) is constant in time, *s*(*t*) increases exponentially. Note that this parametrization still allows for any time-dependent dynamics of the size *s*(*t*), including decreases in size, which occurs for negative *λ*(*t*). This parametrization merely implies that the ‘simplest’ dynamics is exponential growth and that deviations from this are encoded into a fluctuating growth rate *λ*(*t*).

Note that the logarithm of size *x*(*t*) = log *s*(*t*) follows the simple dynamics

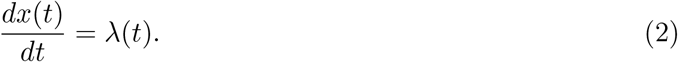

Similarly, for the dynamics of total fluorescence, the simplest assumption is that the amount of fluorescence that is produced per unit time (which for typical reporter constructs is proportional to the number of GFP molecules produced per unit time) is proportional to the current size of the cell. That is, we parametrize the dynamics of total fluorescence *g*(*t*) by a volumic fluorescence production rate *q*(*t*):

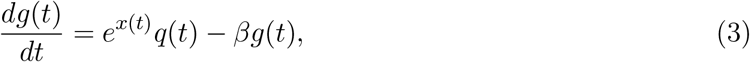

where we have already taken into account that fluorescence decays at some (unknown) rate *β* due to photobleaching and protein decay. Note that under this assumption, if the growth rate *λ*(*t*) and production rate *q*(*t*) are constant in time, the fluorescence per unit volume *g*(*t*)*/s*(*t*) stays constant. For bacterial cells carrying GFP reporter constructs, a simple biophysical interpretation of the volumic production rate *q*(*t*) would be that it is proportional to the concentration of active ribosomes times their average translation rate times the fraction of ribosomes that are translating GFP mRNAs (which in turn would be proportional to the amount of GFP mRNA). Consequently, fluctuations in *q*(*t*) could then be interpreted as fluctuations in either GFP mRNA, ribosome concentration, or average translation rate.

We do not want to make any specific assumptions about the dynamics of growth rate *λ*(*t*) and volumic production rate *q*(*t*) beyond that they vary smoothly in time. To formalize this assumption, we will characterize these time-dependent rates only through their means, variances, and correlation times. The ‘most random’ or ‘least assuming’ distribution over possible continuous curves with a given mean, variance, and correlation time corresponds to the maximum entropy process with a given mean, variance and correlation time ^35,36^. It can be shown that the maximum entropy process priors are equivalent to so called Ornstein-Uhlenbeck processes ^37^ which are given by the Langevin equations,

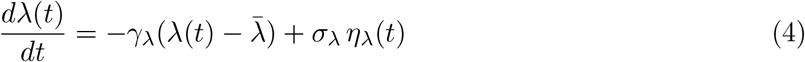

and

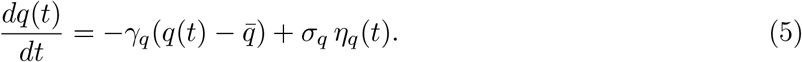

Here, 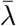 and 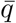 are the mean rates, 1*/γ*_*λ*_ and 1*/γ*_*q*_ the correlation times, *η*_*λ*_(*t*) and *η*_*q*_(*t*) are both white noise with variance 1, and *σ*_*λ*_ and *σ*_*q*_ set the size of the noise. Notably, the steady-state distributions of *λ*(*t*) and *q*(*t*) are Gaussian distributed with variances 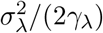 and 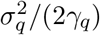, respectively.

In many datasets, cells are tracked over multiple generations so that our prior process should also describe what occurs at cell divisions. We will take into account that, although on average cells divide in half, there are fluctuations in the sizes of the daughters, and that GFP molecules are stochastically divided between the two daughters. In particular, we assume the log size of the daughter cells is drawn from a Gaussian centered around half the cell size of the parent cell with (unknown) variance 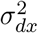, and each GFP molecule is randomly assigned to one of the daughters with probability proportional to the respective size of the daughter cell. Note that this requires that we know the amount of fluorescence per GFP molecule. Although the conversion factor between fluorescence and GFP number can be obtained through calibration experiments ^27^, for many datasets, this will be unknown, and we therefore estimate this conversion factor from the data as well.

Since the measurement noise of the fluorescence follows shot noise, we assume the fluorescence measurement noise is proportional to 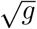 with an unknown prefactor *σ*_*g*_. Finally, we assume the measurement noise of the log size *x* is Gaussian with an unknown standard deviation *σ*_*x*_.

In summary, the full set of parameters characterizing the prior process and measurement noise is 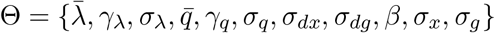. In the next sections, we show how the maximum entropy process prior is employed to generate prior probabilities and infer the true cell state comprised of log size, GFP content, growth, and volumic production given noisy measurements of cell size and GFP content.

### 2.2 An iterative procedure for calculating posterior distributions of cell states

At each time point *t*_*n*_, we have a measurement of the size and total fluorescence 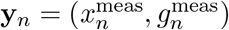 and from these noisy measurements, we want to reconstruct the best estimates of the true underlying states of the cell **z**_*n*_ = [*x*_*n*_, *g*_*n*_, *λ*_*n*_, *q*_*n*_] at each time point. To this end, we use a Bayesian procedure that combines the measurements with our maximum entropy process prior that quantifies, for each neighboring pair of time points *t*_*n*_ and *t*_*n*+1_, the probability *P* (**z**_*n*+1_|**z**_*n*_, Θ) for the cell’s state to evolve from **z**_*n*_ to **z**_*n*+1_ (which depends on the parameters Θ of the process).

We denote by *P* (**z**_*n*_|*D*_1,…,*n*_, Θ) the posterior probability of the state **z**_*n*_ at time point *n*, given both all measurements up to time point *n* and the parameters Θ of the prior process. Similarly, we denote by *P* (**y**_*n*+1_|*D*_1,…,*n*_, Θ) the probability of the measurements **y**_*n*+1_ at time point *t*_*n*+1_ given all previous *n* measurements. As sketched in Fig. 1, assuming the parameters of the process prior Θ are given, these probabilities can be efficiently calculated using an iterative procedure.

**Figure 1:**
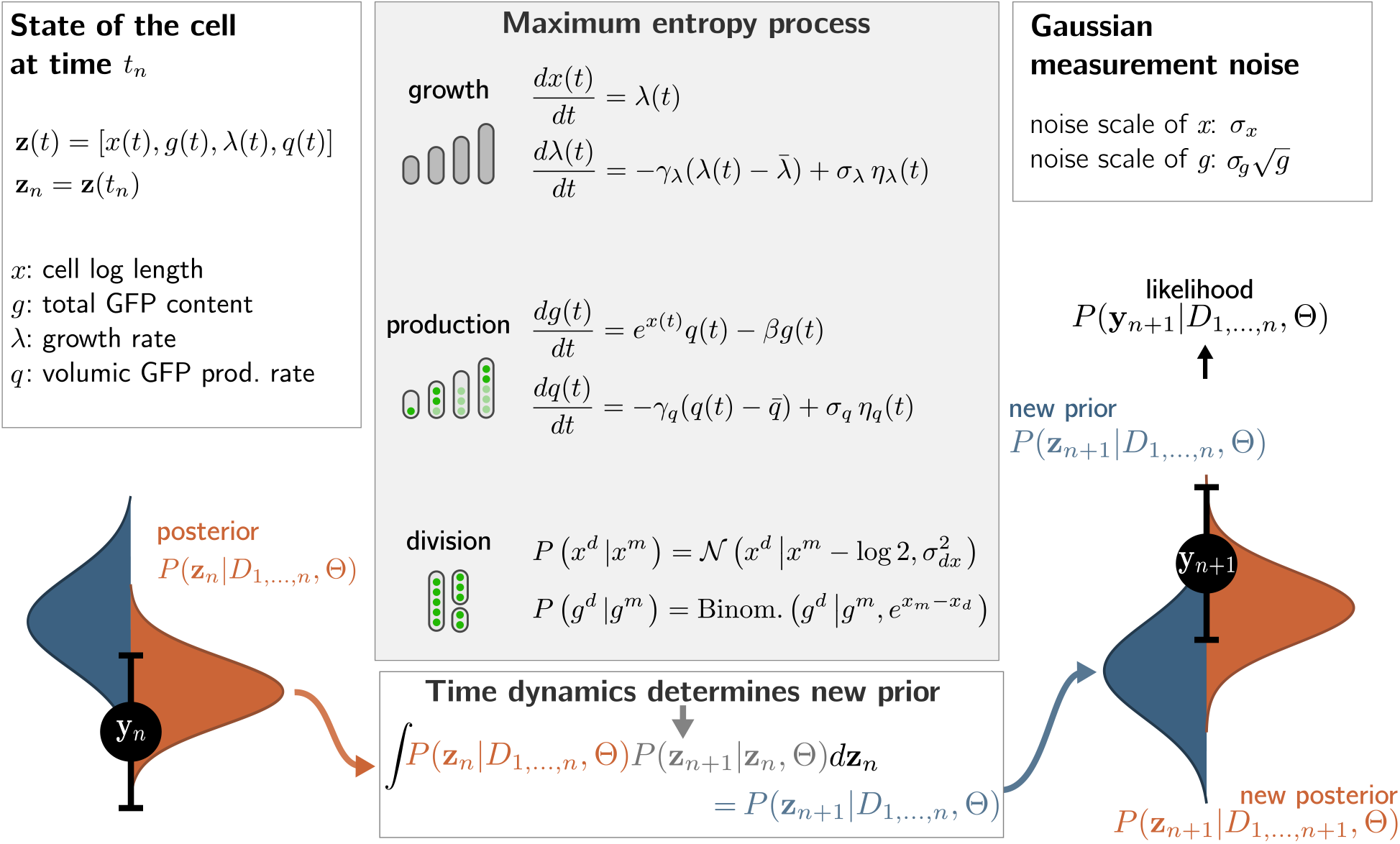
Summary of the iterative procedure for combining the maximum entropy process prior with measurements to obtain posterior distributions over cell states. The state of the cell at the time of the *n*th measurement *z*_*n*_ is described by its log size *x*_*n*_, total fluorescence *g*_*n*_, growth rate *λ*_*n*_ and volumic fluorescence production rate *q*_*n*_ (top left). We denote by *P* (**z**_*n*_ |*D*_1,…,*n*_, Θ) (red, bottom left) the posterior of the cell state conditioned on measurements 1 through *n* and the parameters of the process prior Θ. The maximum entropy process prior (middle) defines the probability *P* (**z**_*n*+1_ |**z**_*n*_, Θ) (bottom, grey) that the cell’s state evolves from **z**_*n*_ to **z**_*n*+1_ in the short interval from measurement *n* to *n*+1. By marginalizing over **z**_*n*_ (middle bottom) we obtain the probability *P* (**z**_*n*+1_ |*D*_1,…,*n*_, Θ) (blue) for the state at time *n* + 1 given the first *n* measurements. Finally, by incorporating the likelihood of measurement **y**_*n*+1_ (black, right) we obtain the posterior *P* (**z**_*n*+1_ | *D*_1,…,*n*+1_, Θ) (bottom right). This procedure can be iterated to obtain the likelihood of the entire dataset given Θ as well as posteriors for **z**_*n*_ at each time point.

In particular, starting from the posterior *P* (**z**_*n*_|*D*_1,…,*n*_, Θ) of the cell’s state given the first *n* measurements, we use the prior process to calculate the probability *P* (**z**_*n*+1_|**z**_*n*_, Θ) that the cell’s state evolves to **z**_*n*+1_ in the short interval from *t*_*n*_ to *t*_*n*+1_. Multiplying these probabilities and integrating over **z**_*n*_ we obtain the probability *P* (**z**_*n*+1_|*D*_1,…,*n*_, Θ) for the state at *t*_*n*+1_ given the first *n* measurements. Multiplying *P* (**z**_*n*+1_|*D*_1,…,*n*_, Θ) by the probability *P* (**y**_*n*+1_|*z*_*n*+1_, Θ) of the measurements at time *t*_*n*+1_and integrating over **z**_*n*+1_, we then obtain the probability *P* (**y**_*n*+1_|*D*_1,…,*n*_, Θ) of the data at time *t*_*n*+1_ given all previous measurements. We similarly obtain the posterior probability *P* (**z**_*n*+1_|*D*_1,…,*n*+1_, Θ) up to time point *t*_*n*+1_. In this way, posterior distributions for the cell’s state given all ‘past’ measurements are calculated for each time point. As explained in the Methods section 4.1, the posterior distribution *P* (**z**_*n*_|*D*, Θ) given *all* data can be written as a production of the posterior *P* (**z**_*n*_|*D*_1,…,*n*_, Θ) given all past measurements and a posterior *P* (**z**_*n*_|*D*_*n*+1,…,*N*_, Θ) given all future measurements, and the probability *P* (**z**_*n*_|*D*_*n*+1,…,*N*_, Θ) can be obtained using a similar recursive procedure.

Because the time dynamics of the total fluorescence *g*(*t*) depend on the dynamics of the log size *x*(*t*) in a non-linear manner, i.e. equation (3), the Ornstein-Uhlenbeck process of *q*(*t*) is non-linearly coupled to the log size *x*(*t*), which is itself the integral of the Ornstein-Uhlenbeck process for *λ*(*t*). This makes the calculation of the priors *P* (**z**_*n*+1_|**z**_*n*_, Θ) analytically intractable, as well as the integrals needed for our recursive calculation of likelihoods and posteriors. To make this task analytically feasible, we use two key approximations. First, we approximate the conditional probability *P* (**z**_*n*+1_|**z**_*n*_, Θ) and the prior distributions as Gaussians characterized by their means and covariance matrices. Second, since the time increment between measurements of time-lapse microscopy is typically short, the error of ignoring the impact of growth rate fluctuations between measurements on the GFP dynamics is very small. Therefore, to solve the integral and obtain the new prior, we approximate the GFP dynamics *g*(*t*) by approximating the growth rate as constant between *t*_*n*_ and *t*_*n*+1_, i.e., neglecting fluctuations in growth rate between consecutive time points. Then, employing Wick’s probability theorem, we calculate the mean vector and the covariance matrix of the new prior distribution (see Supplementary Materials Sec. E.7). Note that, because the distributions *P* (**z**_*n*+1_|**z**_*n*_, Θ) are approximated as multivariate Gaussians, all the posteriors *P* (**z**_*n*_|*D*_1,…,*n*_, Θ), and likelihoods *P* (**y**_*n*_|**z**_*n*_, Θ) are multivariate Gaussians as well, so that all integrals can be performed analytically. Below we confirm the accuracy of our approximations using realistic simulation data.

The overall inference procedure employed by RealTrace is as follows. Using the iterative procedures, we calculate the probabilities *P* (**y**_*n*+1_|*D*_1,…,*n*_, Θ) for each measurement, and multiplying these for all measurements, we obtain the likelihood of the entire dataset. RealTrace then uses numerical optimization to find the set of parameters Θ^*∗*^ that maximizes the likelihood of the dataset. Using these optimal parameters Θ^*∗*^, RealTrace then calculates posterior distributions *P* (**z**_*n*_|*D*, Θ^*∗*^) for the state of each cell at each time point. A longer description of the procedures are described in the Methods (Sec. 4) and all technical details are described in the Supplementary Materials.

### 2.3 Parameters of the prior process and the measurement noise are accurately estimated

We first tested the accuracy of our inference procedure using synthetic datasets. To make sure we use plausible parameters Θ for the prior process and measurement noise, we took the parameters inferred from data of *E. coli* cells growing in a so called ‘Dual Input Mother Machine’ microfluidic device^27^ that we analyze below as a reference set of parameters (see Tab. 1). We then created 100 datasets with randomly chosen parameters Θ in a wide range around these reference values (Tab. 2). We then created 100 datasets by simulating 1000 cell cycles for each of the 100 parameter settings (see Supplementary Materials Sec. E.1 for details).

**Table 1:** Parameters for the simulation of the synthetic data set mimicking the dynamics of *E. coli* growing on glucose and expressing GFP under the control of *hi1*. Note that the bleaching rate is assumed to be 0.

| Parameter | Value | Unit |
| --- | --- | --- |
| $\bar{\lambda}$ | 0.0114 | (1/min) |
| $\gamma_{\lambda}$ | 0.167 | (1/min) |
| $\sigma_{\lambda}^2$ | $4.95 \times 10^{-6}$ | (1/min<sup>2</sup>) |
| $\bar{q}$ | 119 | (1/$\mu\text{m}$/min) |
| $\gamma_q$ | 0.0596 | (1/min) |
| $\sigma_q^2$ | 430 | (1/$\mu\text{m}$/min<sup>2</sup>) |
| $\beta$ | 0 | (1/min) |
| $\sigma_x^2$ | $1.37 \times 10^{-4}$ | |
| $\sigma_g^2$ | 12.3 | |
| $\sigma_{dx}^2$ | 0.00469 | |

**Table 2:** Parameter ranges from which parameter sets are sampled for the simulated datasets.

| Parameter | Min | Max | Unit |
| --- | --- | --- | --- |
| $\bar{\lambda}$ | 0.005 | 0.05 | (1/min) |
| $\gamma_{\lambda}$ | 0.05 | 0.5 | (1/min) |
| CV | 0.2 | 0.8 |  |
| $\bar{q}$ | 100 | 1000 | (1/$\mu\text{m}/\text{min}$) |
| $\gamma_q$ | 0.05 | 0.5 | (1/min) |
| CV($q$) | 0.2 | 0.8 | |
| $\beta$ | $10^{-4}$ | $10^{-3}$ | (1/min) |
| $\sigma_x^2$ | $2 \times 10^{-5}$ | $2 \times 10^{-4}$ | |
| $\sigma_g^2$ | 1.5 | 15 | |
| $\sigma_{dx}^2$ | $10^{-4}$ | $10^{-2}$ | |
| $\sigma_{dg}^2$ | 1 | 100 | |

We note that, because some of the parameters, e.g., the bleaching rate *β*, might be known from separate calibration experiments, RealTrace allows the user to fix known parameters so that only the unknown parameters are optimized. In addition, bounds for each parameter can be set to ease the numerical search if the initial parameters are far away from the optimum. However, in the experiments reported here, no bounds were set.

We initialized the numerical optimization from a random set of parameters (with distributions shown in Fig. S1). We find that the maximum likelihood estimates Θ^*∗*^ closely recover the true parameters that were used for each of the simulated datasets without any systematic bias, and typically within a few percent of the true value for all of the parameters (Fig. 2**a**). Notably, the sizes of the measurement noise of cell size *σ*_*x*_ and GFP content *σ*_*g*_ were estimated very precisely, i.e. within 2 % of the true values.

**Figure 2:**
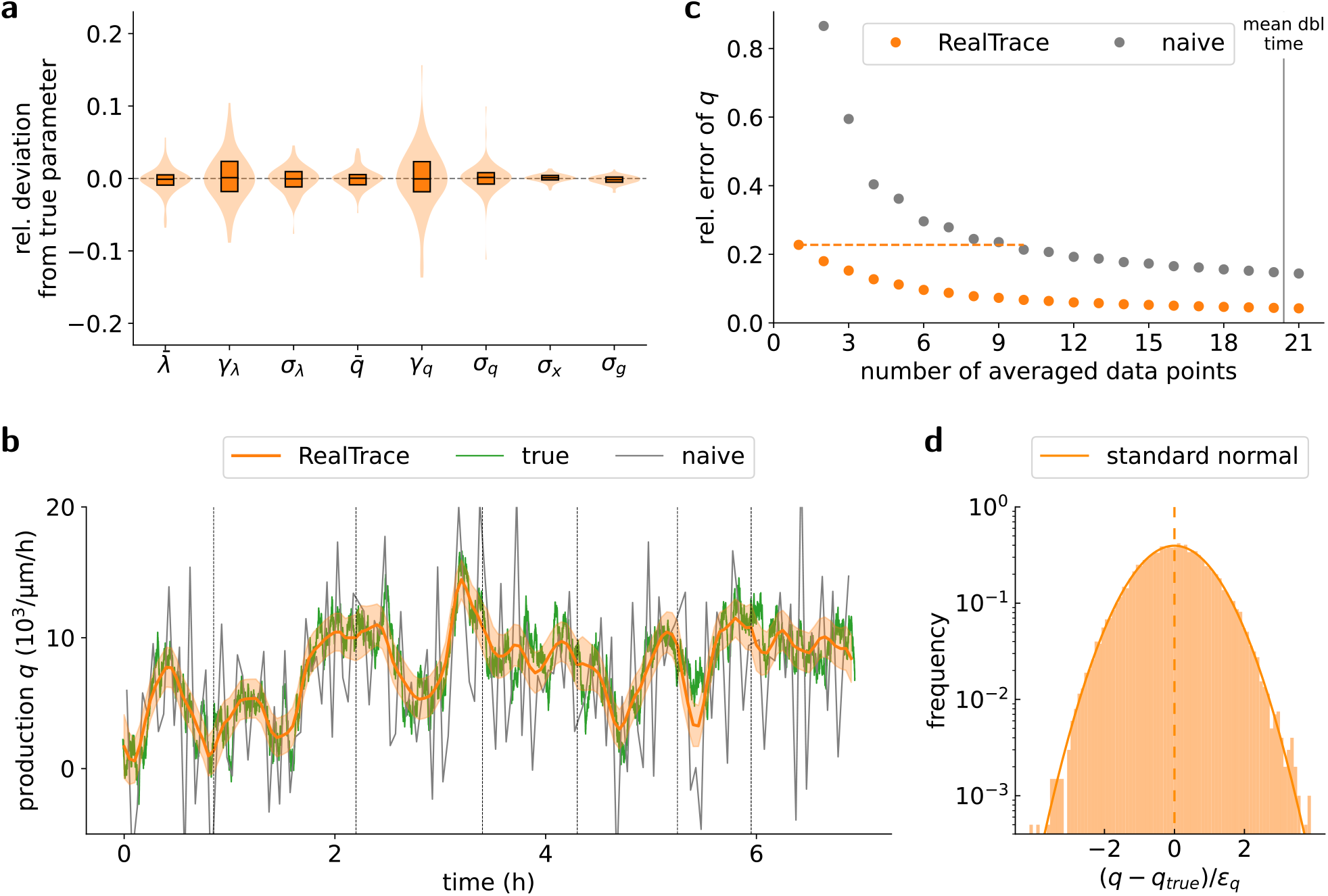
Performance of RealTrace on synthetic data with known ground truth. **a** Accuracy of parameter estimates for 100 synthetic data sets with randomly drawn parameters. For each parameter Θ^*i*^ the relative deviation is calculated as 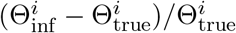 and the figure shows medias (lines), interquartile ranges (boxes), and the distribution of deviations (violin plots) for each of the parameters across the 100 datasets. We see that the optimization of the likelihood *P* (*D* |Θ) accurately recovers the parameters used to generate the datasets. **b** Example estimates of the volumic GFP production rate *q* (mean and standard deviation of the posterior as orange line and ribbon) compared to the ground truh (green) and estimates calculated from naively taking differences between time points (grey). **c** Relative errors 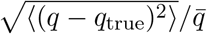 in the estimate of the average volumic production rate for time windows of different length for ReaTrace (orange) and naive estimates from differences across the time window (grey). The vertical line around 21 time points marks the mean doubling time. **d** Distribution of the ratio (*q* − *q*_*true*_)*/ε*_*q*_, which corresponds to the deviation of the estimated volumic production rate *q* from the ground truth *q*_*true*_ relative to the error bar *ε*_*q*_ that RealTrace reports (given by the standard deviation of the posterior). Note that the distribution of (*q* − *q*_*true*_)*/ε*_*q*_ almost perfectly matches a standard normal, implying that the estimates for the production rate neither systematically over-nor underestimate the true production rate and that the error bars are accurate. The results of panels b-d are for a dataset generated with the reference parameters of table 1.

Although cell divisions can, of course, occur at any point in time, in our inference procedure, we make the simplifying assumption that cell divisions always occur exactly at measurement time points. As a consequence, we systematically underestimate the parameters *σ*_*dx*_ and *σ*_*dg*_ that quantify the cell division noise (Fig. S2). However, this does not compromise the accuracy of the posterior distributions of the cell states, as we show in the next section.

These results suggest that the parameters Θ of the prior process and measurement noise can be very robustly recovered and that the optimization of these parameters is not generally hindered by local optima in the likelihood surface. However, it is important to stress that the parameters Θ^*∗*^ are not the main objective of the inference. The role of the process prior is solely the distribution *P* (**z**_*n*+1_|**z**_*n*_), i.e., to indicate how much the current cell’s state is likely to change over the short time interval between measurement time points. The main objective is to infer the states of the cells and provide error bars on these estimates. In the next section, we assess RealTrace’s accuracy on this task.

### 2.4 Posterior distributions reveal true underlying processes

The inference procedure of RealTrace calculates posterior distributions for the state of the cell at each time point that are given by multi-variate Gaussians. The mean values of these posteriors represent the best guesses for the true values of cell log size, total GFP content, growth rate, and volumic production rate, while their standard deviations provide natural error bars for the estimates. To estimate RealTrace’s accuracy we used a synthetic dataset that was generated with parameters (Tab. 1) matching those inferred from a dataset of *E. coli* cells growing in a microfluidic device (see Supplementary Materials Sec. E.1 for details, and example traces of the synthetic data in Fig. S3).

We here focus on the accuracy of the estimates of the volumic production rate *q* as this variable is arguably most difficult to infer, i.e., it is not directly observed and suffers from measurement noise in both cell size and GFP content. Fig. 2**b** shows that the posteriors of *q* accurately track the true dynamics, and are far more accurate than naive estimates based on fluorescence differences between consecutive time points (analogous results for growth rate are shown in Fig. S4). The reader may note that the true volumic production (in green) is systematically more ‘noisy, i.e., less smooth, than the posterior means (orange line). However, this is not an error of the inference. Since the estimates are uncertain, the means of the posteriors will by definition have attenuated variation. If we were instead to sample *q* from its posterior distribution at each time point, we would get a curve that is similarly ‘noisy’ as the ground truth.

It is instructive to compare RealTrace’s accuracy with estimates of the volumic production rate *q* that would be obtained by simply calculate differences in total fluorescence over windows of nearby time points. As shown in Fig. 2**c**, averaging over multiple time points reduces the deviations of the estimates from the ground truth (see Supplementary Materials Sec. E.2 and Fig. S5 which shows that relative errors decrease as a power-law with window size). However, such time averaging dramatically reduces the time resolution. For example, to reach the accuracy of RealTrace’s estimates, around 10 data points are necessary, which corresponds to about half a cell cycle. Moreover, when averaging over time intervals of equal size, RealTrace is consistently at least threefold more accurate than a naive estimate. For example, for 21 measurement points, just over the mean doubling time, the relative error of RealTrace is around 3.5 % compared to 13.7 % for averaged naive estimates.

For downstream analyses of the time dynamics of cell states, it is not only important that the estimates are accurate, but also that the method provides reliable *error bars* for its estimates. Only with realistic error bars is it possible to rigorously distinguish subtle changes that are biological in origin from mere apparent differences resulting from measurement errors. We thus also tested the accuracy of the accuracy estimates that RealTrace reports. In particular, for each estimate of the volumic production *q*, RealTrace reports an error bar *ε*_*q*_. If these error bars are accurate, then the ratio (*q − q*_true_)*/ε*_*q*_ of the estimate *q* from the ground truth *q*_true_ relative to the reported error bar *ε*_*q*_, should show a standard Gaussian distribution with mean zero and variance one. Fig. 2**d** shows that the histogram of this ratio indeed very accurately follows a standard normal distribution, confirming the accuracy of the error bars that RealTrace reports. This implies that the predictions neither systematically under-nor overestimate the production rate and that the error bars correctly reflect the uncertainty of the predictions. Analogous results for the estimated growth rate are shown in Fig. S6. Finally, we confirmed that this accuracy of RealTrace not only applies to this example dataset, but that the same accuracy is observed across all 100 synthetic datasets (Fig. S7).

### 2.5 Examples of RealTrace’s application to diverse time-lapse fluorescence microscopy data from different organisms

RealTrace can be applied very widely to time-lapse fluorescence microscopy data. Although RealTrace can, of course, be applied to data from biological entities of constant, it was specifically designed for tracking of biological objects for which growth and production of fluorescence fluctuate continuously. To demonstrate RealTrace’s broad applicability, we apply RealTrace to three datasets of this type (Fig. 3): 1. lineages of *E. coli* cells growing in a microfluidic device called the Mother Machine ^26,27^ that carry transcriptional GFP reporters of both constitutive and ribosomal promoters, 2. nuclei of mouse embryonic stem cells carrying a GFP reporter of the RALY gene ^38^, and 3. entire *C. elegans* larvae expressing GFP from the constitutive *eft-3* promoter^13^.

**Figure 3:**
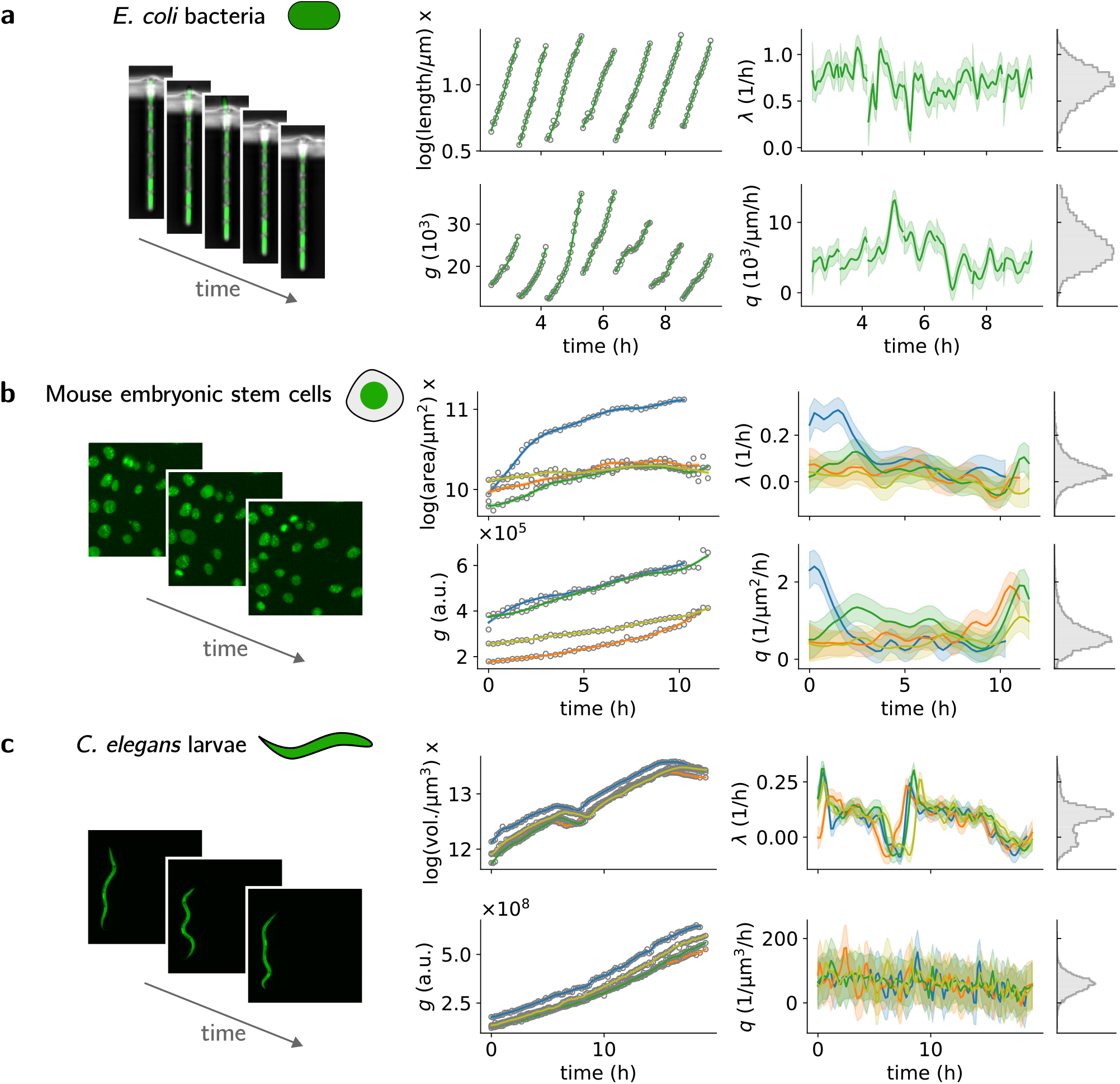
RealTrace applications to diverse time-lapse fluorescence microscopy data sets. Each row of panels shows example raw image data on the left, raw log size and fluorescence measurements (open circles) with inferred time traces (lines) in the middle, and example inferred time traces of growth rate *λ* and volumic production *q* (lines) with error bars (bands) on the right. The histograms show distributions of growth-rate and volumic production in the entire dataset. In rows **b** and **c**, the different colors refer to different stem cells and larvae, respectively. **a** One growth channel from Mother Machine containing *E. coli* cells that grow on glucose and express GFP regulated by a constitutive promoter, *hi1*. **b** Mouse embryonic stem cell nuclei with GFP reporting the expression of *RALY* (data from Ref. ^38^). **c** *C. elegans* larvae during the third and fourth larval stages expressing GFP-luciferase under the control of the ubiquitously expressed *eft-3* promoter (data from Ref. ^13^).

For each of the datasets, RealTrace uncovers novel quantitative features in the dynamics that are hidden under the measurement noise and that cannot be recovered by simple smoothing or averaging of the measurements. Below, we give an in-depth discussion of RealTrace’s findings on the *E. coli* data, and here we briefly highlight some of the observations on the mouse stem cell and C. elegans datasets.

How much does gene expression fluctuate in time in a single cell, on what time scale do these fluctuations occur, and to what extent are fluctuations in growth rate and gene expression correlated? These are fundamental questions about gene expression at the single-cell level that should, in principle, be answerable with time-lapse fluorescence microscopy data. However, the noise in the measurements makes it very challenging to obtain accurate quantitative estimates.

As can be seen in Figs. 3**b** and S8, the time traces of the measured nuclear size and total fluorescence (reporting expression of the nuclear RNA-binding protein RALY) show evidence of substantial measurement noise, i.e., although general trends are evident, the actual measured values erratically jump around this trend from one time point to the next. Consequently, just as we so for the synthetic data in the last section, naive estimates of the volumic GFP production rate *q* are extremely noisy, and averaging over time windows cannot address this issue (Fig. S9). In contrast, RealTrace recovers accurate estimates with realistic error bars. As a consequence, the true distribution of volumic production rates can only be recovered by RealTrace (Fig. S10) and this shows, for example, that fluctuations in volumic production *q* are asymmetric, with a long tail toward high production rates.

RealTrace also implements methods for rigorously calculating two-point correlations between state variables at different time points (Sec. E.8 of the Supplementary Materials). This allows us to rigorously quantify how long fluctuations in growth rate and volumic production rate last, and the correlation between these rates (Fig. S11). We find that volumic production fluctuations last almost twice as long as growth rate fluctuations, i.e. correlation times of 0.9 and 0.55 hours respectively (Fig. S11**a** and **b**). Notably, these correlation times are short relative to the cell cycle time, which is on the order of 12 hours for these mouse embryonic stem cells, suggesting that these fluctuations are not related to the cell cycle. Strikingly, growth rate and volumic production rate are positively correlated over a time scale of about 2 hours, with a clear peak in correlation at the same time point (Fig. S11**c**), suggesting that these fluctuations might result from the same underlying cause. More discussion of the results on the mouse embryonic stem cell data is in section B of the Supplementary Materials.

For the time traces of the *C. elegans* larvae, RealTrace infers that the volumic production from the constitutive *eft-3* promoter is fairly constant across time and larvae (Fig. 3**c**, bottom right). In contrast, the dynamics of growth rate shows a complex pattern that is quite reproducible across larvae and includes an event where the larvae’s size shrinks and the growth rate becomes negative (Fig. 3**c**, top right). This dramatic event, which occurs at the 4th moulting, is also reflected in a bimodal distribution of growth rates overall. RealTrace’s accurate estimates of the growth rates across time in individual larvae allow for precise quantification of the variation in this moulting event across larvae, which cannot be obtained by simple time averaging (Figs. S12 and S13). Further discussion of the results on the *C. elegans* data are in section C of the Supplementary Materials.

In summary, these results show that RealTrace can be applied to time-lapse microscopy data from a broad range of biological systems, and that it is able to recover subtle quantitative features of the biological dynamics, including the dynamics on short time scales and their correlation structure, that would otherwise be invisible. To further demonstrate RealTrace’s ability to uncover novel dynamical features, we next present a more in-depth study of the *E. coli* dataset.

### 2.6 Growth rate fluctuations increase in amplitude and duration at slow growth

For single-celled organisms such as bacteria, growth rate is arguably one of the most important phenotypes determining evolutionary success. It has been well appreciated that, at the single-cell level, even genetically identical cells can exhibit substantial growth rate fluctuations. Moreover, since fast-growing cells naturally outcompete slow-growing cells, the growth rate at the population level is a complex function of the distribution of single-cell growth rates ^39^, and this can be exploited in so called bet-hedging strategies ^40,41^.

Beyond its crucial role for fitness, the fluctuations in growth rate across time in single cells are also interesting because they reflect fluctuations in the rate of biomass production by the biochemical reaction network inside the cell, and thus provide quantitative information about how intra- and extra-cellular noise affects the overall functioning of this reaction network.

We obtained datasets of *E. coli* cells growing in minimal growth media with either acetate, glycerol, or glucose as a carbon source. For each condition, we used RealTrace’s results to calculate both the average of the instantaneous growth rate and quantified its variability by its coefficient of variation (i.e., the standard deviation divided by mean, see Sec. E.9 of the Supplementary Materials for details). We find that the growth rate fluctuations are substantial, i.e., with CV ranging from 30% in glucose to 55% in acetate, meaning that instantaneous growth rates can easily vary by more than two-fold across cells (Fig. 4**a**). In addition, the growth rate fluctuations substantially increase as the average growth rate decreases, i.e., growth rate fluctuations in acetate are almost twice as large as in glucose.

**Figure 4:**
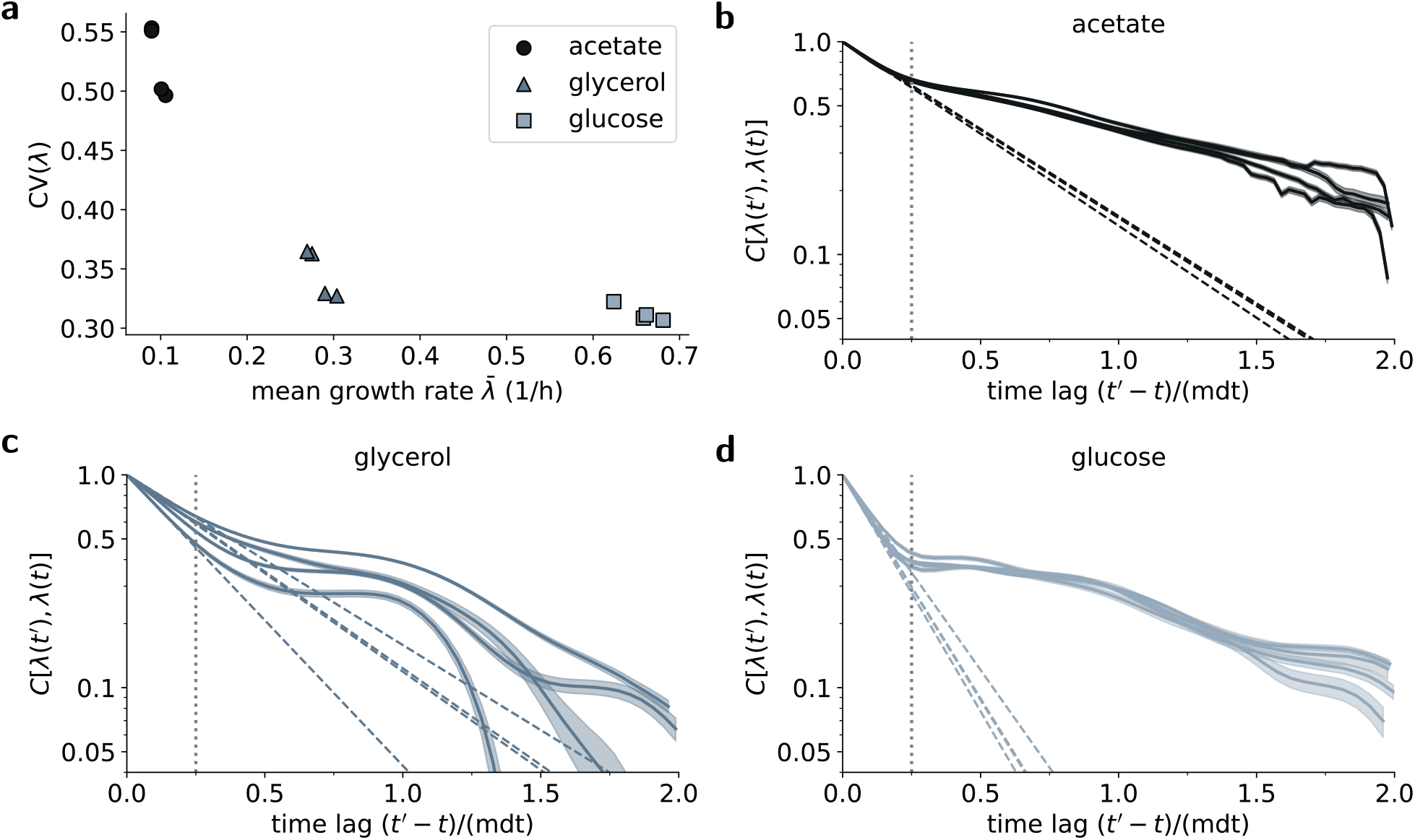
Growth rate fluctuations systematically decrease with mean growth rate across conditions. **a** The extent of growth rate fluctuations is quantified by the coefficient of variation (CV) and plotted as a function of the mean growth rate across conditions. Each symbol corresponds to one replicate experiment. **b-d** The auto-correlation function of the growth rate in each condition. Each ribbon shows the posterior mean auto-correlation and its error bars for one replicate experiment. The dashed lines show the exponentials 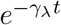 with *γ*_*λ*_ the fitted decay rate of growth-rate fluctuations in the corresponding experiment. In each condition, the auto-correlation follows a simple exponential decay with rate *γ*_*λ*_ until approximately 0.25 of the mean doubling time (vertical dotted lines).

Beyond the amplitude of the growth rate fluctuations, we also quantified how these fluctuations decay, i.e. how quickly cells revert to the mean growth rate, by calculating auto-correlation function of the growth rate (see Supplementary Materials Sec. E.10 for details). The results show that growth rate fluctuations decay faster in fast growth conditions, even relative to the mean doubling time in the respective condition (Fig. 4**b**-**d**). In particular, in each condition, the correlation functions exhibit two regimes: an initial fast exponential decay followed by a period with much slower decay. Interestingly, in each condition, the switch from the fast to slower decay occurs at approximately 1/4 of the mean doubling time. Because the initial exponential decay is steepest in glucose and least steep in acetate, much larger correlations persist in acetate than in glucose. In particular, in acetate, there are still substantial correlations in growth rate after two doublings. Thus, cell growth fluctuations in slow growth conditions are not only larger but also last for more doublings. This implies that the difference between the average single-cell growth rate and the population growth rate is largest in acetate^39^. Finally, we note that the shapes of the empirically determined auto-correlation functions (Fig. 4**b**-**d**) are very different from the simple exponential decay that our process prior assumes. This underscores that the only role of the prior is to implement the expected smoothness of the dynamics and that the inferred can differ greatly from that assumed by the prior.

### 2.7 Growth and volumic production systematically vary across the cell cycle

It is often assumed that when bacterial cells are growing exponentially that they are in so-called ‘balanced growth’ where the concentrations of all their components are constant over time. However, we know this cannot be completely the case. For example, as the DNA is replicated, the relative frequencies of different genomic loci must change in time. Indeed, recent single-cell RNA-seq analysis has uncovered a systematic dependence of mRNA levels on the stage of chromosome replication ^42^. We thus asked whether, within the substantial variation across single-cells, there is also a systematic dependence of growth rate and volumic production on the cell cycle stage.

In particular, the cell division cycle in *E. coli* is well described by an adder model in which cells add a certain volume during their cell cycle, irrespective of their birth size or growth rate ^8–10^. The added length since birth is thus a natural parametrization of the cell cycle stage. The distribution of added cell length during the cell cycle is remarkably reproducible between data sets of the same condition and shifts towards increased added length and thus larger cell sizes in the faster growth condition (Fig. S14). We quantify the cell cycle stage of a single cell at a given time point as the added length since birth normalized by the total added length in that cell’s cell cycle, thereby aligning all cell cycles on the unit interval (see Supplementary Materials Sec. E.3 for details). Even though the instantaneous growth and volumic production rates fluctuate strongly across cells (see Figs. S15-S17), averaging over many cell traces reveals systematic patterns along the cell cycle.

Strikingly, we find that in all conditions, cells grow faster towards the end of the cell cycle, following a very similar pattern across all three growth conditions (Fig. 5**a**-**c**). Thus, cells grow faster than exponentially within the cell cycle. Like the growth rate fluctuations across cells, the amplitude of this cell cycle variation is largest in the slow growth condition, i.e., around 20 % of the mean growth rate in acetatete and around 10 % in glucose. Moreover, neither the minimum nor the maximum growth rate is at the start or end of the cell cycle. In acetate, the minimum is around 1/4 of the cell cycle, and the maximum is just before the next division. In faster growth conditions, both extrema occur earlier in the cell cycle. Recent work suggests that deviations from exponential growth within single cell cycles are common across species ^43–45^.

**Figure 5:**
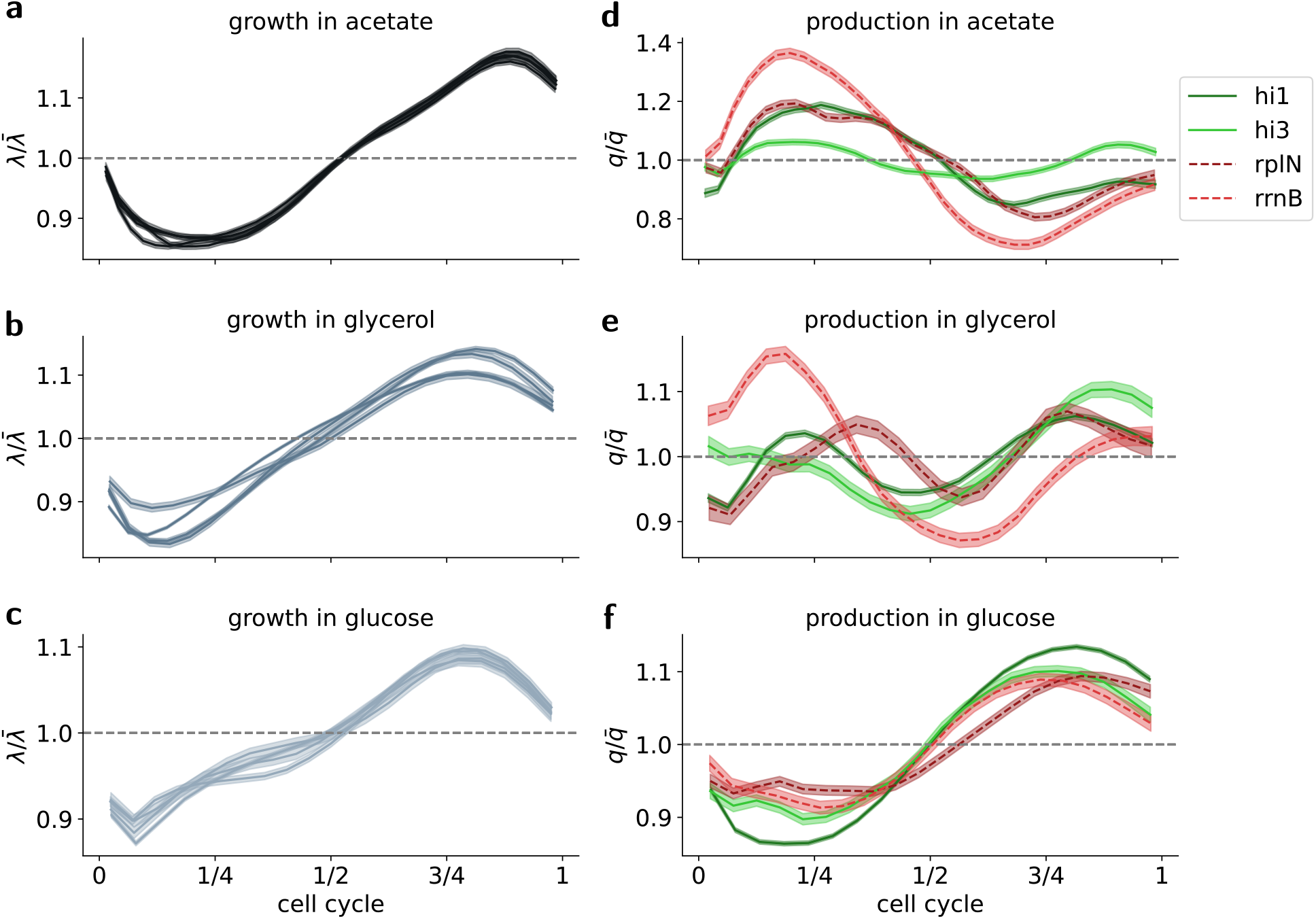
Growth and volumic production rate vary systematically across the cell cycles. **a** Average growth rate as a function of cell cycle stage for cells growing on acetate. The cell cycle is parameterized by the added length since birth, normalized by the total length added during the cell cycle. Growth rate is normalized by the mean growth rate across all cells and time. Each ribbon corresponds to mean plus and minus one standard error in one replicate experiment. **b** and **c** As in panel **a**, but with cells growing on glycerol and on glucose, respectively. **d** Volumic GFP production rate normalized by the respective mean production rate as a function of cell cycle stage. The four data sets correspond to four different promoters that regulate the expression of GFP (see legend) with synthetic constitutive promoters in green and ribosomal promoters in red. Lines and ribbons correspond to mean plus and minus one standard error. **e** and **f** As in **d** but from cells growing in glycerol **f** and glucose, respectively.

We similarly measured systematic variation in volumic production rate across the cell cycle for two synthetic constitutive promoters (green lines in panels **d**-**f** of Fig. 5) and two ribosomal promoters (red lines in panels **d**-**f** of Fig. 5). Like for growth rate, we find that the amplitude of the systematic variations in volumic production are largest in slow growth and smallest in fast growth, e.g. the ribosomal RNA promoter *rrnB* deviates up to around 40 % from the mean production rate in acetate. Notably, however, the fluctuations across cells are still larger than the variation across the cell cycle (Fig. S15).

Surprisingly, in contrast to the cell cycle dependence of growth rate, the patterns of cell cycle dependence in volumic production rate vary strongly across conditions, and even across promoters in the same condition. In particular, whereas in the fast growth condition (glucose) the volumic production rate increases along the cell cycle just as the growth rate, in the slow growth condition (acetate) the pattern is the opposite with the highest volumic production early in the cell cycle. At an intermediate growth rate in glycerol, the patterns are more complex and show more variation across promoters, even between constitutive promoters. Notably, all these systematic cell cycle dependencies are similar if the cell cycle stage is not defined by added length since birth but by time since birth (Fig. S18).

The cell cycle dependencies show that the physiology of exponentially growing bacteria varies systematically across the cell cycle. The fact that the growth rate shows very similar cell-cycle patterns in all conditions suggest that these might be driven by a common mechanism. In contrast, the volumic production rates show patterns that vary across conditions and across promoters. It is striking that the patterns in volumic production across the cell cycle are almost reversed between fast and slow growth, and that they are more complex and varied across promoters at intermediate growth. One possible interpretation is that there are multiple mechanisms affecting volumic production and that one mechanism dominates in slow growth, another dominates in fast growth, and that at an intermediate growth rate more complex patterns emerge because different mechanisms have similar strength and different promoters are affected to different extents by each mechanism. Notably, even different constitutive promoters can exhibit different cell cycle dependencies and the volumic production from ribosomal promoters is not directly tied to variations in growth rate.

It is currently unclear what the mechanisms underlying these complex cell cycle dependencies might be. They could reflect not only variation in transcription and mRNA decay rates, but also variation in ribosome concentrations and translation elongation rates. Whatever these mechanisms are, their strength must systematically vary across growth conditions. In addition, when different promoters exhibit different patterns in the same condition, these differences must reflect a different pattern in transcription since these transcriptional reporters only vary in their promoter sequence.

### 2.8 Inferring responses to switching conditions

To uncover the principles of gene regulation at the single-cell level, it may be especially informative to observe how cells adapt to a sudden switch in their growth environment. Not only the average growth rate but also the rates of production of all proteins will transition from a previous ‘steady state’ to a new one and it is interesting to quantify the transient dynamics of growth rate and volumic production and how they vary across single cells.

To enable rigorous analysis of datasets with environments that are not constant, we extended RealTrace so that not only the parameters of the prior process, but also the experimental parameters such as exposure time and acquisition frequency may be different for different time segments of the datasets. RealTrace infers separate sets of optimal parameters Θ^*∗*^ for each time segment and posterior distributions for the states of the cells are calculated using the same iterative procedure as described before, but using different parameters Θ^*∗*^ for different time points.

To demonstrate RealTrace’s ability to analyze datasets with changing conditions, we applied it to a dataset in which *E. coli* cells grow in the mother machine and their growth media are switched from minimal media with glucose to minimal media with glycerol (a poorer nutrient). An example single-cell lineage shows that both the growth rate and volumic production rate of the constitutive promoter *hi1* decrease in glycerol and that growth is almost completely arrested directly after the switch to glycerol (Fig. 6**a**).

**Figure 6:**
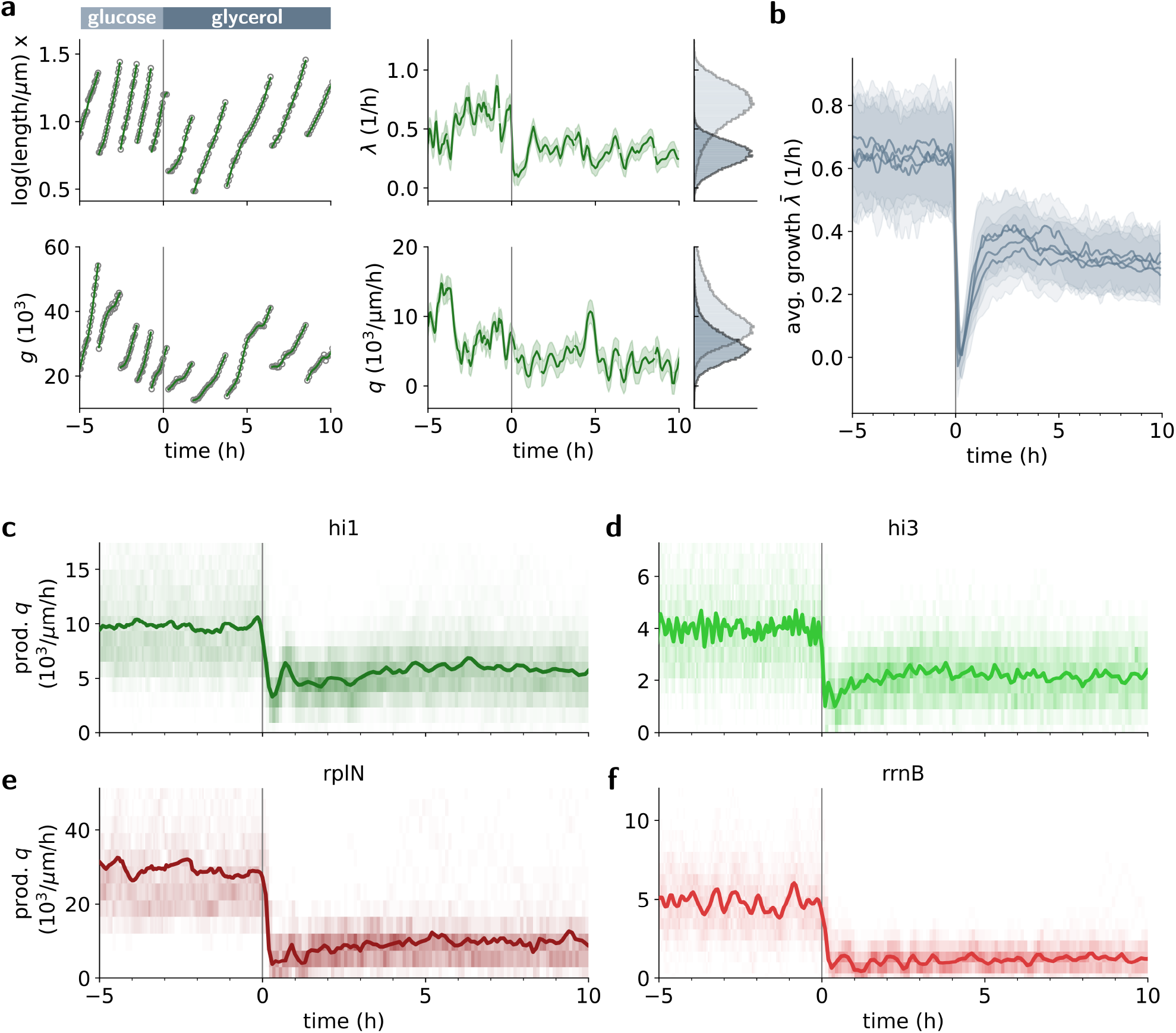
Response of single *E. coli* cells to a switch in nutrients. **a** Example of size (top left), growth rate (top right), total fluorescence (bottom left) and volumic production rate (bottom right) for a single cell lineage carrying a transcriptional reporter of the constitutive promoter *hi1* as growth media switch from glucose to glycerol 0 h (vertical line). The dots correspond to measurements and the curves with ribbons to the means and standard deviations of the posteriors inferred by RealTrace. The histograms on the right show the distributions of growth rate and volumic production in glucose (light) and glycerol (dark). **b** Average growth rate as a function of time across 4 replicate experiments. The lines and shaded areas show means and standard deviations across the population for each replicate. **c**-**f** Average volumc production as a function of time for the promoters *hi1, hi3, rplN*, and *rrnB*, respectively. The solid lines show population averages and the shaded areas show histograms of the distribution across the population at each time point.

In a previous study, we observed that, upon a switch in nutrients, different *E. coli* cells may respond on very different time scales, so that there is no clear average behavior as a function of time ^46^. We also observed highly variable responses across cells to the addition of a DNA-damaging agent to the growth medium ^47^. In contrast, we recently found that upon loss of carbon source, the time-dependent responses are highly reproducible across single cells ^48^.

Here we find that, upon a switch in nutrients from glucose to glycerol, the single cells show very consistent time-dependent responses. Upon the switch, all cells show an immediate drop in growth rate, coming almost to growth arrest, which is then followed by an overshoot in growth rate, peaking at about 2 hours, which subsequently takes 6-8 hours (3-4 cell cycles) to fully settle down. Notably, this dynamics of growth rate is not only highly consistent across single cells, but also across multiple replicate experiments. In contrast, while the volumic production rates of all 4 promoters also drop sharply immediately after the switch, all promoters almost immediately take on their new steady-state level, with only a mild further adaptation in the first 3 hours for *hi1, hi3*, and *rplN*.

It is striking that the growth rate undergoes a so much longer transient than the volumic production rates. It is tempting to speculate that the time scale for the growth rate to reach a new steady-state corresponds to the time scale it takes to replace the cell’s proteome with a new one, i.e. it takes 3-4 cell cycles for the old proteome to be diluted out. It is puzzling, however, how the different promoters ‘know’ so quickly what their new steady-state volumic production rates should be. This suggests that these new steady-state production rates might be set not by changes in the proteome such as concentrations of ribosomes and transcription factors.

## 3 Discussion

By enabling quantitative measurements of the behavior of single cells or organisms over time, time-lapse fluorescence microscopy provides one of the most powerful approaches toward unraveling the mechanisms underlying complex biological behaviors. However, because the true states of biological systems change smoothly in time, much of the dynamics on short time scales is hidden under unavoidable measurement noise.

Here we have presented a new method, called RealTrace, that leverages the fact that while biological changes are correlated on short time scales, measurement errors are all independent, to rigorously disentangle measurement noise from true biological fluctuations. Importantly, by using maximum entropy process priors, RealTrace is able to infer the true underlying biological dynamics without having to make specific assumptions about the form that the dynamics might take.

Using realistic synthetic data, we showed that RealTrace not only accurately estimates the true biological dynamics without bias, but also provides accurate error bars for its estimate. This is especially important for downstream analyses, because it allows to rigorously distinguish subtle biological differences from merely apparent differences caused by measurement errors. Accurate error bars are also important to assess whether the data are consistent with mathematical models of the behavior of the system. Beyond estimates of instantaneous cell states, RealTrace also rigorously calculates correlation functions between observables at different time points, allowing for accurate quantification of the time dynamics of fluctuations.

Our applications to real datasets show that, in each case, RealTrace uncovers subtle features of the dynamics that would otherwise be hidden below measurement noise. In particular, our in-depth analysis of the single-cell growth and expression dynamics of *E. coli* cells growing in different environments uncovered a host of novel dynamical features that are not explained by current models of growth and gene expression dynamics in bacteria. This includes the observation that single-cell fluctuations in growth rate become large and last longer in slower growth conditions, and that the growth rate varies systematically across the cell cycle in a way that is independent of growth condition, but that volumic production rates vary across the cell cycle in a way that depends both on the condition and on the promoter. The fact that even different constitutive promoters, that are not thought to contain any regulatory sites beyond the polymerase binding site, show different patterns of cell cycle dependence in production is especially challenging to explain, since these differences must stem from differences in transcription (as the reporters are otherwise identical). Finally, upon a switch in nutrients, we found that volumic production rates from different promoters quickly reach a new steady-state, but that growth rate shows a much longer transient dynamics with an initial almost complete growth arrest, followed by an overshoot that takes several cell cycles to settle down. All these observations are new insights into the physiology and gene expression of bacteria at the single-cell level, which hint at underlying mechanisms that are not yet incorporated in current models.

We envisage that RealTrace can similarly provide important new insights in a very wide range of systems. RealTrace applies to essentially any kind of time-lapse fluorescence microscopy data. To demonstrate this broad applicability, we analyzed time-lapse fluorescence microscopy data from three very different biological systems: *E. coli* cells growing in a microfluidic device, nuclei of mouse embryonic stem cells and whole *C. elegans* larvae. Moreover, RealTrace has been designed to analyze datasets in which conditions change during the experiment. RealTrace can of course also be applied to data where cells are growth-arrested, e.g. we recently analyzed the gene expression responses of *E. coli* cells upon the loss of nutrients^48^. Importantly, beyond application to fluorescent reporters of gene expression, RealTrace is also directly applicable to data with fluorescent reporters of other biological activities, e.g. fluorescent biosensors that visualize intracellular molecules such as the abundance of calcium or antibiotic compounds.

## 4 Methods

### 4.1 Decomposition of the posterior distributions and the likelihood

We seek to calculate posterior distributions for the true state at each measurement point, which includes the true log size, total GFP, growth rate, and volumic production rate. To estimate the parameters of the prior process and the measurement noise, we also calculate the total likelihood of the data, which we then numerically optimize with respect to the parameters. We then use these optimal parameters to calculate and report the full posterior distribution of the cell states at each time point. We can exploit the Markovian property of the prior process to factorize the likelihood and decompose the full posterior into a forward and a backward part, i.e. the probability of the current state of the cell given all data in the past and all data in the future, respectively. Then, using an iterative procedure similar to the forward/backward algorithm used in hidden Markov model analysis, we can determine the contributions of individual time points to the total likelihood and the forward and backward parts of each time point.

First, the likelihood of the entire dataset consisting of *N* measurements *D* = {**y**_1_, …, **y**_*N*_} can be factorized into likelihoods of measurements at individual time points **y**_*n*_ conditioned on all previous measurements

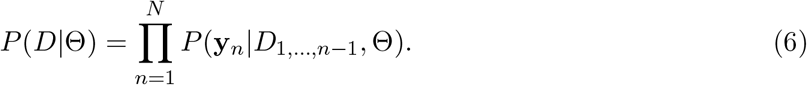

Second, as shown in Supplement section E.4, the full posterior for the state *z*_*n*_ at time point *n* conditioned on all measurements *D* can be decomposed into a forward and a backward part as follows

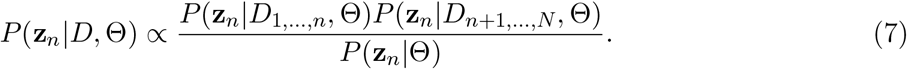

Here *P* (**z**_*n*_|*D*_1,…,*n*_, Θ) is the forward part incorporating all measurements up to the time point *n* and *P* (**z**_*n*_|*D*_*n*+1,…,*N*_, Θ) is the backward part conditioned on the remaining measurements in the future of time point *n*. The prior distribution *P* (**z**_*n*_|Θ) in Eq. (7) is uniform over *x*_*n*_ and *g*_*n*_ and the priors for the growth rate *λ*_*n*_ and volumic production rate *q*_*n*_ are given by the stationary distribution of the maximum entropy process prior, which are equivalent to Ornstein-Uhlenbeck processes.

In the next sections, we will outline the calculation of the forward part *P* (**z**_*n*_|*D*_1,…,*n*_, Θ). To calculate the backward part of the posterior *P* (**z**_*n*_|*D*_*n*+1,…,*N*_, Θ), we use the same scheme as for the calculation of the forward part but with time-reversed differential equations and reversed equations for cell divisions (see Sec. E.7 of the Supplementary Materials for details).

### 4.2 Calculation of the prior distribution

At each step of the iterative procedure, we start from the current posterior distribution *P* (**z**_*n*_|*D*_1,…,*n*_, Θ) for the state of the cell at time point *n* given all measurements up to this time point, and then use the prior process to calculate a prior distribution *P* (**z**_*n*+1_|*D*_1,…,*n*_, Θ) for the next time point *n* + 1. The prior process provides the conditional distribution *P* (**z**_*n*+1_|**z**_*n*_, Θ), i.e. the probability of the state of the cell at time point *n* + 1 given a state of the cell at time point *n*. The conditional distribution only depends on the parameters of the prior process and not on the measurement noise parameters. To make the calculations trackable, we approximate the conditional distribution *P* (**z**_*n*+1_|**z**_*n*_, Θ) as a multivariate Gaussian. Consequently, we only need to calculate the first two moments of the distribution. Second, to cope with the non-linear coupling in the GFP dynamics, we ignore the effect of growth rate fluctuations on the GFP production between time points,

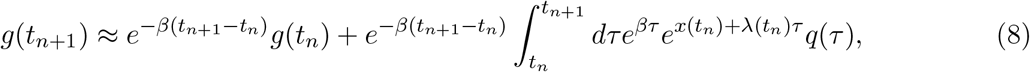

where we approximated *x*(*τ*) *≈ x*(*t*_*n*_) + *λ*(*t*_*n*_)*τ* inside the integral. As long as measurements are taken sufficiently frequently, this approximation will be very accurate, i.e. as long as the growth rate changes little between consecutive time points. We then calculate expectation values for the second moments of the Gaussian *P* (**z**_*n*+1_|*D*_1,…,*n*_, Θ) by integrating over **z**_*n*_ weighted by the posterior *P* (**z**_*n*_|*D*_1,…,*n*_, Θ) using Wick’s probability theorem. In this way, we obtain the new prior distribution *P* (**z**_*n*+1_|*D*_1,…,*n*_, Θ) (see also Fig. 1). Using this new prior, we calculate the likelihood and the forward part of the posterior of the next time point. A more detailed derivation is given in Sec. E.7 of the Supplementary Materials.

### 4.3 Calculation of the posterior and the likelihood

Given the prior distribution *P* (**z**_*n*+1_|*D*_1,…,*n*_, Θ), we calculate the likelihood of the measurement and the forward part of the posterior. The likelihood of the single measurement **y**_*n*+1_ is given by,

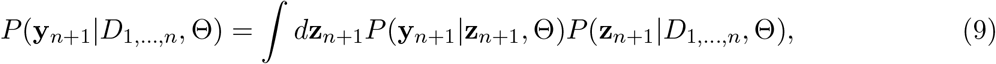

Since we assume Gaussian measurement noise, the probability of obtaining the measurement **y**_*n*+1_ given the true state of the cell (*x*_*n*+1_, *g*_*n*+1_) reads,

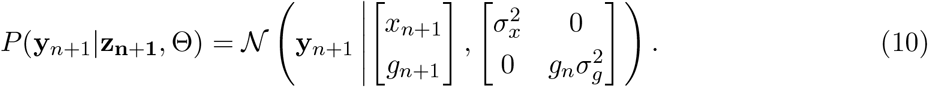

This probability distribution only depends on the measurement noise parameters, but not the parameters of the prior process. Note that, in this way, we obtain likelihoods for each measurement, which together determine the total likelihood as given by equation (6).

To obtain the forward part of the posterior for the time point *n* + 1, we incorporate the measurement using

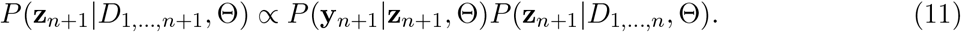

The explicit expressions for the likelihood and the posterior are shown in the Supplementary Materials (Sec. E.5).

### 4.4 Joint posterior distributions for two time points

The calculation of correlation functions includes statistics of the state of the cell at two time points. Since two posterior distributions along a lineage are not independent, we thus need to calculate the joint posterior distributions of two time points. To calculate joint posterior distributions of two time points **z**_*n*+*m*_ and **z**_*n*_ that are conditioned on all measurements, we again formulate an iterative procedure. In short, we calculate the prior distribution *P* (**z**_*n*+1_, **z**_*n*_|*D*_1,…,*n*_, Θ) given the posterior distribution for **z**_*n*_, similar to the prior distributions for single time points. This allows us to calculate the forward part of the posterior *P* (**z**_*n*+1_, **z**_*n*_|*D*_1,…,*n*+1_, Θ). To iterate this procedure, we integrate the distribution *P* (**z**_*n*+2_, **z**_*n*+1_, **z**_*n*_|*D*_1,…,*n*+1_) = *P* (**z**_*n*+2_|**z**_*n*+1_, *D*_1,…,*n*+1_)*P* (**z**_*n*+1_, **z**_*n*_|*D*_1,…,*n*+1_) over the intermediate time point **z**_*n*+1_ to obtain the new prior distribution. In this way, we calculate all distributions *P* (**z**_*n*+*m*_, **z**_*n*_|*D*_1,…,*n*+*m*_, Θ), which are the forward parts of the two-time-point posteriors. Importantly, this posterior incorporates the intermediate measurements **y**_*n*+1_, …, **y**_*n*+*m*_. Then, similar to the single time point posteriors before, the posterior distribution that is conditioned on all measurements reads,

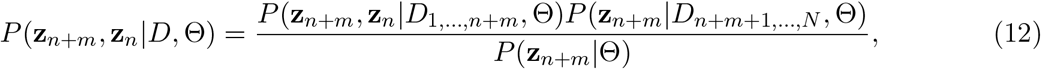

where the prior *P* (**z**_*n*+*m*_|Θ) is chosen as before. A detailed derivation for the joint posterior can be found in the Supplementary Materials in Sec. E.8, and the calculation for the auto-correlation function is given in Sec. E.10.

### 4.5 Experimental data

We analyzed time traces of 90 single *C. elegans* larvae ^13^. The *C. elegans* larvae ubiquitously express GFP from the *eft-3* promoter. The size measurements correspond to the measured volume of the single larvae at each time point.

The mouse embryonic stem cell data set consists of 170 mouse embryonic stem cell nuclei traces published in Ref. ^38^. Here, the size corresponds to the area of the nucleus of the cells and the fluorescence tracks the expression of the *RALY* gene, which encodes the nuclear RNA-binding protein Raly.

The experiments with *E. coli* cells closely follow the experimental procedures previously described in Ref. ^46^. We used 4 strains that each have promoter-GFPmut2 cassettes integrated at the HKO22 locus of the chromosome of the *E. coli* K-12 MG1655 strain. The promoters used included two synthetic constitutive promoters called *hi1* and *hi3*, which each express at a level similar to native ribosomal promoters. In addition, we used the promoter of the ribosomal protein *rplN*, The synthetic promoter sequences were taken from Ref. ^49^ and all promoter sequences are provided in the Supplementary Materials Sec. F.

Cells were grown in the dual-input Mother Machine ^27^ using M9 minimal media supplemented with 0.05 % acetate, 0.4 % glycerol or 0.2 % glucose at 37 ^*°*^C. Images were acquired every 12 min, 6 min, or 3 min for cells grown on acetate, glycerol, and glucose, respectively. The microscopy data were analyzed with a new version of the image analysis software MoMA introduced in Ref.^27^. The new version, called DeepMoMA, uses a trained version of U-Net to segment the cells.

### 4.6 Code availablity

RealTrace is implemented in `C++`,

The code is freely available from https://github.com/nimwegenLab/RealTrace.

## Acknowledgements

We would like to thank Daan de Groot and Théo Gervais for many discussions and feedback. This research was supported by the Swiss National Science Foundation’s grants numbers 159673 and 184937 to Erik van Nimwegen, and Sinergia grant number 189910 to David Suter and Erik van Nimwegen.

## Supplementary Materials

### A Figures with results on the synthetic data

**Figure S1:**
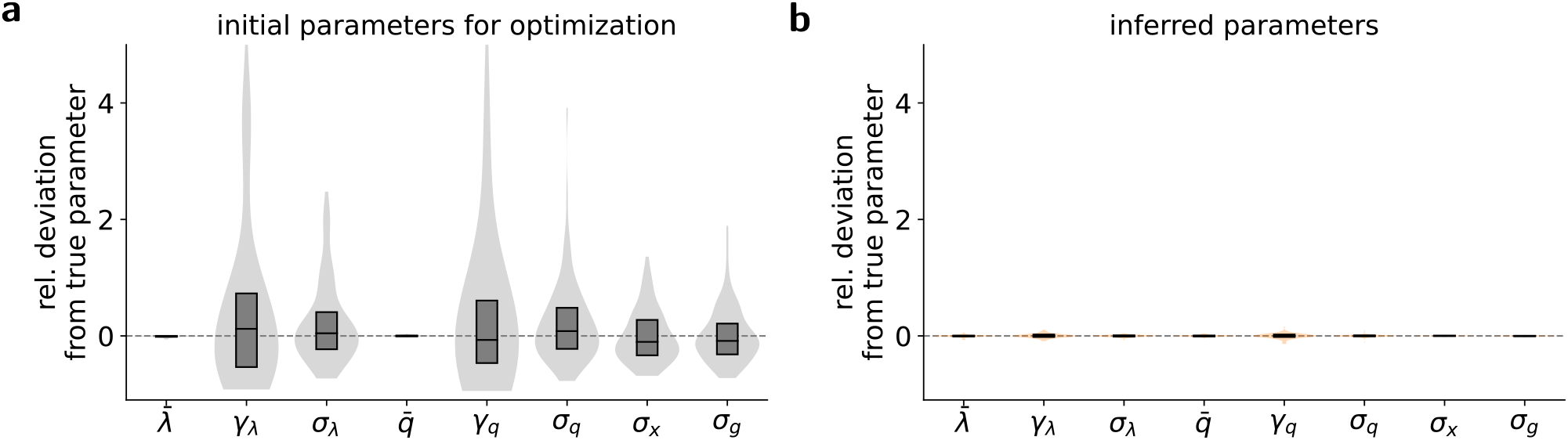
**a** The numerical optimization of the likelihood is initialized with random parameters (see Sec. E.1 for details) apart from the means, 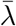 and 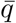, which can be directly estimated from the noisy input traces. The violin plots and box plots (median and inter-quartile range) summarize the distributions of the relative deviations of the initial parameters from the true parameters. **b** Violin plots and box plots of the inferred parameters, replotted from Fig. 2**b**.

**Figure S2:**
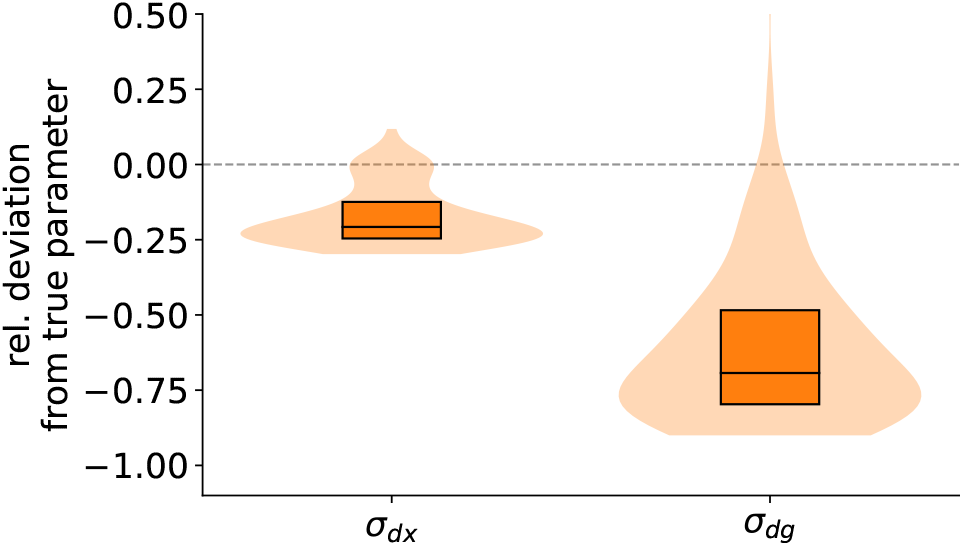
When a single cell at time point *n* has divided into two cells at time point *n* + 1, the division could in principle have occurred at any time *t* between *t*_*n*_ and *t*_*n*+1_. However, marginalizing over this unknown time *t* would make the inference procedure much more complex. We thus chose to make the approximation that we assume each division occurs exactly at the time point where two cells are first observed. As shown in this figure, this approximation results in a systematic underestimation of the parameters for cell divisions. However, we have seen that the posterior distributions for cell states are extremely precise despite this simplification. Thus, this approximation does not compromise the cell dynamics estimates of RealTrace.

**Figure S3:**
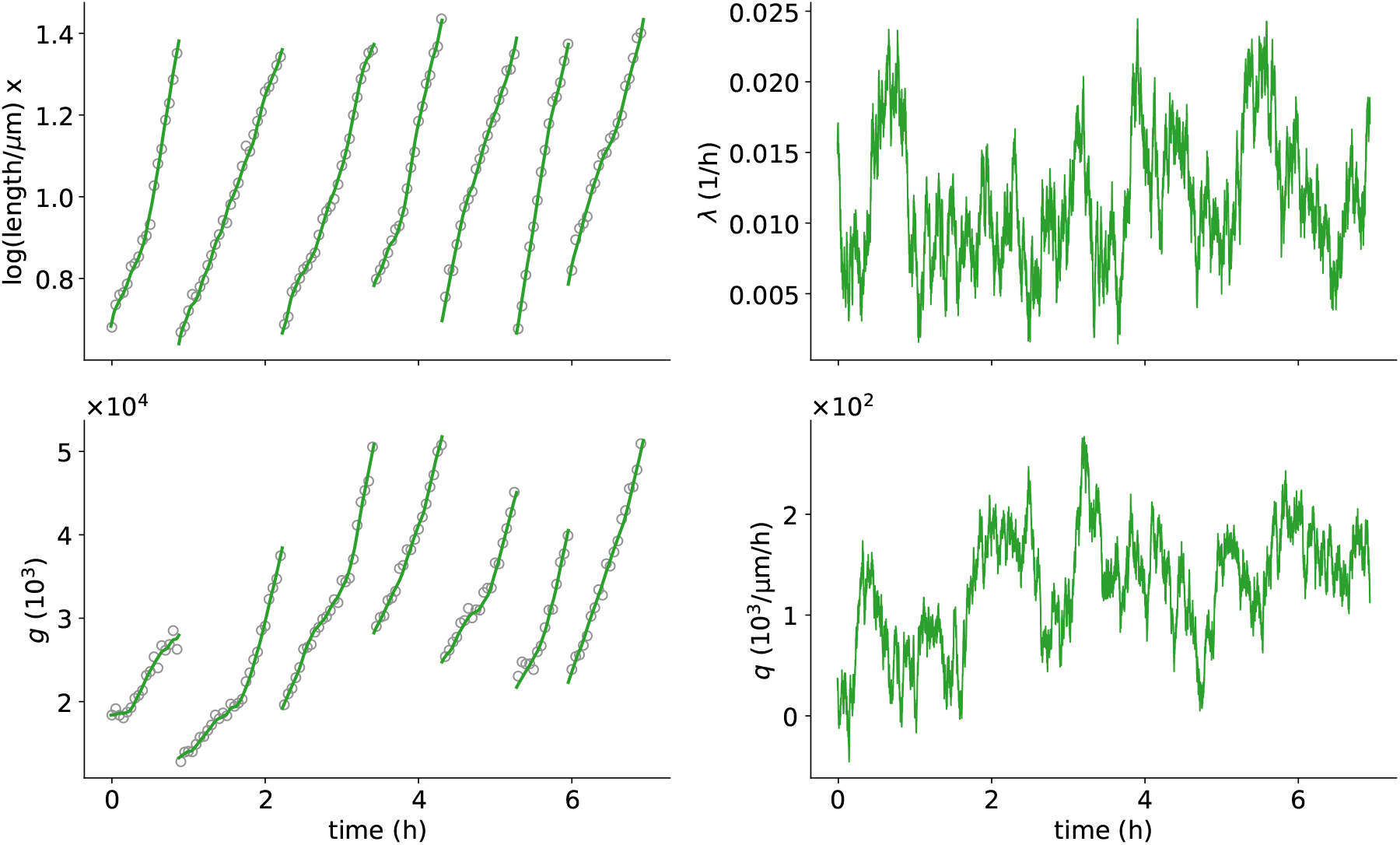
Cell dynamics of an example lineage of the synthetic data set shown in Fig. 2. The true process shown in green follows the model formulated to generate prior distributions. Then, we add Gaussian noise to the log length and the total GFP content to mimic noisy measurements (grey dots). The noisy measurements are the sole input used by RealTrace to infer the true underlying process. Details of the simulation are given in Sec. E.1 of the Supplementary Methods, and the parameters of the simulation are given in Table 1.

**Figure S4:**
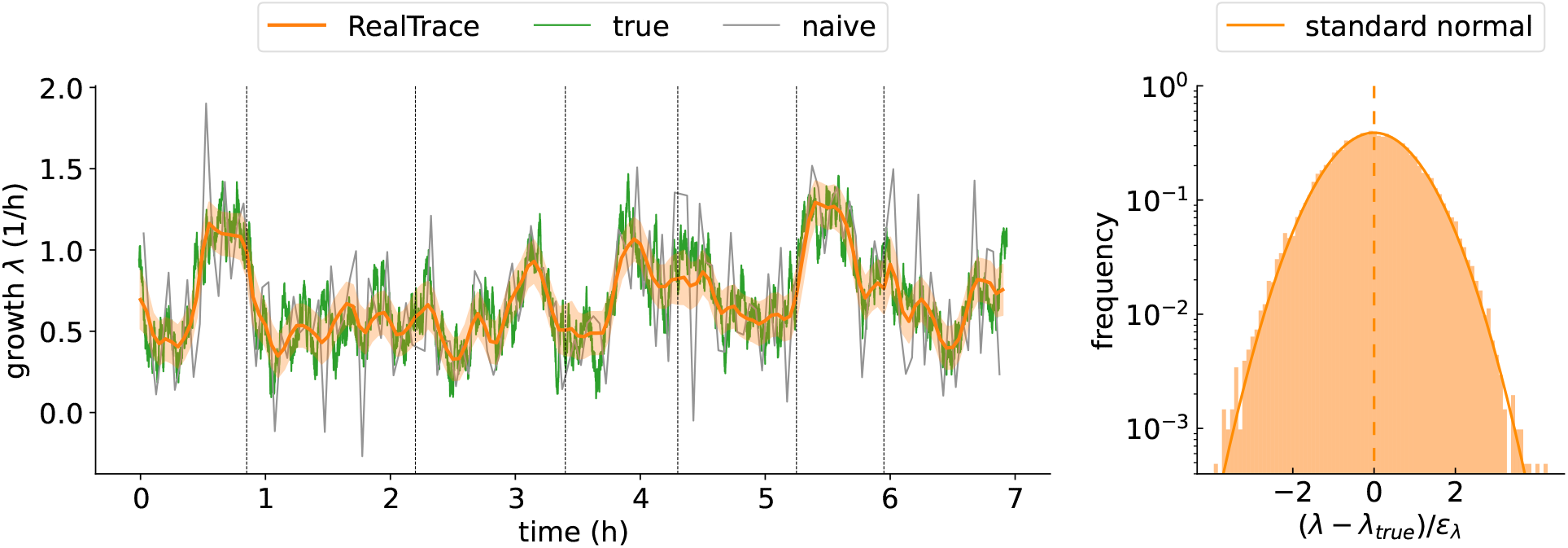
Left panel: Comparison of the true growth rate (green), the growth rate inferred by RealTrace (orange), and the naive estimate of growth rate from finite differences in log-size (grey). The orange curve and ribbons correspond to the mean and plus/minus one standard deviation of RealTrace’s posterior. The growth rate estimates by RealTrace closely follow the ground truth. Right panel: The histogram of the deviation of the mean of the growth rate posterior from the ground truth, divided by the standard deviation of the posterior *ε*_*λ*_ is very close to a standard normal. Thus, RealTrace neither systematically over-nor underestimates the growth rate, and the error bars (standard deviation of the posterior) correctly reflect the uncertainty of the estimates. These results are analogous to the results shown in Fig. 2**b** and **d** for the volumic production rate instead of growth rate.

**Figure S5:**
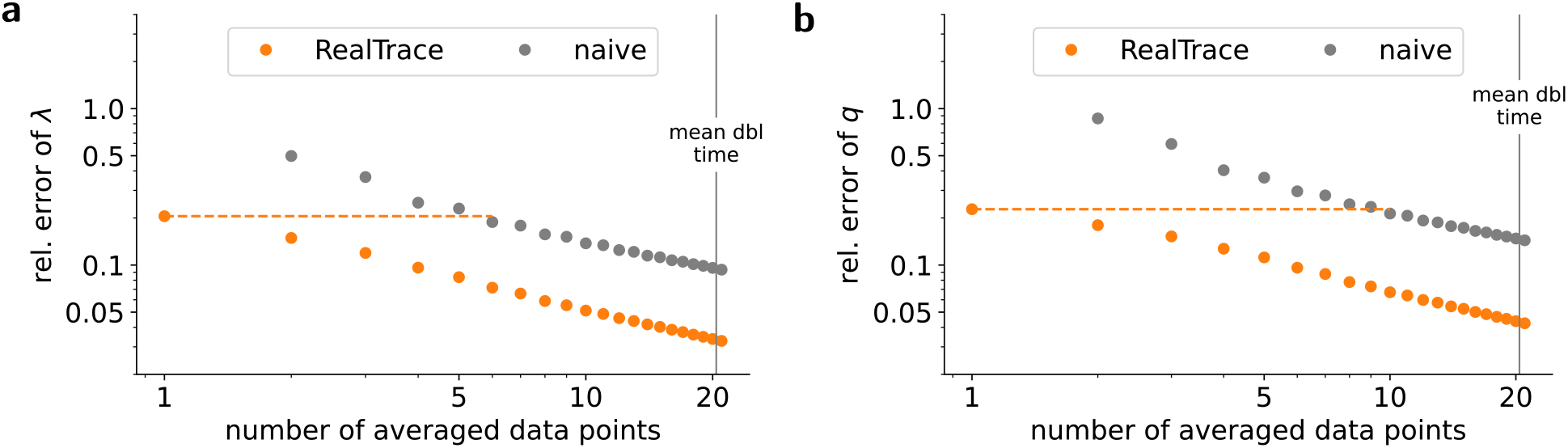
Relative root mean squared deviations of **a** the growth rate and **b** the production rate. The results are shown on a log-log scale to highlight that the relative error continues to decrease the more data points are included in the average.

**Figure S6:**
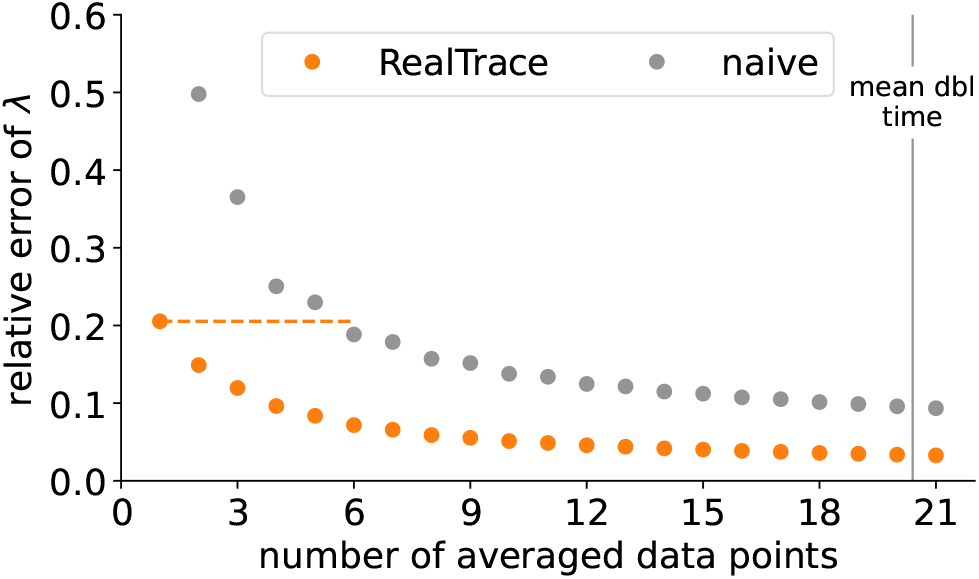
Relative root mean squared deviations between the estimated average growth rate and true average growth rate over time windows of data points of different lengths for both RealTrace (orange) and naive time averages (grey). The mound doubling time is indicated as a vertical line. Note that this plot is analogous to Fig. 2**c** of the main text, which shows the same statistics for volumic production rates instead of growth rates.

**Figure S7:**
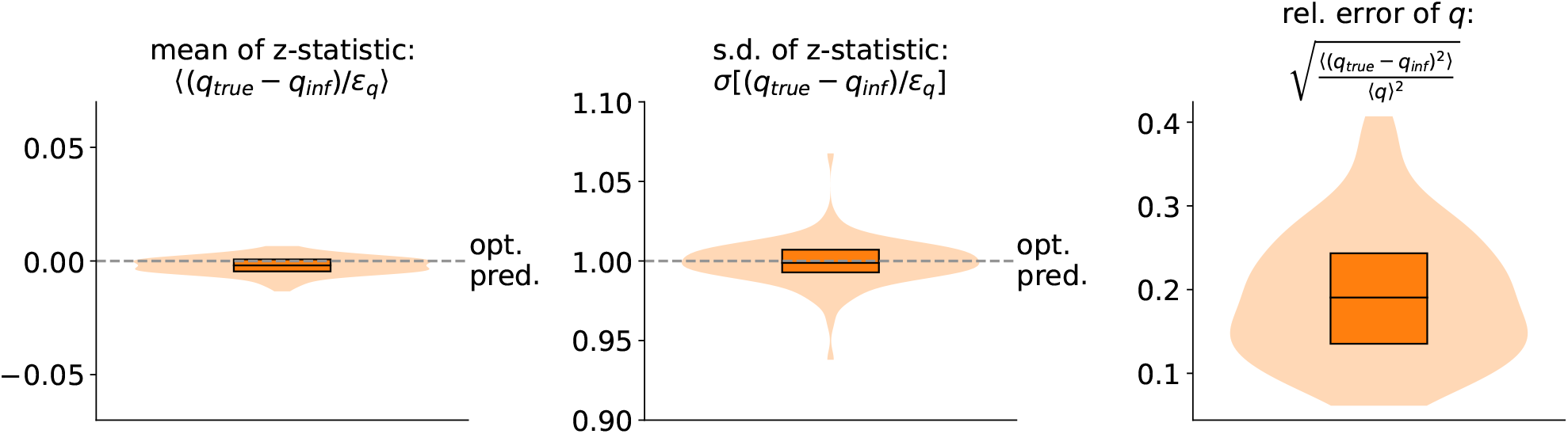
RealTrace accurately estimates volumic production rates across all 100 dataset with random parameters Θ. For each dataset, we compared all inferred volumic production rates *q*_inf_ with the corresponding ground truth *q*_true_ and the reported error-bar *ε*_*q*_, calculating a *z*-statistic *z* = (*q*_true_ − *q*_inf_)*/ε*_*q*_ for each observation. The figure shows the histograms (across the 100 datasets) of the average of *z* (left panel), its standard-deviation (middle panel) and the relative error (right panel). Each panel summarizes the histogram by a violin plot, its median (line), and its inter-quartile range (box). The figure shows that across all datasets, RealTrace neither systematically over-nor underestimates the growth rate (left panel), that the reported error bars correctly reflect the accuracy of the estimates (middle panel), and that the relative errors of the estimates in *q* tend to be around 20% for most datasets (right panel).

### B Supplementary results on mouse embryonic stem cells

The measured log size and fluorescence of five randomly chosen example cell cycles are shown in Fig. S8 together with the inferred, true log size and fluorescence. Fig. S9 shows the corresponding volumic production rates of the same example cell cycles. Note that volumic production rate as inferred by RealTrace smoothly varies over time (Fig. S9**a**). In contrast, naive estimates suffer from noise in both the size and fluorescence measurements and consequently jump around erratically (Fig. S9**b**). Remarkably, even averaging over 11 time points only partially removes the effects of the measurement errors and the averaged naive estimates of volumic production have little in common with the true underlying biological fluctuations.

Because RealTrace provides realistic estimates of the volumic production rate, it is possible to obtain a realistic distribution of these rates across cells and time (Fig S10**a**). RealTrace predicts that the most likely volumic production rate fluctuates between 0 and 1 (*µ*m^2^h)^*−*1^ with a steep tail at low rates and a much longer tail towards higher rates. In contrast, the naive, instantaneous volumic production rates fluctuate between *−*2 and 4 (*µ*m^2^h)^*−*1^ with many negative production rates, and averaging over multiple time points does not improve the estimation of the true distribution (Fig S10**b**). Averaging over multiple time points dampens the impact of measurement errors on the estimates for the volumic production rate but also averages true biological fluctuations. Thus, while the distribution for the naive rate becomes narrower as the estimates are averaged over an increasing number of time points, this is the result of smaller errors due to measurement noise and smaller biological fluctuations on longer time scales. Thus, to estimate the variability in the volumic production rate (or the growth rate) of single cells, averaging naive estimates does not overcome the measurement noise.

Finally, RealTrace allows us to calculate the auto- and cross-correlation functions of growth and production rates (Fig. S11). This allows us to quantify how quickly growth and volumic production rate fluctuations decay. The auto-correlation function of growth rates (Fig. S11**a**) shows that growth rate fluctuations drop exponentially on a time scale of about half an hour (1*/γ*_*λ*_ = 0.55 *h*) which corresponds to only 4 % of the typical cell cycle time, which is around 12 *h*. As shown in Fig. S11**b**, volumic production rate fluctuations last about twice as long (1*/γ*_*q*_ = 0.91 *h*). Even more interesting, we can also calculate the cross-correlation between growth and volumic production rate fluctuations at different time lags (Fig. S11**c**). We find that the volumic production rate shows a substantial positive correlation with growth rate in a time window of around 2 hours, with a peak in correlation at zero lag, i.e., growth and production rates at the same time point. This correlation of growth and volumic production at the same time point suggests that both rates might be affected by a common mechanism.

**Figure S8:**
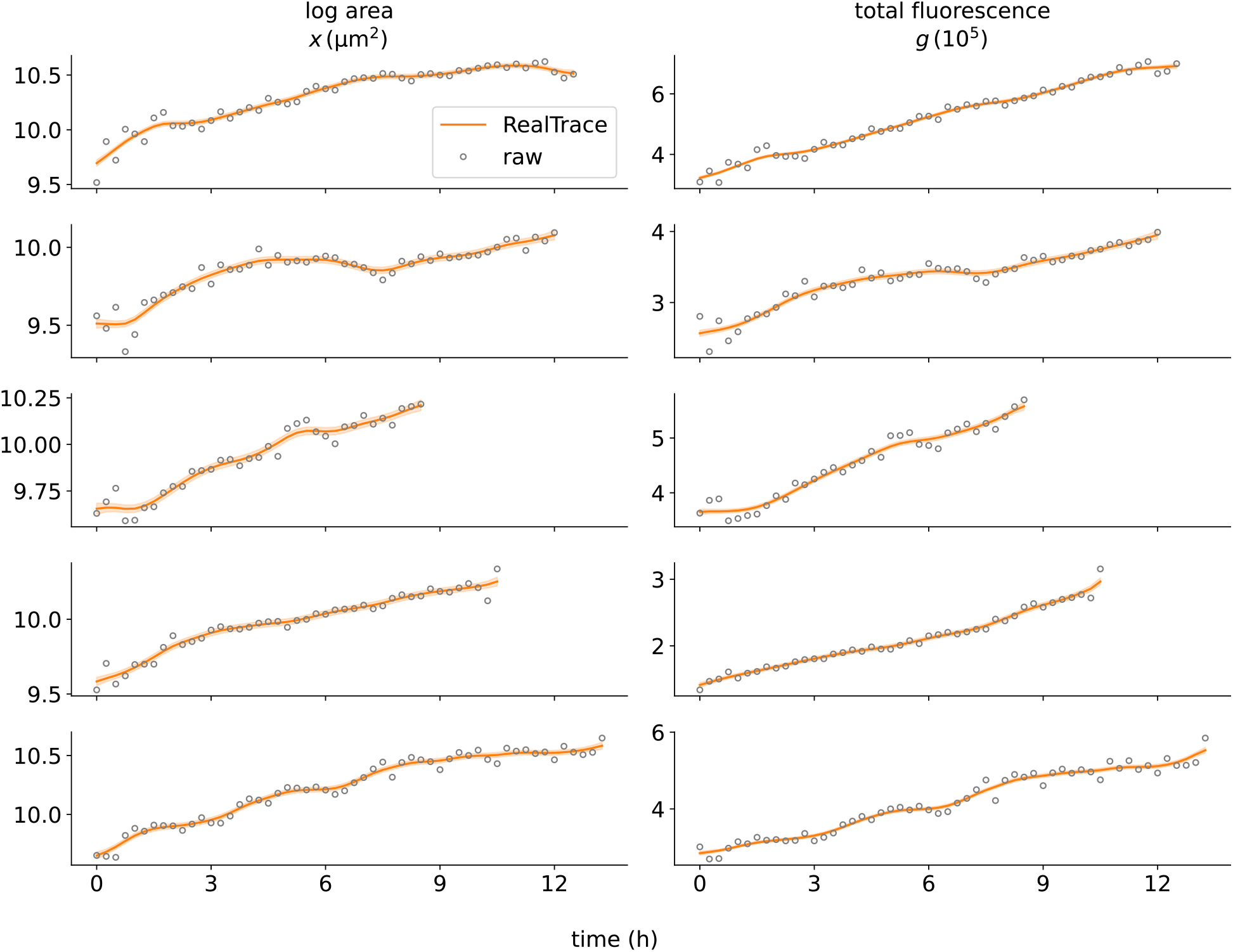
Example traces of log-area and total fluorescence single mouse embryonic stem cell nuclei. Each row is one example cell cycle. The grey circles represent the raw measurements, and the orange line shows the inferred true log area and total fluorescence as inferred by RealTrace, with error bars indicated by the transparent ribbons.

**Figure S9:**
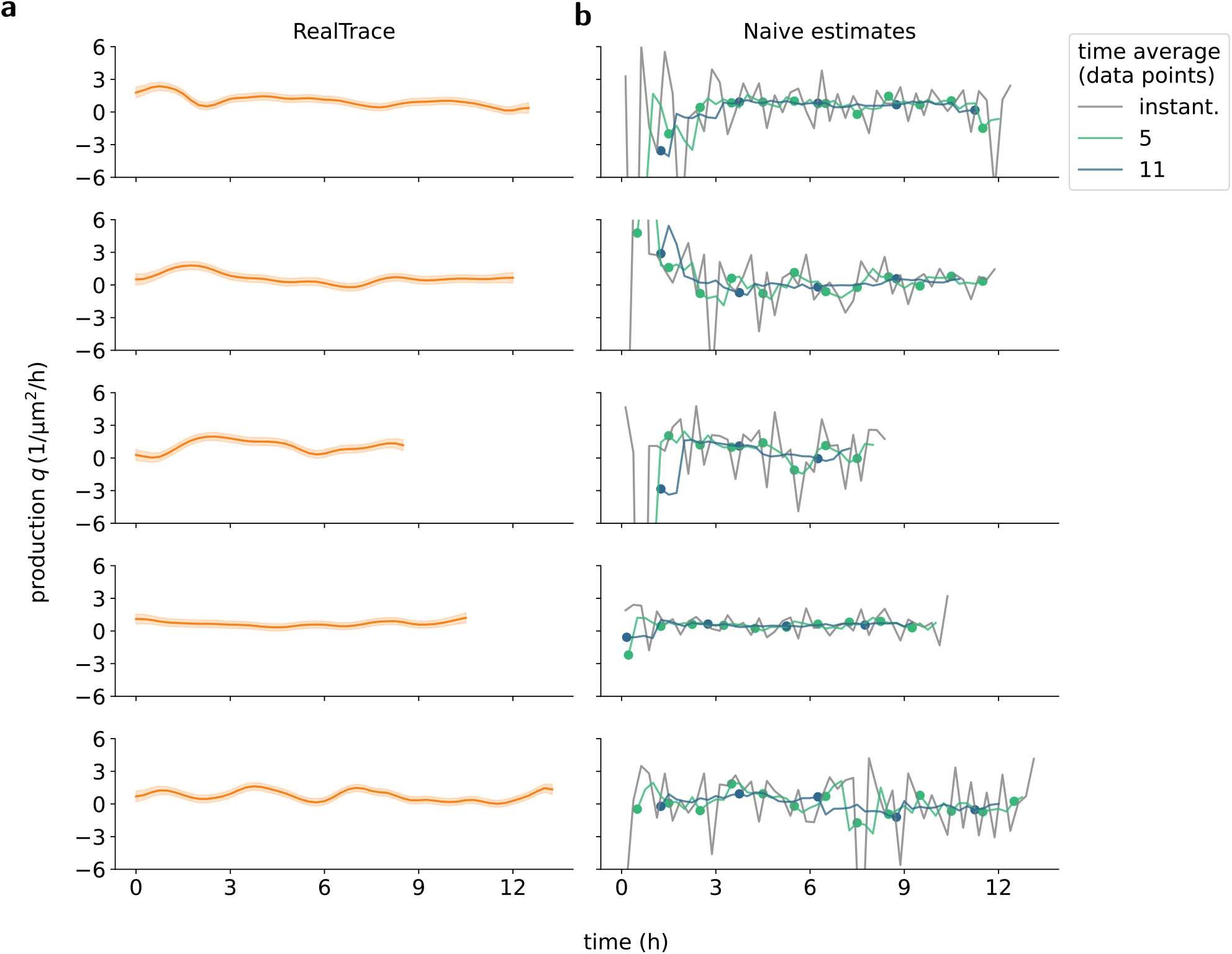
Volumic production rates of single mouse embryonic stem cell nuclei. The rows correspond to the same example cell cycles of Fig. S8. **a** Posterior means (lines) and standard deviations (ribbons) of volumic production rates inferred by RealTrace. **b** Grey traces show the volumic production rate calculated by taking into account only two consecutive time points. Traces in green and blue correspond to running averages taking into account five or eleven consecutive data points, respectively. Markers show the averaged volumic production rates of non-overlapping windows.

**Figure S10:**
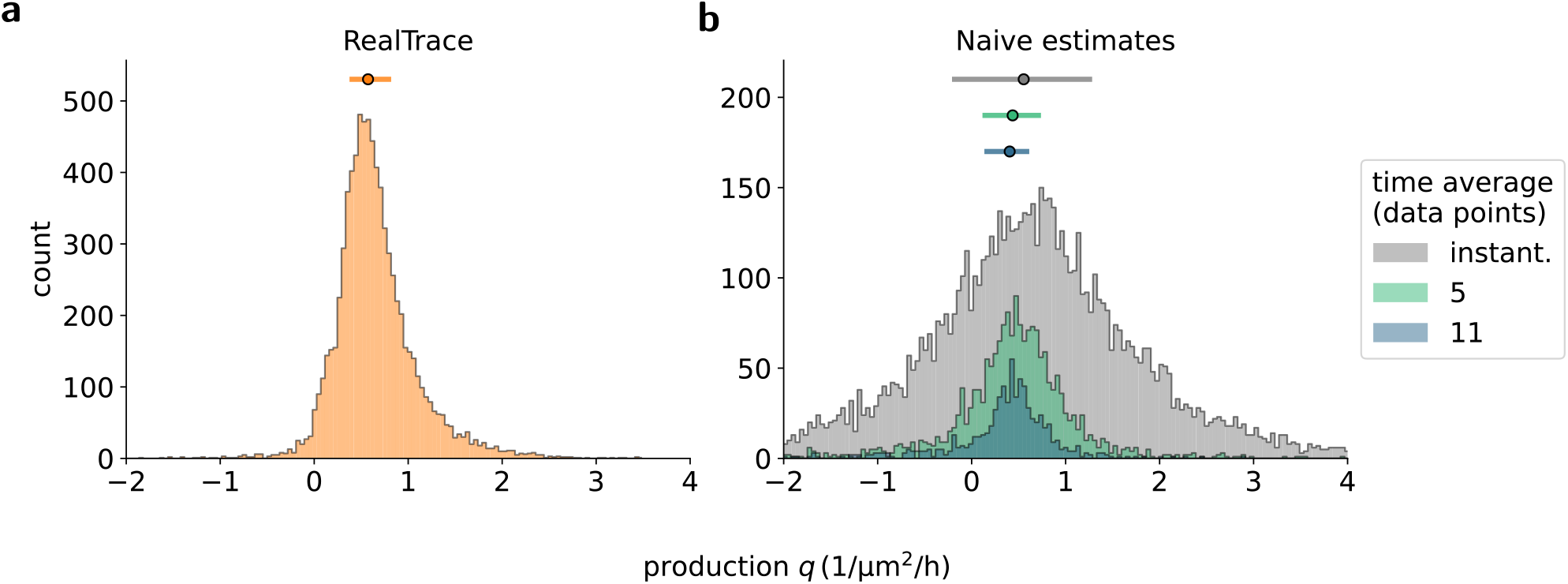
Distribution of the volumic production rate. **a** Distribution of volumic production rates that are estimated by RealTrace. Shown is the distribution across the mean of the posteriors for the volumic production rate. The circle and bar at the top indicate the median and interquartile range of the distribution. **b** Distribution of production rates that are naively estimated. The distribution in grey corresponds to the naive estimates calculated from two consecutive time points. The green and blue distributions correspond to naive estimates of the volumic production rates averaged over five and eleven time points of non-overlapping time windows, respectively.

**Figure S11:**
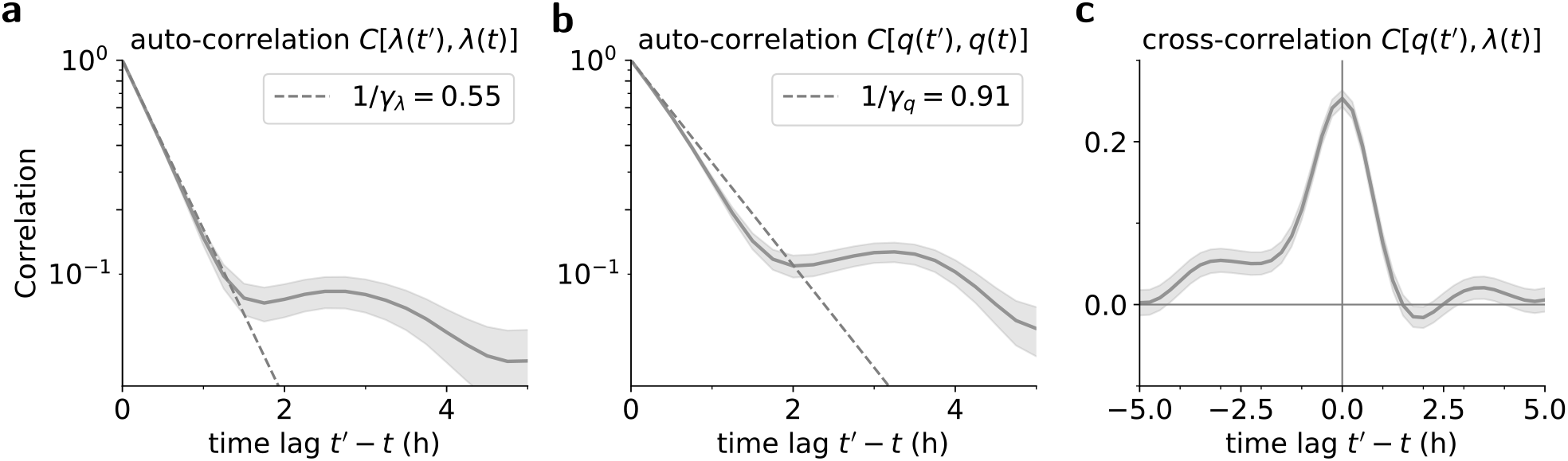
Correlation functions for mouse embryonic stem cells. **a** Auto-correlation function of the growth rate, the dashed line indicates the fitted correlation time scale of 0.55 h. **b** Auto-correlation function of the volumic production rate, the fitted correlation time scale of 0.91 h is significantly longer than for the growth rate fluctuations. **c** Cross-correlation function between the volumic production and the growth rate. The rates exhibit positive cross-correlations for time lags around 1 h in either direction, with the strongest correlation at the same time point, i.e., time lag zero.

### C Supplementary results on single *C. elegans* larvae traces

The *C. elegans* traces consist of the total volume of single animals and the fluorescence of GFP. The ubiquitously expressed GFP is used for tracking single animals. We, therefore, focus on the size dynamics and the growth of individuals.

Fig. S12 shows the growth rate of five randomly chosen example larvae between the 3rd and 4th moulting. The growth rate inferred by RealTrace shown in Fig. S12**a** shows that, across all larvae, the growth is fastest after the 3rd moulting, followed by a period of relatively constant growth before larvae enter the moulting phase and the growth rate falls below 0 around 80 % into the larval stage.

In contrast, naive estimates fluctuate strongly from one time point to the next, indicating that the measurement noise drives the growth rate estimates. Averaging the naive growth rate over windows of multiple time points dampens the impact of measurement errors, and the averaged rate exhibits smaller fluctuations from one window to the next. However, averaging over multiple time points both averages measurement errors and biological fluctuations. For example, the consistently high growth rate early in the larval stage that is clear in RealTrace’s inferences, cannot be reliably resolved by averaged naive estimates. Thus, while RealTrace can infer biological fluctuations on any timescale, naive estimates, which are averaged over many time points, are blind to true fluctuations on short time scales. For example, the consistently higher growth rate at around 10 % of the larval stage is not visible in the naive growth rate averaged over 11 time points.

We calculate the average growth between the 3rd and 4th moulting shown in Fig. S13. We find that the growth of individual larvae, as inferred by RealTrace, is similar to the average growth across many individuals. The average growth rate based on naive estimates displays a similar dependency on the larval stage. However, as already shown in Fig. S12**b**, the naively estimated growth rate of individuals is dominated by measurement errors. As a result, naive estimates make it difficult to assess how similar the growth rates of individuals across the larval stages are, which provides crucial insights into the size control of organisms ^13^.

**Figure S12:**
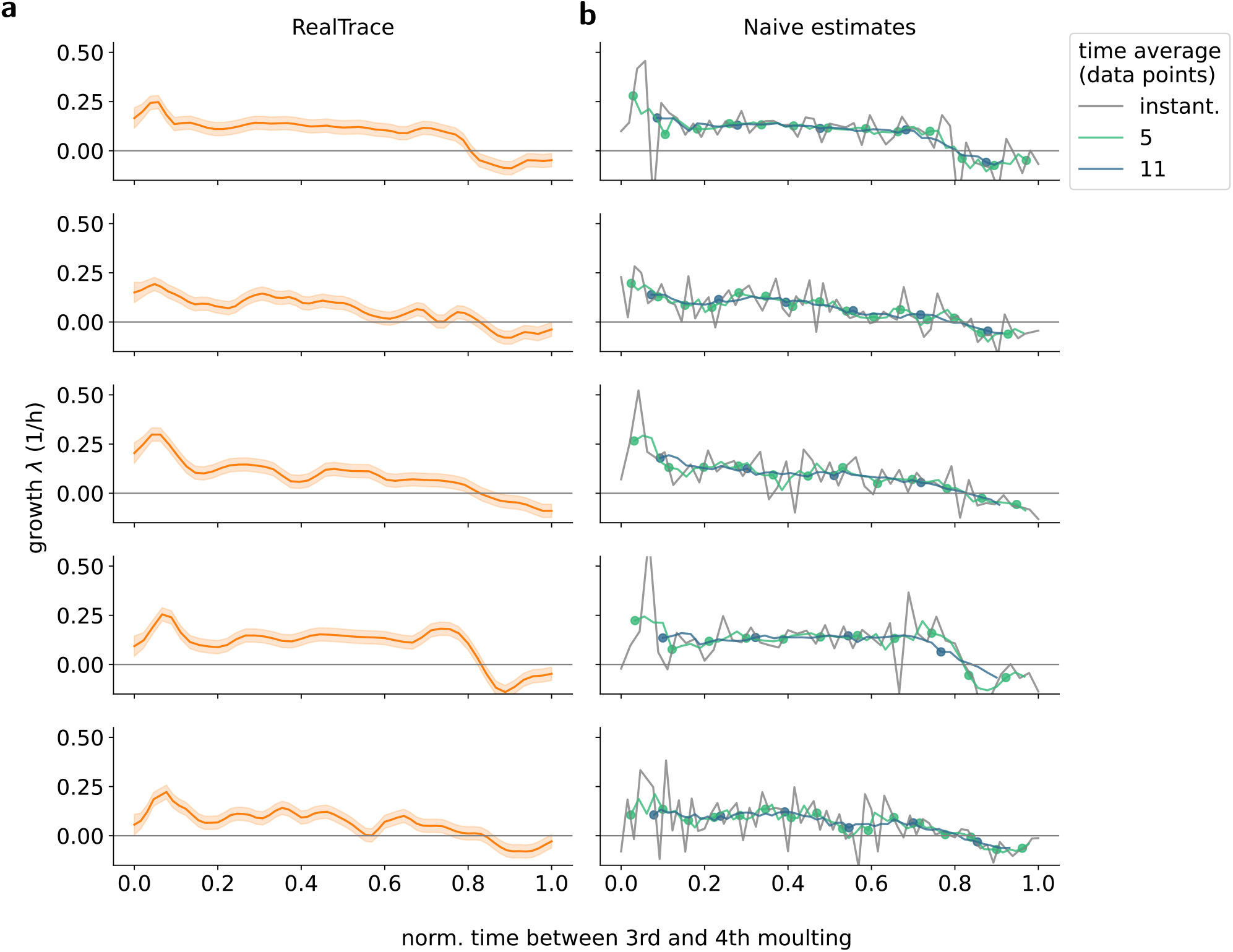
Growth of single *C*.*elegans* larvae between the 3rd and 4th moulting. **a** RealTrace estimate of the growth rate (posterior mean plus and minus one standard deviation). **b** Naive estimates for the growth rate. The traces in grey correspond to growth rates estimated from only two consecutive time points. Traces in green and blue correspond to running averages of the growth rate estimates taking into account five or eleven data points, respectively. Markers show the averaged growth rate of non-overlapping windows.

**Figure S13:**
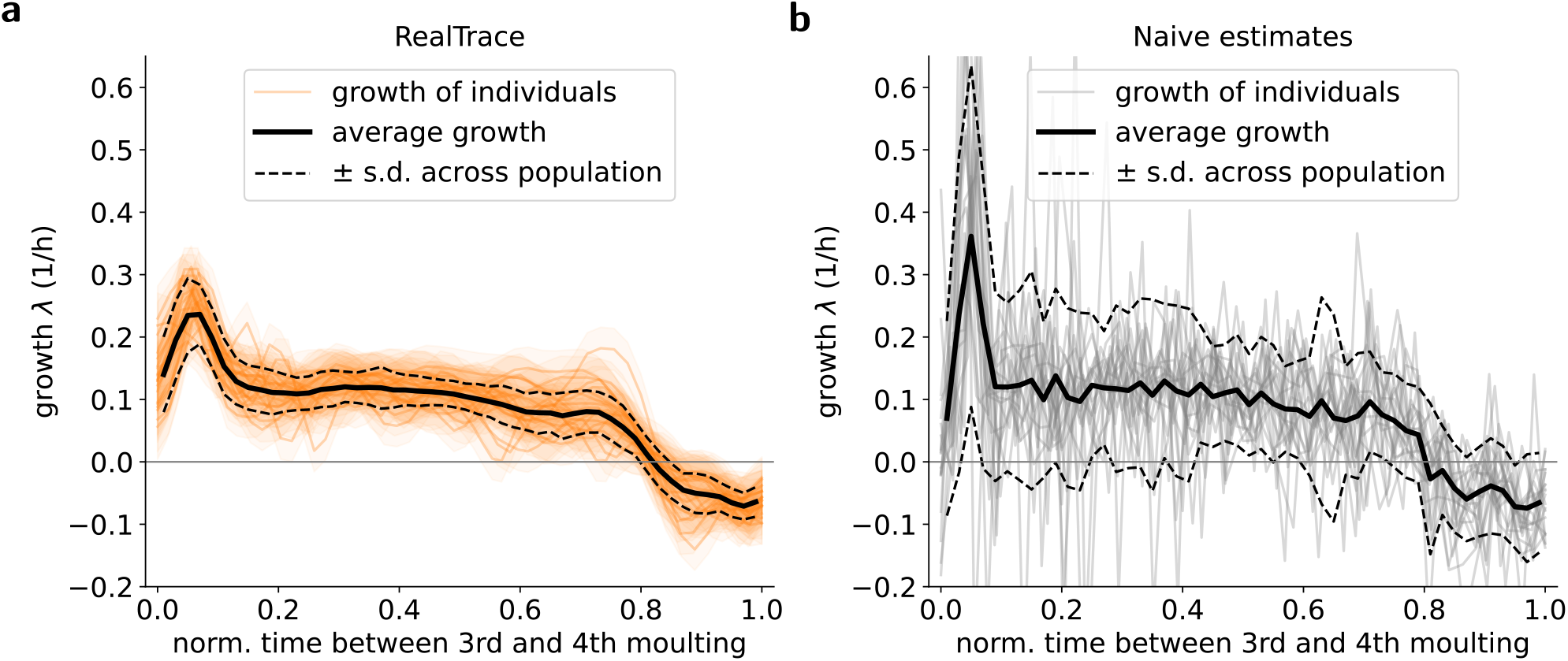
Growth of 20 single *C*.*elegans* larvae between the 3rd and 4th moulting. **a** The orange lines and shaded regions show posterior means and standard deviations for individual larvae. The black solid line and dotted lines show the average growth rate across individuals and its standard deviation, quantifying the variability across larvae. **b** Growth rates of individuals calculated by taking discrete derivatives of the log volume (grey lines). The solid and dashed black lines show the population average and the standard deviation across the population. The strong increase in estimated variability compared to RealTrace is driven by the measurement noise.

### D Supplementary figures for the *E. coli* datasets

**Figure S14:**
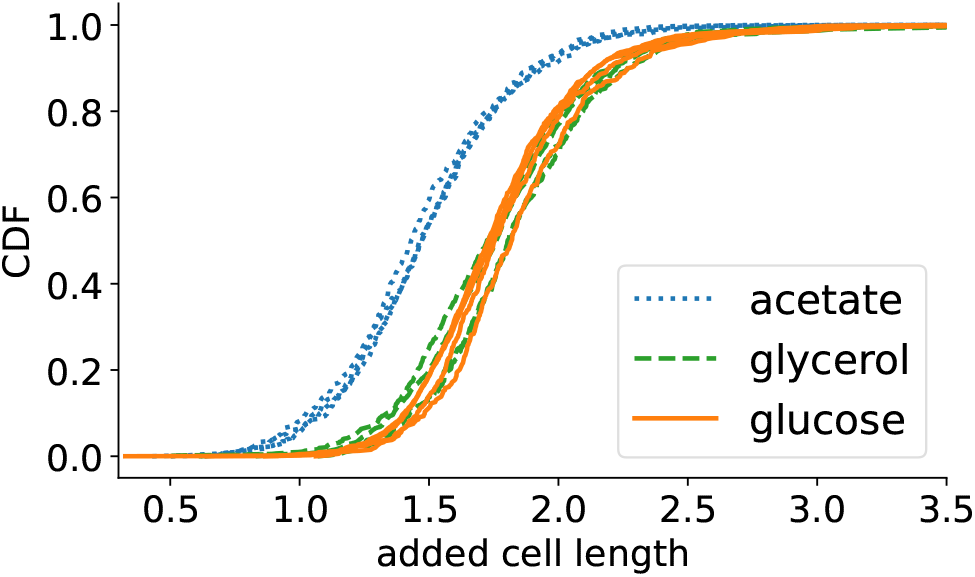
Cummulative distributions of added lengths of individual cell cycles for cells growing in acetate (blue), glycerol (green), and glucose (orange). Each line corresponds to the cumulative distribution of added lengths from one replicate experiment. Note that, even though the added length varies significantly across cell cycles, the distributions are highly reproducible across data sets.

**Figure S15:**
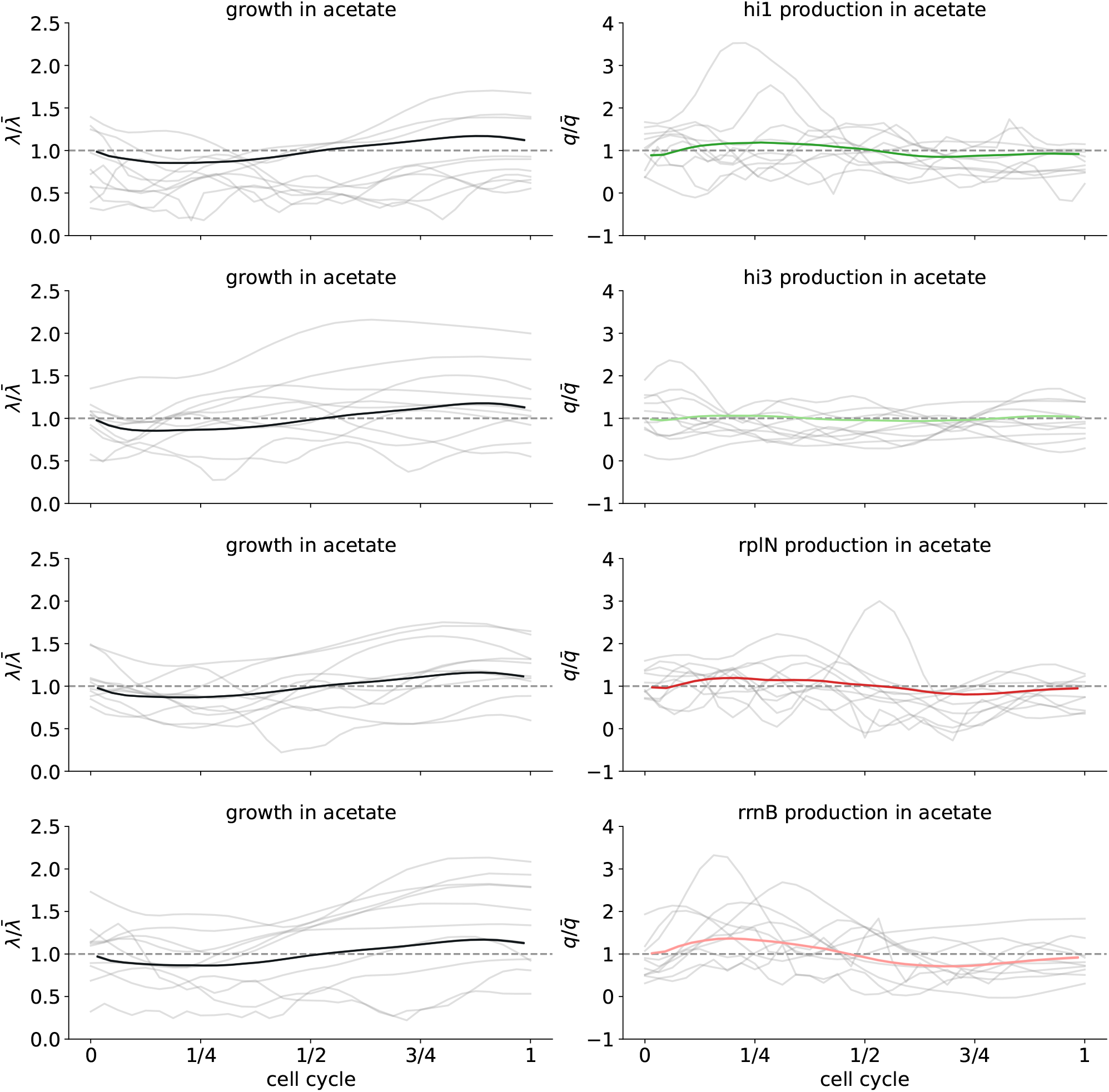
Growth and volumic production rates across the cell cycle for *E. coli* cells growing in minimal media with acetate. Each row corresponds to data from an experiment with a different transcriptional reporter (see titles). Left panels show the average growth rate as a function of cell cycle stage (black line) and 10 example single-cell traces (grey lines). Right panels show average volumic production as a function of cell cycle stage (green lines for synthetic constitutive promoters and red lines for ribosomal promoters) and 10 example single-cell traces of volumic production rate (grey lines).

**Figure S16:**
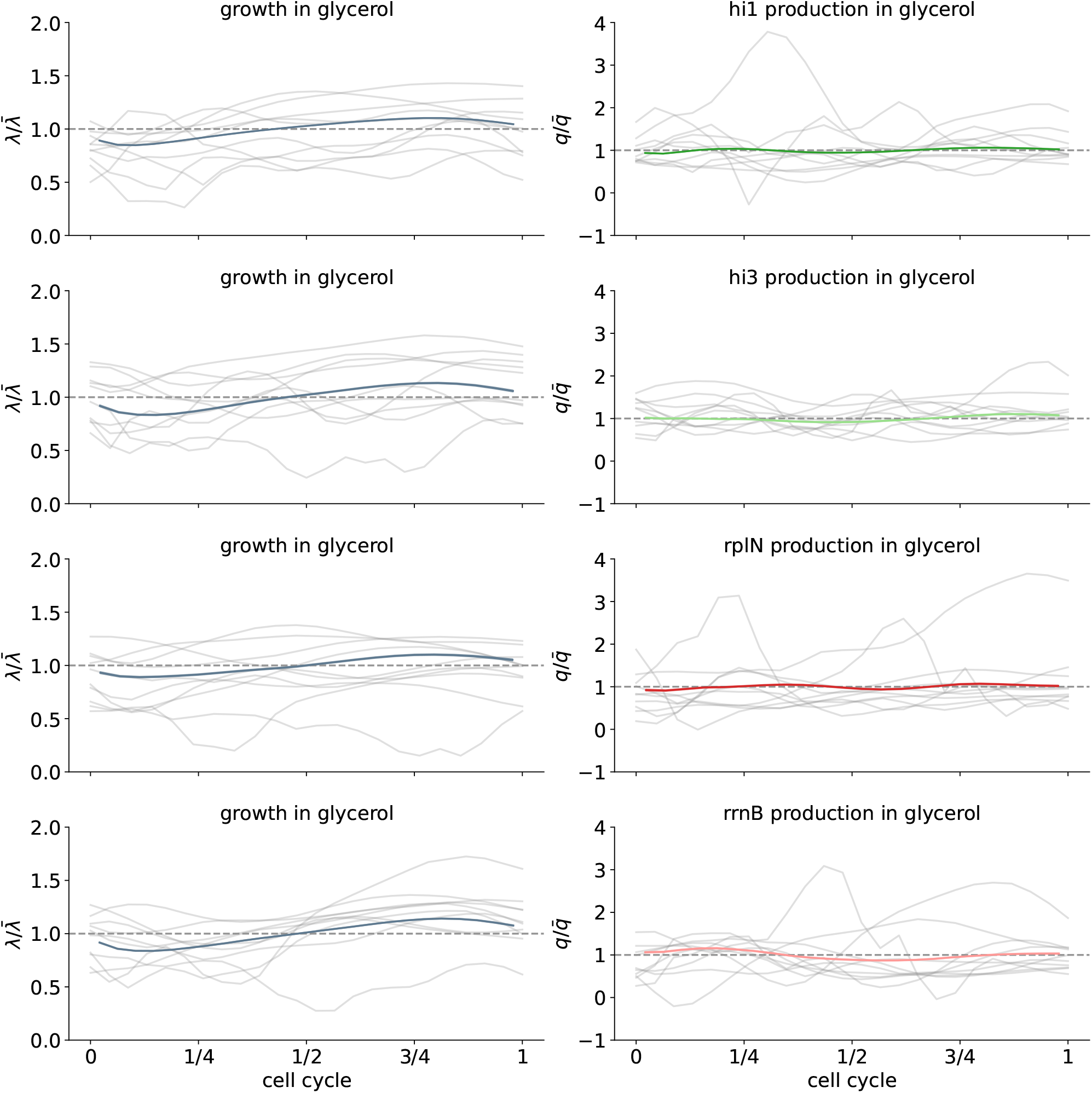
Growth and volumic production rates across the cell cycle for *E. coli* cells growing in minimal media with glycerol. Each row corresponds to data from an experiment with a different transcriptional reporter (see titles). Left panels show the average growth rate as a function of cell cycle stage (black line) and 10 example single-cell traces (grey lines). Right panels show average volumic production as a function of cell cycle stage (green lines for synthetic constitutive promoters and red lines for ribosomal promoters) and 10 example single-cell traces of volumic production rate (grey lines).

**Figure S17:**
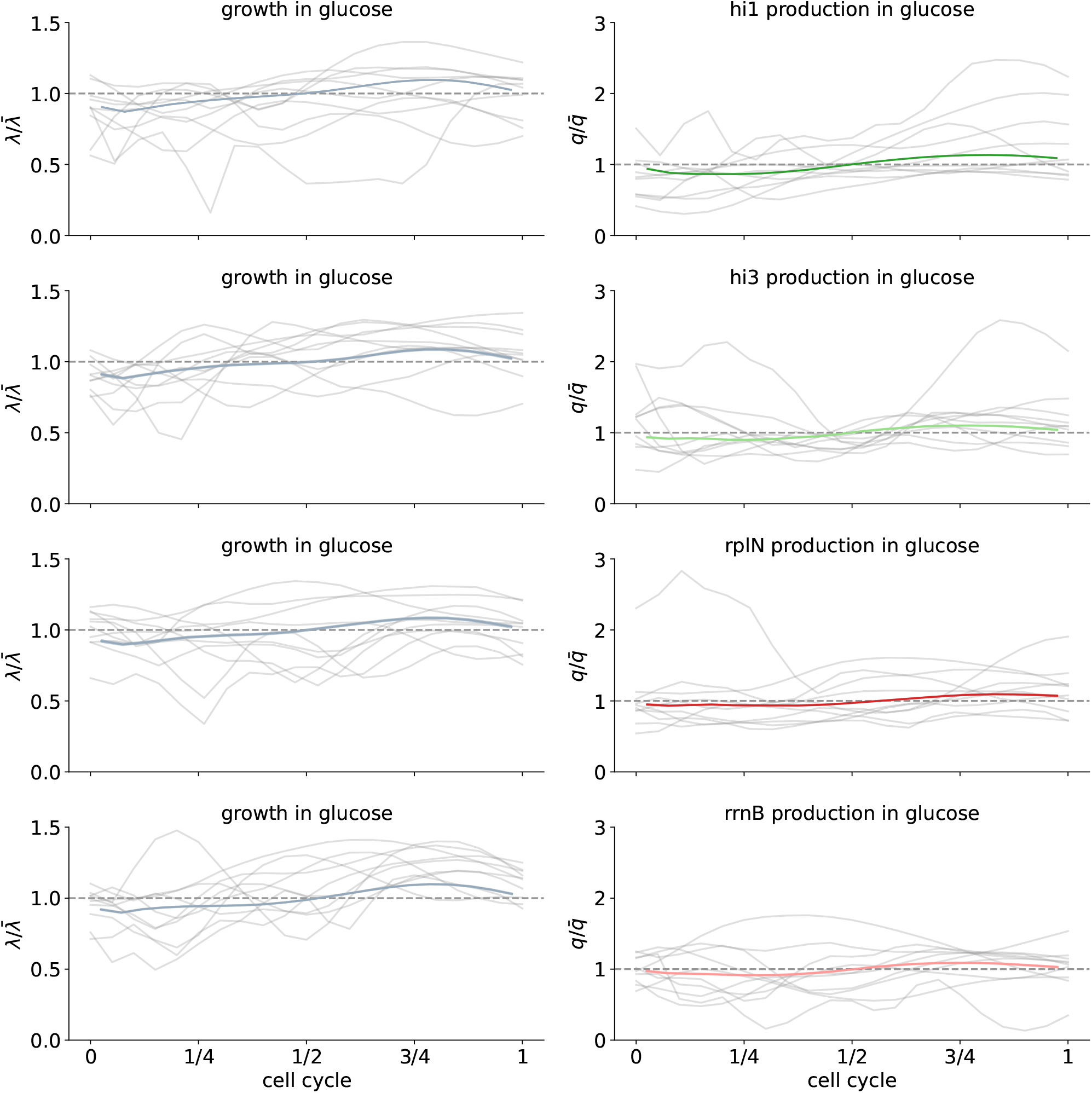
Growth and volumic production rates across the cell cycle for *E. coli* cells growing in minimal media with glycerol. Each row corresponds to data from an experiment with a different transcriptional reporter (see titles). Left panels show the average growth rate as a function of cell cycle stage (black line) and 10 example single-cell traces (grey lines). Right panels show average volumic production as a function of cell cycle stage (green lines for synthetic constitutive promoters and red lines for ribosomal promoters) and 10 example single-cell traces of volumic production rate (grey lines).

**Figure S18:**
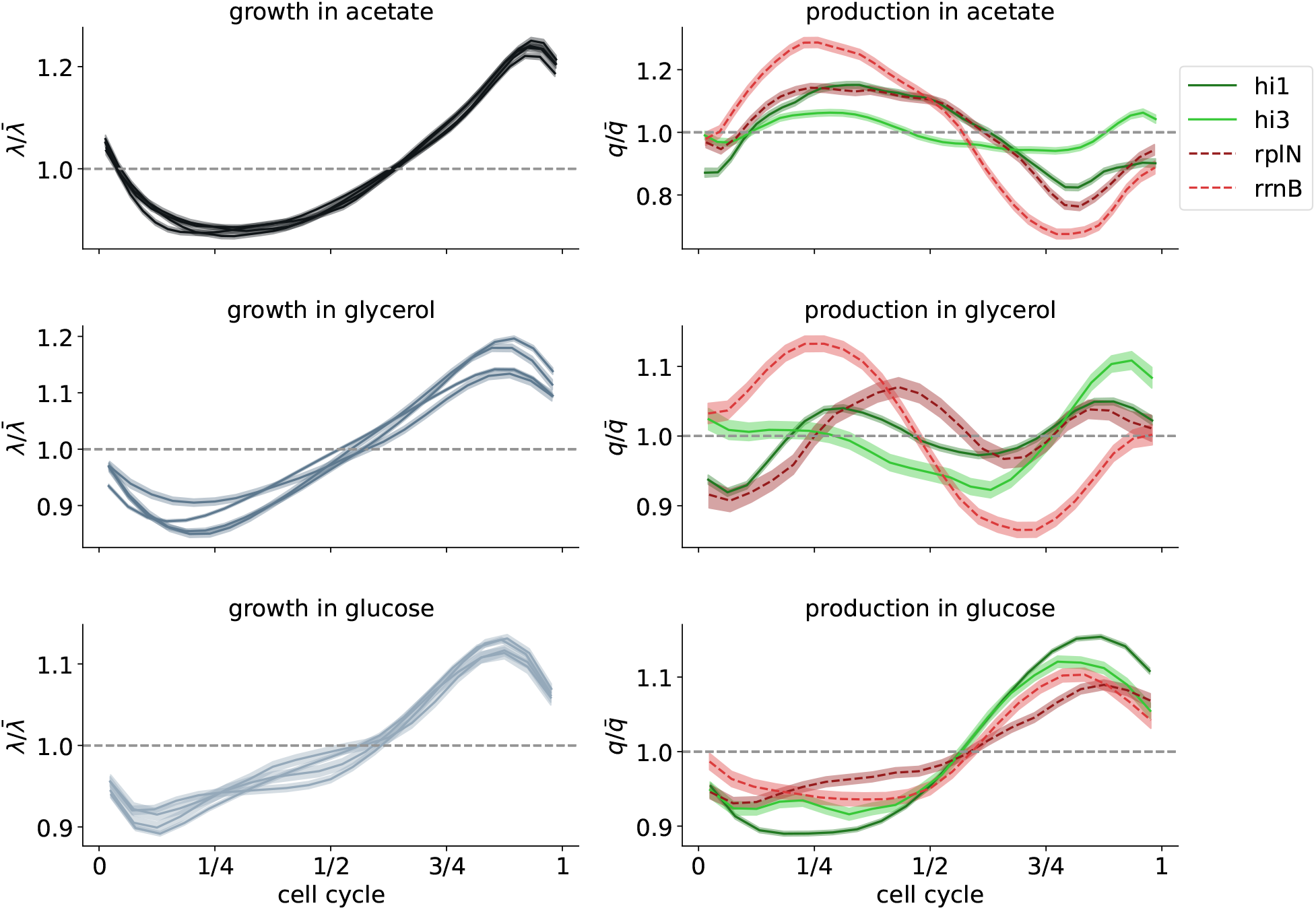
Cell cycle dependency of the growth and production rate when the cell cycle is parameterized by the time since birth. Specifically, the cell cycle is parameterized as (*t* − *t*_*b*_)*/t*_*cc*_, where *t* is the time, *t*_*b*_ is the time at birth, and *t*_*cc*_ is the total time of the corresponding cell cycle. As the total growth, the total added length per time, increases during the cell cycle, the patterns for the cell cycle dependencies are stretched early in the cell cycle, parameterized by the time compared to the cell cycle parameterized by the added length shown in Fig. 5.

### E Supplementary Methods

#### E.1 Simulated data sets

To validate the effectiveness and accuracy of the inference procedure we use simulated data sets in which the true biological dynamics is generated according to the stochastic process assumed by the process prior.

Note that at each cell division, there are two daughter cells and, depending on the experimental setup, in real datasets, some daughters may be tracked and others may not be. Consequently, in real datasets, cells are connected in a lineage tree and RealTrace has been designed to be able to analyze datasets with cells (or other biological objects) in a lineage tree of any topology. To mimic this feature of real data, we constructed simulated datasets with cells in random lineage trees as follows. Each data set consists of 10 independent lineages with 100 cell cycles in each lineage, so that the dataset consists of 1000 cell cycles in total. In order to avoid transients, we first simulate 5 cell cycles for each lineage, where, at each division, a random daughter is chosen to track further. The cells at the end of these 5 cycles are the roots of the lineage trees that will make up the dataset. We then create a lineage tree as follows. We pick a random cell from all cells that currently exist and simulate it until it divides. During the cell cycle, the stochastic dynamics of the cell’s state are integrated using an Euler scheme with a time step of *dt* = 0.001 min. When the log size has increased by 1, the cell divides. The sizes of the daughters are chosen from the Gaussian distribution with variance 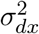, the fluorescence is converted into GFP copy number by dividing by 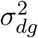, and each GFP molecule is then randomly assigned to a daughter with probability proportional to daughter size. The other cell state variables (i.e. growth rate and volumic production rate) are copied from the mother to both daughters. After each division, we then randomly pick one of the existing cells to simulate the next cell cycle. By iterating this, we create a random lineage tree with 100 cell cycles.

The parameters of each simulated dataset were drawn uniformly randomly in log-space from a given range around realistic parameter settings as listed in Table 2. For the parameters of the stochastic dynamics of growth rate and volumic production rate we randomly choose a mean, an inverse correlation time, and a coefficient of variation. These then set the amplitudes of the white noise terms *σ*_*λ*_ and *σ*_*q*_.

For the numerical optimization, the parameter is initialized in the following way. The means, namely 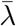 and 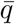, are directly estimated from the noisy data. The remaining parameters can not be estimated easily in practice, and, therefore, we initialize them by adding noise in log space to the true parameters. Specifically, we add Gaussian noise with a standard deviation of 1. The distributions of initial parameters are shown in Fig. S1. The parameters *σ*_*λ*_, *σ*_*q*_, *σ*_*x*_, and *σ*_*g*_ are squared in the implementation of RealTrace and are initialized and optimized as such, ensuring that the standard deviations *σ*_(*·*)_ are always positive. Yet, for clarity, we plot the standard deviations in Fig. S1. As a result, the distributions for the initial parameters of these parameters are narrower in this plot. For the optimization, very broad bounds were set, corresponding to a 5 % and 20-fold of the respective initial value.

The parameters of the synthetic data set mimicking the dynamics of *E. coli* growing on glucose and expressing GFP under the control of *hi1* are given in Table 1. We simulate 1000 cells after discarding the first 20 cell cycles. Cells divide after adding a fixed log-length of log(2). To simplify the comparison with the naive estimates, in this simulation, we set the bleaching to 0 (see also Sec. E.2).

#### E.2 Comparison of RealTrace estimates and naive estimates

The comparisons of the RealTrace and naive estimates shown in Fig. 2**b** and **c** require calculation of the naive estimates and averaging over windows of different numbers of consecutive data points. The naive estimates of volumic production rate are calculated by taking the difference in measured fluorescence at consecutive time points and dividing it by both the length of the time interval and the size of the cell. In particular, the volumic production rate 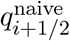 between time points *i* and *i* + 1 is calculated as

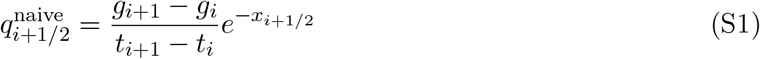

with the interpolated log-length given by

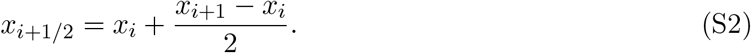

Note that if there were bleaching, we would also have to correct this naive estimate for the bleaching between consecutive time points, which would be an additional source of errors. We therefore set the bleaching rate to 0 in these simulations so that the accuracy of the naive estimates in this scenario is an upper limit. To estimate production rates for pairs of data points including a cell division, we assume that cells divide symmetrically.

To compare average volumic production rates over sliding windows of different sizes we calculate 1. the true average production rate by averaging the true production rate over the time points in the window, 2. RealTrace’s estimate of the average production rate by a weighted average of the posterior means taking into account their error-bars, and 3. an average of the naive estimates of production rate over the time points in the window. Note that sliding windows may include a cell division. In this case, we first average the production rate over the daughter cells, such that all time points contribute equally to the average of the sliding window. Importantly, we calculate the sliding average for the true, RealTrace-inferred, and naive estimates in the same manner.

Finally, in Figs. S6, S5 and S4 we show naive estimates for the growth rate, which are calculated according to

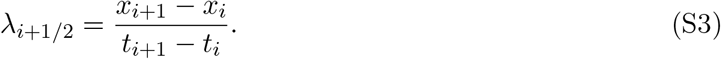

#### E.3 Cell cycle dependencies

To calculate cell cycle dependencies of the growth and production rate, we parametrize the cell cycle by the added length since birth. We quantify the stage of cell cycle progression *ϕ ∈* [0, 1] as the added length since birth *δl* divided by the total added length Δ*l* in the corresponding cell cycle, i.e. *ϕ* = *δl/*Δ*l*. Thus, for a given cell, its measurement time points *i* are mapped to values *ϕ*_*i*_ = *δl*_*i*_*/*Δ*l* in the interval 0 to 1.

To calculate average growth rate (or volumic production) as a function of cell cycle stage we need to take into account that fast growing cells will have less measurement time points per cell cycle than slow growing cells. Consequently, if we were to count every measurement towards the average, the average would be biased toward the statistics of slower growing cells. We thus proceed as follows. We first calculate the average number of time points *n* per cell cycle. We then pick *n* + 1 values *ϕ*_*k*_ = *k/n*, with *k* running from 0 to *n*. For each cell, we then estimate the growth rate (or volumic production rate) at each stage *ϕ*_*k*_ by linearly interpolating between the last measurement time point before *ϕ*_*k*_ and the first measurement time point after *ϕ*_*k*_ in the cell cycle of the corresponding cell. To take the error-bars of the estimates into account, this estimate is weighted with the inverse of the variance of its posterior, which in turn we estimate as the average of the variances of the time points before and after *ϕ*_*k*_. In this way, each cell contributes exactly one observation for each value of *ϕ*_*k*_. We similarly computed the cell cycle dependency parametrized by the time since birth in Fig. S18.

#### E.4 Decomposing the posterior distribution

To incorporate all data points into the posterior distribution of a time point **z**_*n*_ the posterior is decomposed into a forward and a backward part. The posterior conditioned on all data points *D*_1,…,*N*_ can be written as

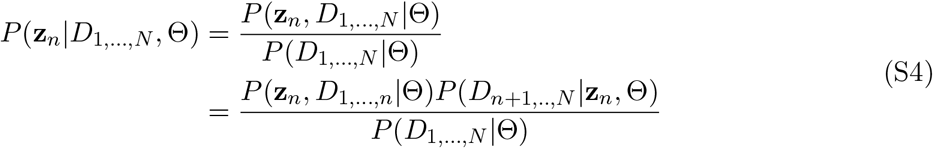

This decomposition of *P* (**z**_*n*_, *D*_1,…,*N*_ |Θ) into a part including all data points up to *n* and a part including the data points from *n* + 1 to *N* holds because the model is Markovian. Using Bayes’ theorem, we can rewrite *P* (*D*_*n*+1,..,*N*_ |**z**_*n*_, Θ) in terms of the posterior *P* (**z**_*n*_|*D*_*n*+1,..,*N*_, Θ) as follows

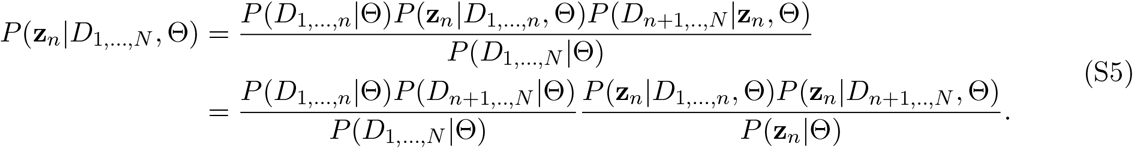

Note that the first ratio in the above expression is just a normalizing factor that does not depend on **z**_*n*_. Thus, to calculate the posterior over all data we just need the posterior conditioned on all the past, the posterior conditioned on all the future, end the prior probability *P* (**z**_*n*_|Θ) given no data. This prior distribution *P* (**z**_*n*_|Θ) is the product of a uniform distributions over *x* and *g*, and the stationary distributions for the OU processes of *λ* and *q* given the parameters of the prior. Thus, the priors for *λ* and *q* are Gaussian distributions with means 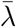 and 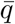 and variances 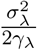 and 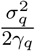 respectively.

#### E.5 Likelihood and posterior distribution calculation

The total likelihood

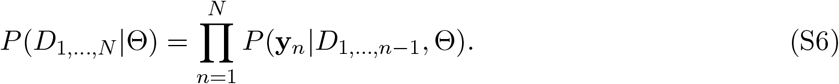

is maximized numerically to find the optimal parameters Θ^*∗*^ using the Nelder-Mead simplex algorithm ^50,51^. In this section, we explicitly show the calculation of the likelihood of a single time point *P* (**y**_*n*_|*D*_1,…,*n−*1_, Θ).

We assume Gaussian measurement noise, thus the probability of the measurement **y**_*n*_ given a true state of the cell **z**_*n*_ reads,

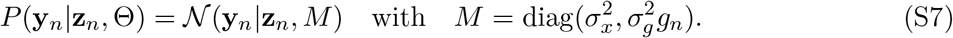

To calculate the likelihood of the measurement, we need to solve the integral,

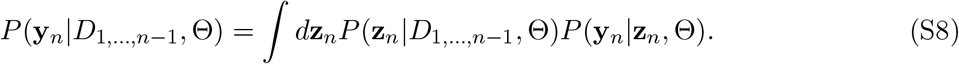

The prior distribution is Gaussian, with mean ⟨**z**_*n*_⟩ and covariance Cov(**z**_*n*_, **z**_*n*_),

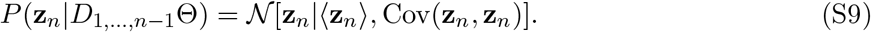

Since the measurements are only taken on *x* and *g*, we define a mean vector and covariances matrix over a subset of entries of the prior distribution,

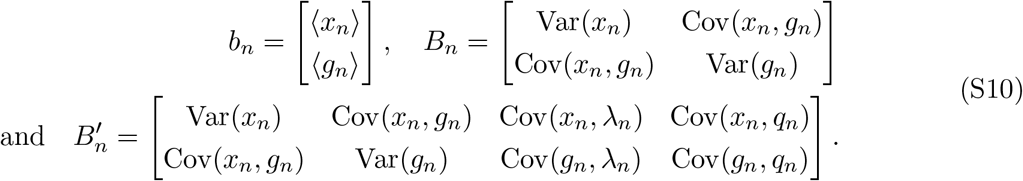

Then, the likelihood of the measurement **y**_*n*_ reads,

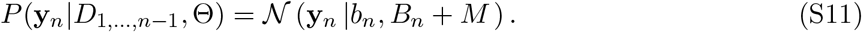

Here, we assumed that the variance Var(*g*_*n*_) is small compared to the mean ⟨*g*_*n*_⟩ and used the approximation,

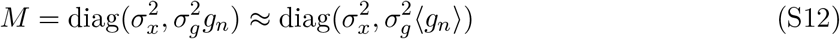

to perform the integral. In order to move forward to the next time point, we first calculate the posterior distribution incorporating the measurement **y**_*n*_,

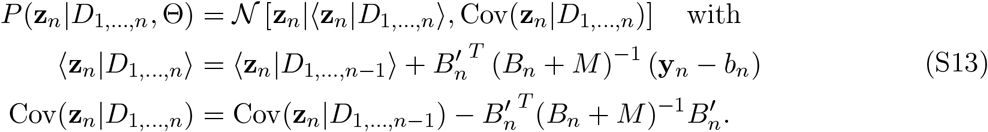

Given the posterior distribution for **z**_*n*_, the prior distribution for the next time point **z**_*n*+1_ is given by,

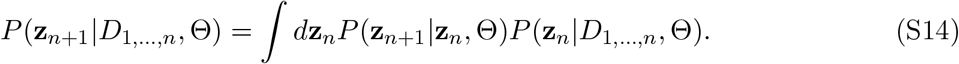

The conditional probability *P* (**z**_*n*+1_|**z**_*n*_, Θ) is approximated by a Gaussian distribution, whose mean vector and covariance matrix are determined by the model. The calculation of this prior distribution is shown in the next sections.

#### E.6 Initialization of the prior distributions at the first and last time points

The prior distribution for the first time point

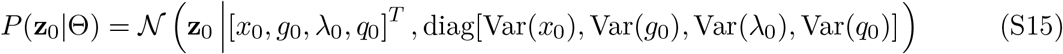

of each cell lineage is calculated as follows. The means and variances of the OU processes, namely for *λ*_0_ and *q*_0_ are given by their respective stationary probability distributions,

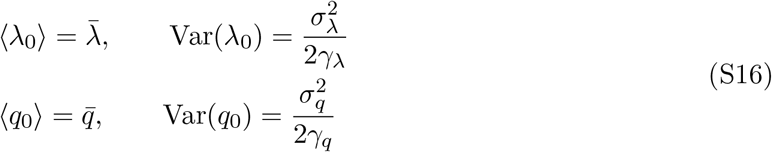

The mean and variances for *x*_0_ and *g*_0_ are estimated from the data itself. The distribution for the first time point of the cell lineage is estimated from all measurements of *x*_*i*_ and *g*_*i*_ at cell birth in the data set, denoted as **x**_*birth*_ and **g**_*birth*_ respectively.

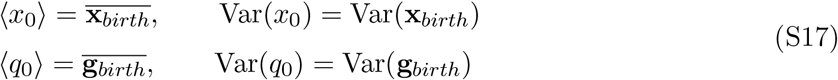

Similarly, for the calculation of the backward direction, the distributions for *x* and *g* are estimated from measurements at cell division. The distributions of *λ* and *q* are the same as for the forward direction.

#### E.7 Calculation of the prior distribution over one measurement time interval

##### E.7.1 Prerequisite for the calculation of the mean and covariance of the prior distribution

In this section, we show the derivation of the new prior distribution *P* (**z**_*n*+1_|**z**_*n*_, Θ) at time point *t*_*n*+1_. To simplify the notation, we will consider the step from *t*_*n*_ = 0 to *t*_*n*+1_ = *t* without loss of generality. The solutions of the model for a given **z**_0_ = (*x*_0_, *g*_0_, *λ*_0_, *q*_0_) are

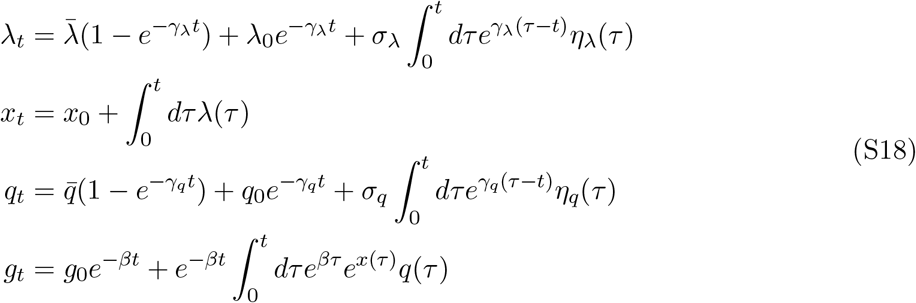

Notably, the equation for the GFP content *g*_*t*_ includes a term that non-linearly depends on the log-size *x*(*t*) through the integral of *e*^*x*(*τ*)^. To keep the calculation analytically tractable calculate we use the approximation *x*(*τ*) = *x*_0_ + *λ*_0_*τ* . That is, we approximate the cell size by assuming a constant growth rate *λ*_0_ between time zero and tine *t*. The solution for *g*_*t*_ then becomes

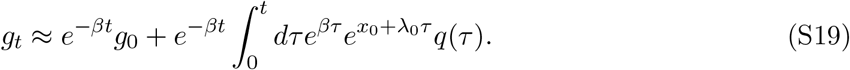

In the following derivations, we will use this approximation.

These equations define a mean and covariance of the cell’s state conditioned on the previous time points that we will calculate in the next sections. These means and covariances uniquely define a Gaussian. Thus, we approximate the conditional distribution *P* (**z**_*t*_|**z**_0_, Θ) as a Gaussian. We then integrate over the cell state **z**_0_ weighted by the posterior for **z**_0_, which is itself a Gaussian. We denote the means 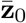 and covariances *C*_0_ of the posterior of **z**_0_ as,

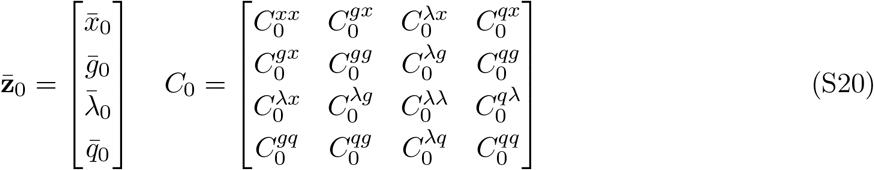

In the calculations, we will make use of the following, known integrals,

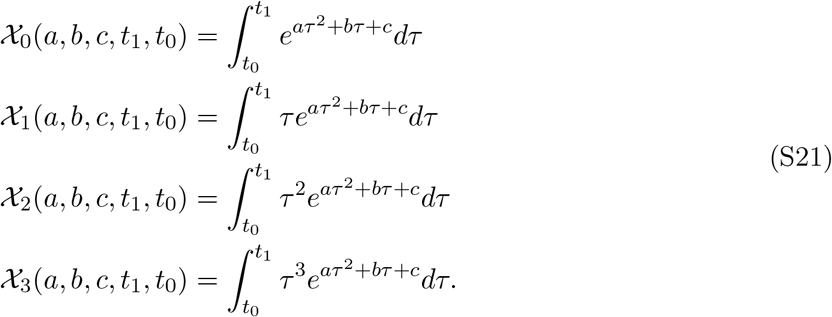

##### E.7.2 Mean function of the prior distribution

To calculate the mean of the prior distribution

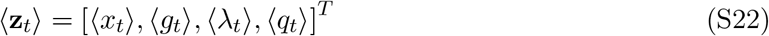

we first calculate the mean of the probability distribution conditioned on the initial condition **z**_0_ and then use the general identity

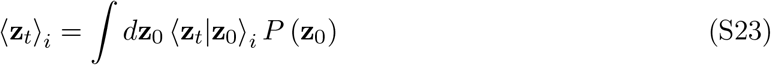

to obtain the marginal distribution *P* (**z**_*t*_|*D*_1,…,*t*_, Θ) (here *D*_1,…,*t*_ denotes the measurements up to the time *t*). Here, *P* (**z**_0_) is the forward part of the posterior of the time point 0. To illustrate the procedure, we calculate the mean function of the growth rate. For the conditioned mean, we find

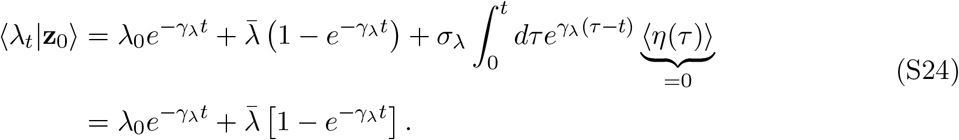

To calculate the mean of the marginal distribution, we use Eq. (S23) and Wick’s probability theorem and find,

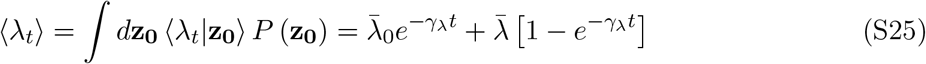

The remaining mean functions are calculated in a similar way,

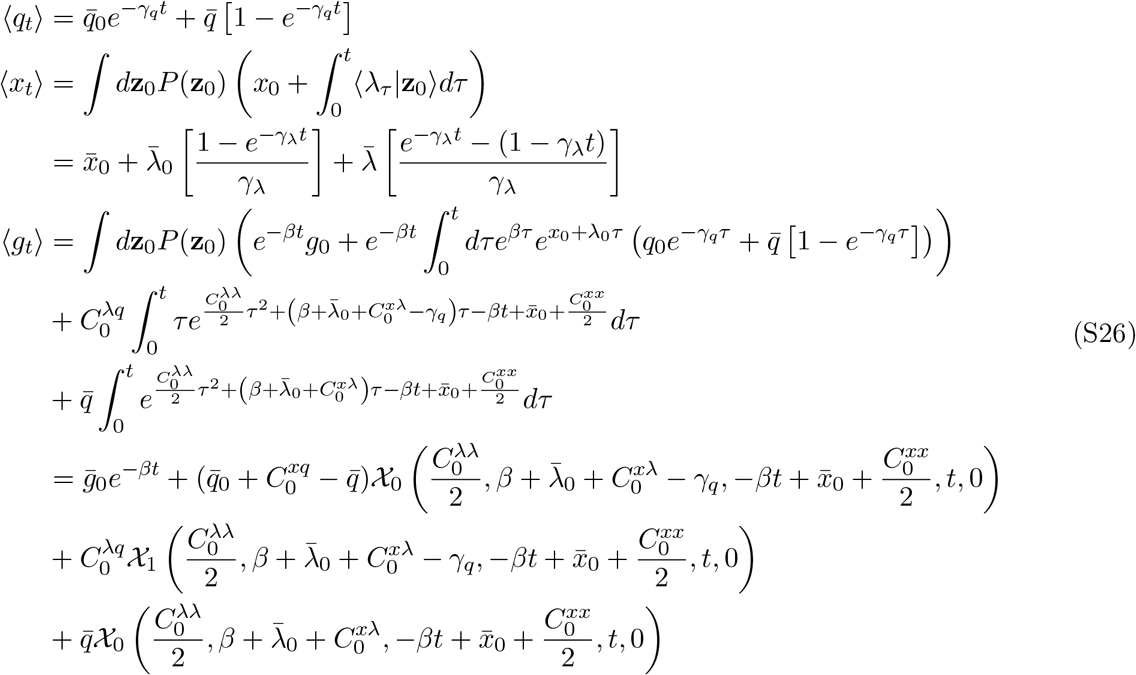

##### E.7.3 Calculation of the covariance matrix

The calculation of the covariance matrix of the prior follows a similar strategy as the calculation of the mean functions. The covariance elements that need to be calculated are

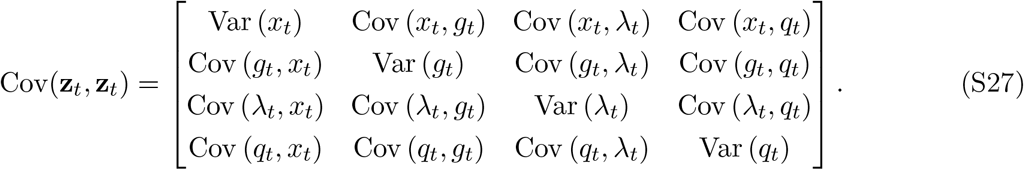

We calculate the covariance of the conditioned probability distribution, i.e. Cov(**z**_*t*_, **z**_*t*_|**z**_0_) and then use the identity

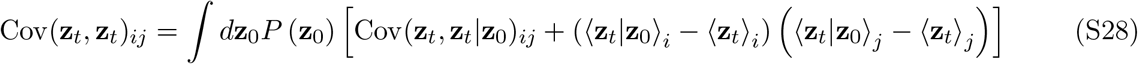

to calculate the covariance matrix of the new prior distribution. To solve the integral we use Wick’s probability theorem. This procedure is similar to the calculation of the means and we illustrate the calculation showing the derivation for Var(*λ*_*t*_). The conditioned variance of *λ*_*t*_ reads

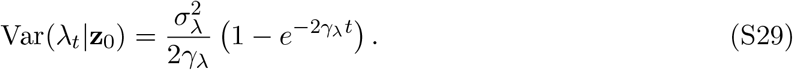

To calculate the unconditioned variance, we use Eq. (S28) and find

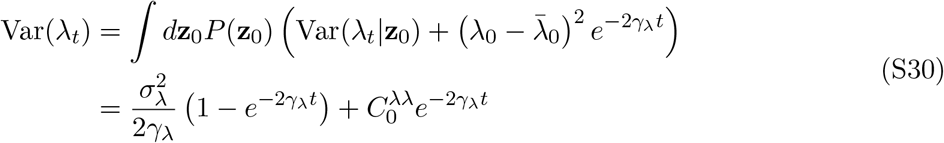

The other entries of the covariance matrix can be calculated in a similar way. However, the covariance elements including *g*_*t*_ are very lengthy, which is why they are calculated with the aid of Mathematica (see `calculations_notes` in https://github.com/nimwegenLab/RealTrace)). The Mathematica script performs the following steps: Calculate the covariance function using Eq. (S28), integrate over **z**_0_, and finally integrate over time. The remaining covariance elements are,

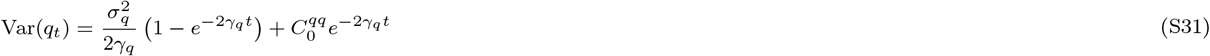

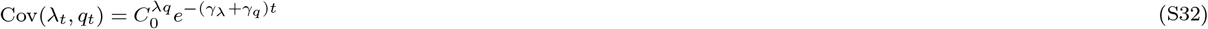

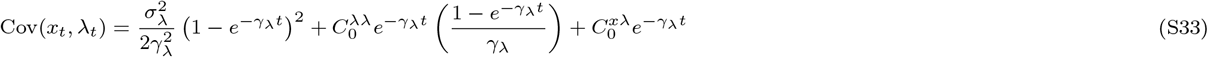

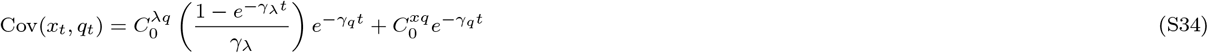

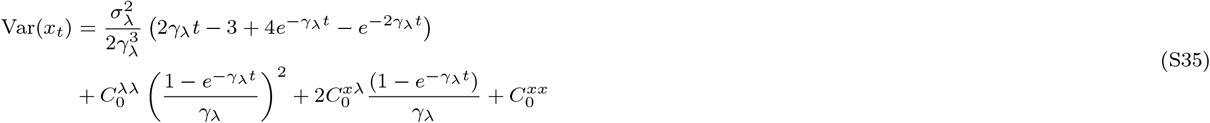

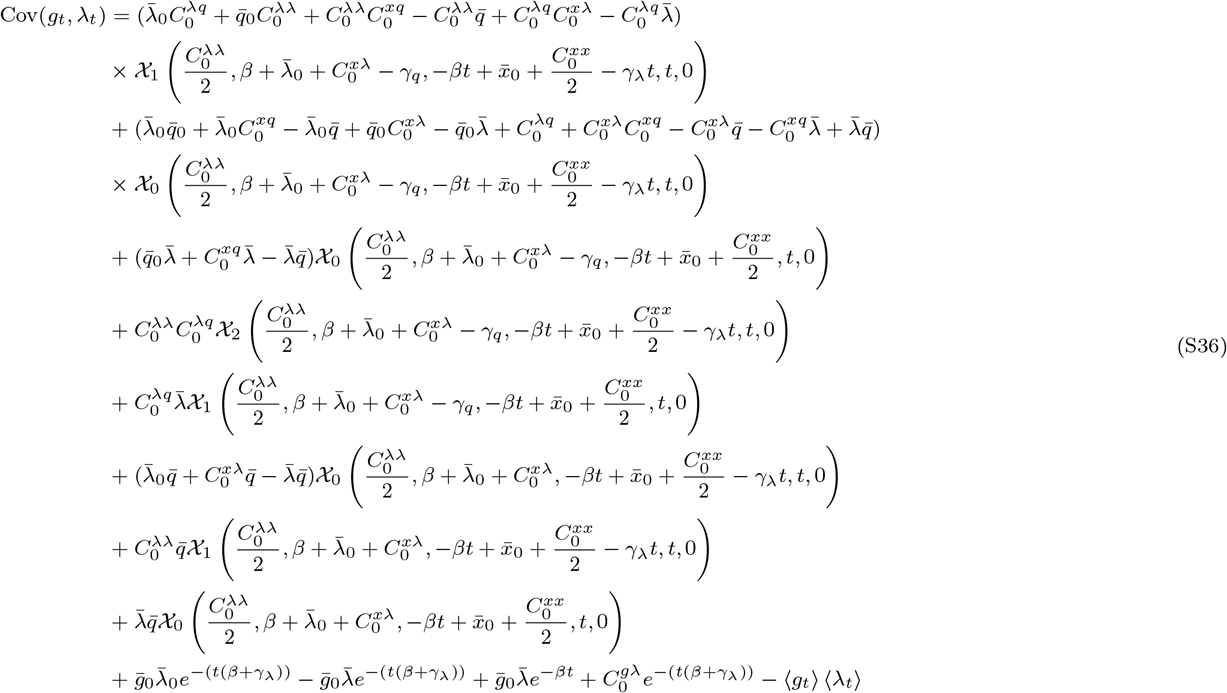

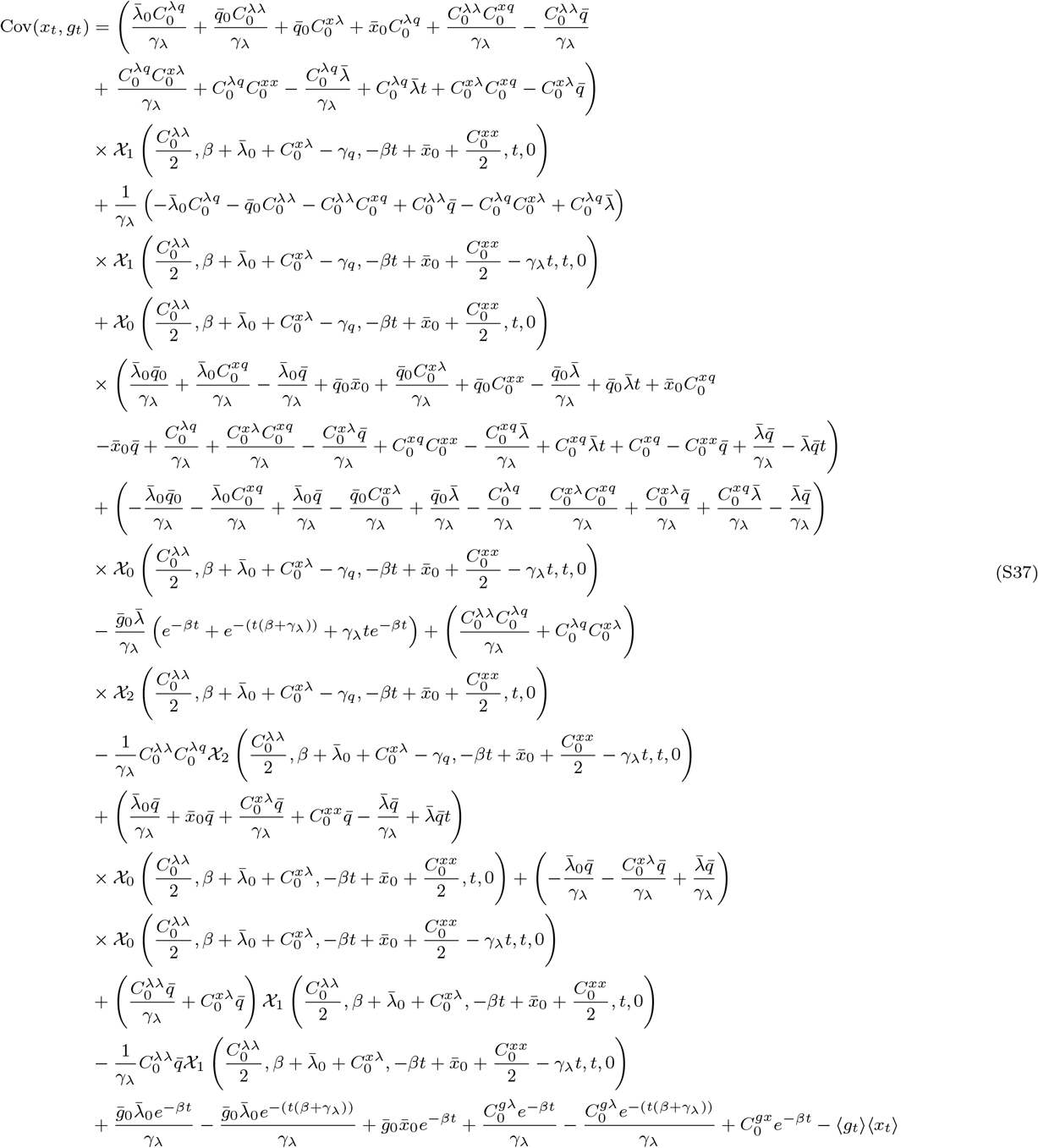

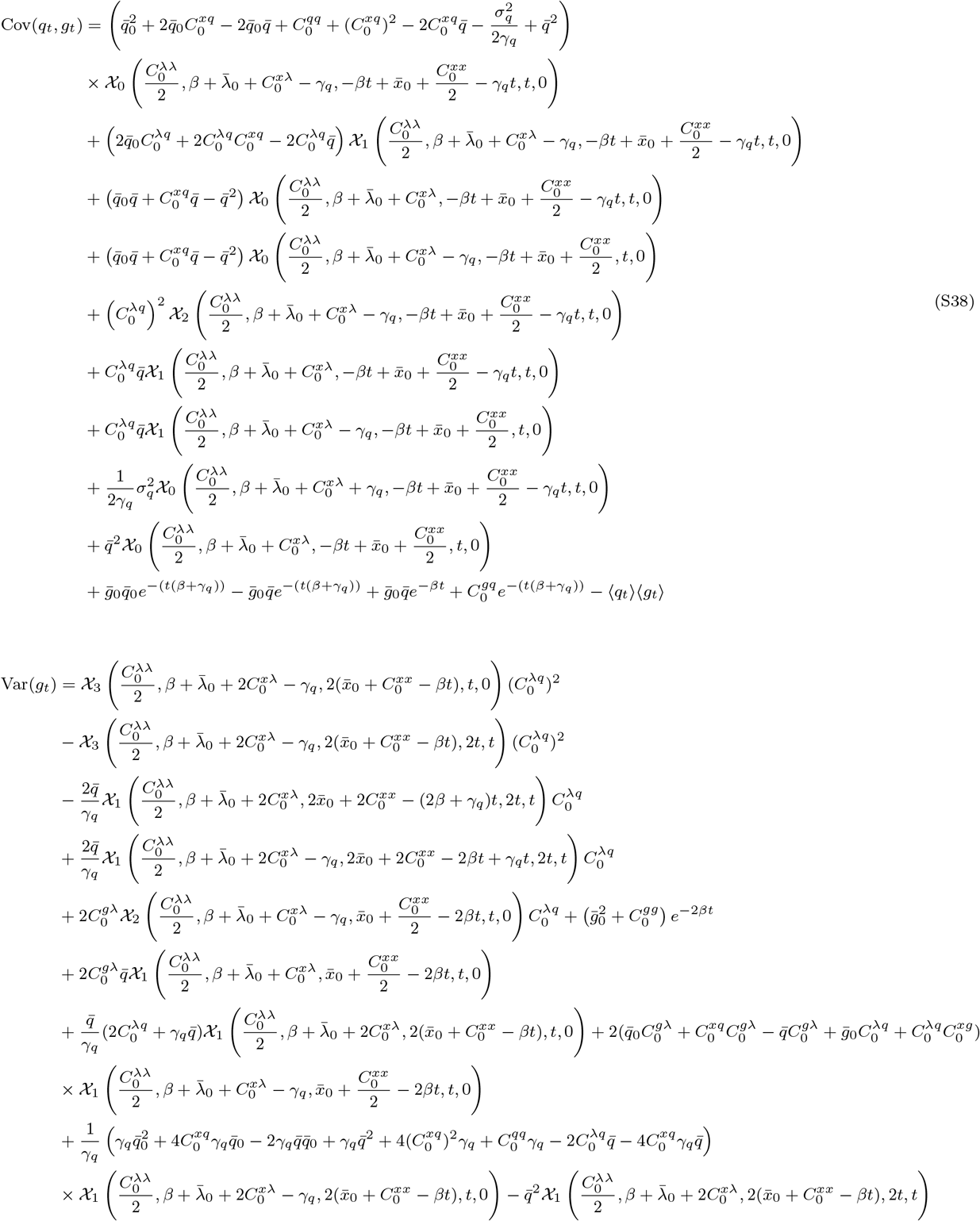

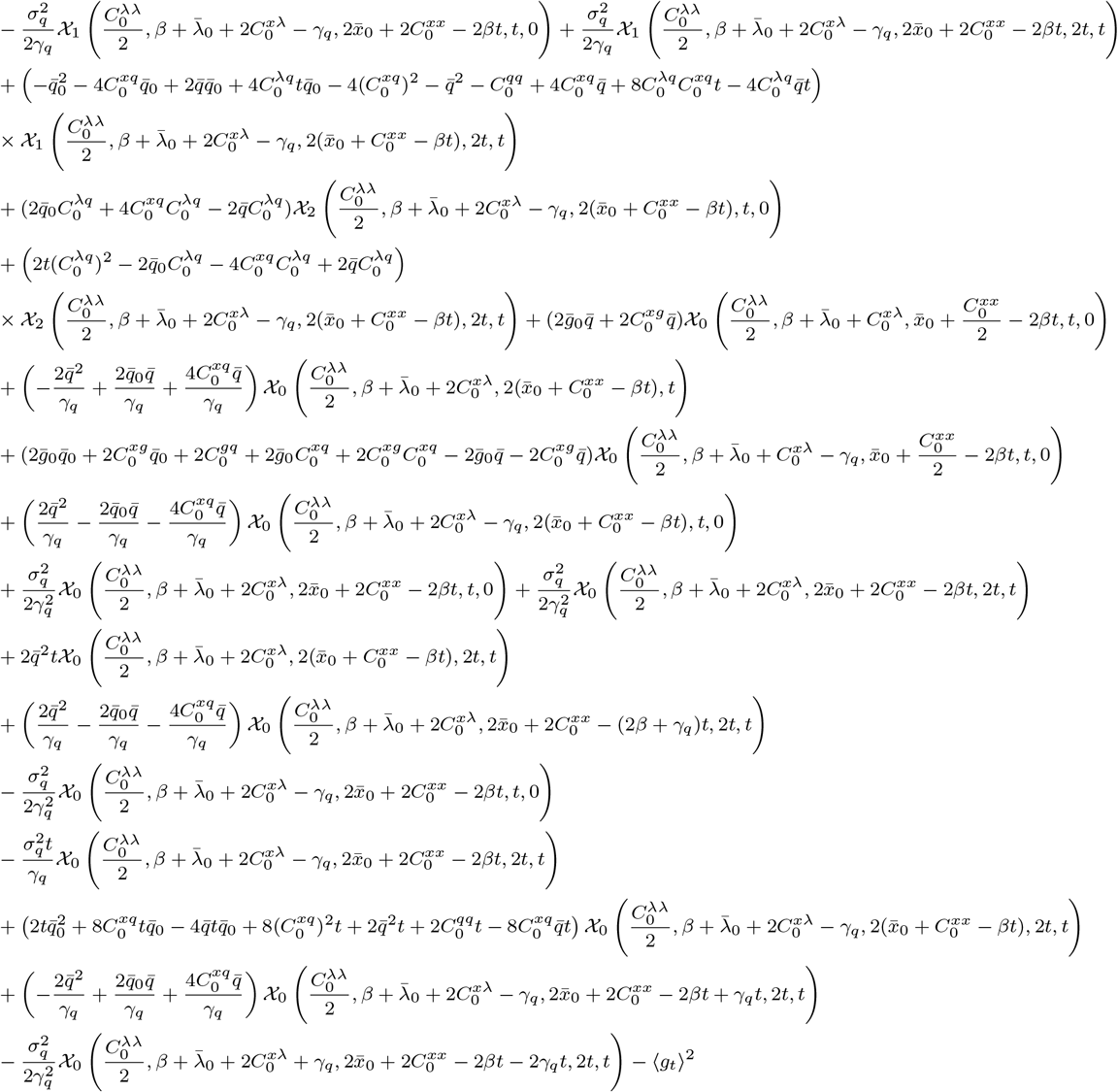

##### E.7.4 Calculating the prior distribution in the backward direction

The decomposition of the posterior distribution requires the calculation of prior and posterior distributions in the backward direction. For that, the time in the model is reversed. The calculation becomes analogous to the forward direction with the parameter transformations

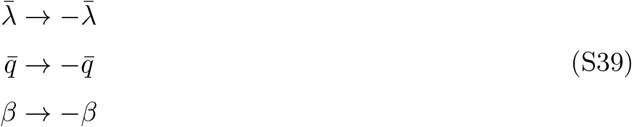

and the variable transformations

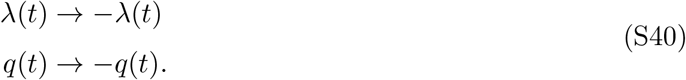

##### E.7.5 Cell division

In case the cell divides between *t*_*n*_ and *t*_*n*+1_, the prior is calculated differently. We simplify the task by assuming the cell grows right up to *t*_*n*+1_ and then divides instantaneously. Thus, after the prior for **z**_*n*+1_ using the model for growth and production is calculated, we calculate a new prior from the former one using the model for cell division. To have a more transparent notation, we will denote the state of the mother cell as **z**_*m*_ and the state of the daughter cell as **z**_*d*_.

In the prior process for the cell division the log size *x* is drawn from a Gaussian centered at *x*_*m*_ *−* log(2), while GFP molecules are binomially distributed to a daughter cell. We ignore correlations between daughter cells. Thus, in the case where both daughter cells are tracked in the experiments, they obtain the same prior distribution after the division and are treated independently afterward.

###### Forwards direction

For cell divisions, we assume that the length of the daughter cell is drawn from a Gaussian distribution. The GFP molecules are binomially distributed with a probability *L*_*d*_*/L*_*m*_ to the daughter cell *d, L*_*d*_ being the daughter cell length and *L*_*m*_ being the length of the mother cell. The means of the prior are:

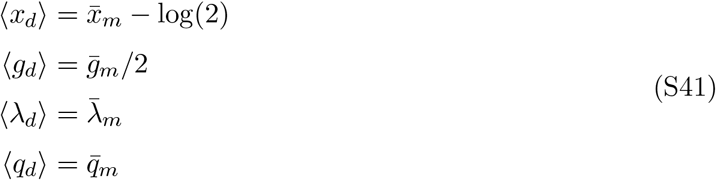

Next, we calculate the covariance matrix of the prior. The conditioned mean of *g*_*d*_ can be written as

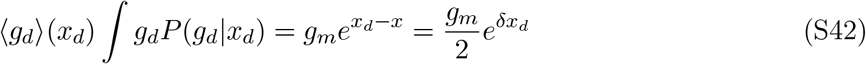

with *δx*_*d*_ = *x*_*d*_ *−* ⟨*x*_*d*_⟩ and using *x*_*m*_ = ⟨*x*_*d*_⟩ + log(2).

Since we model *x*_*d*_ to be drawn from a Gaussian we have 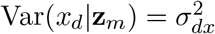 and 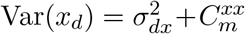. We denote *f* as the probability of a GFP molecule going to the daughter cell of length *L*_*d*_, and thus, 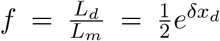 . Since we assume GFP molecules to be binomially distributed, the variance conditioned on the length of the daughter cell reads,

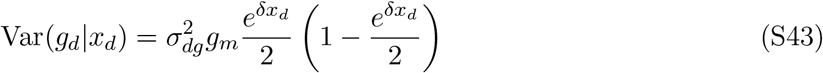

The unconditioned variance becomes,

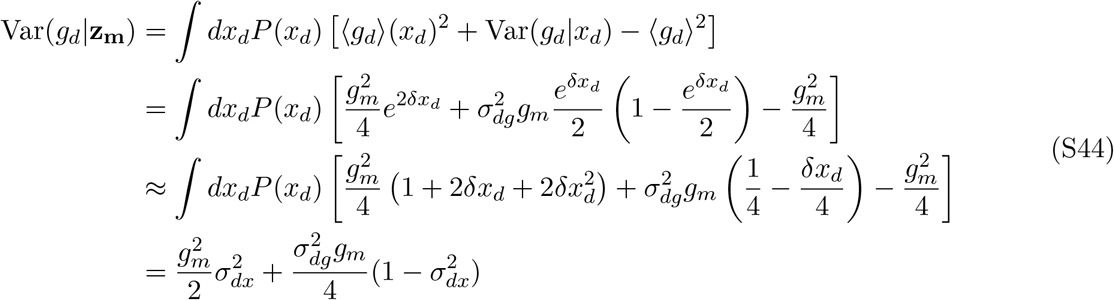

where we linearized exp(*δx*_*d*_) assuming that the deviation of *x*_*d*_ from its mean is small. Then, integrating over the state of the mother cell using Eq. (S28), we obtain

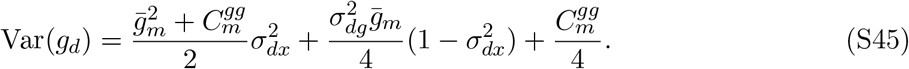

For the covariance we have

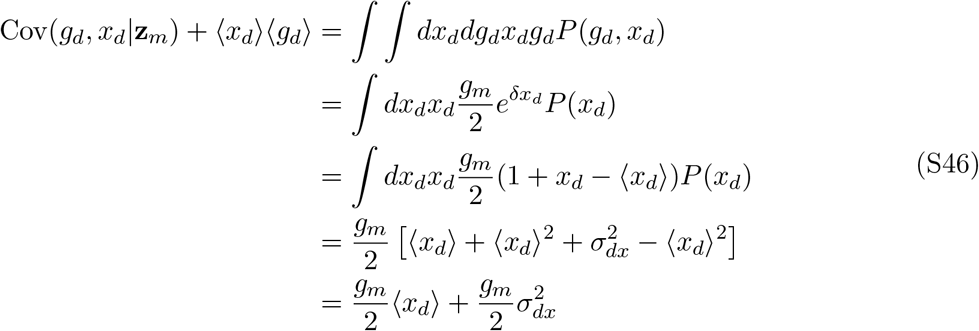

And thus, we obtain using Eq. (S28)

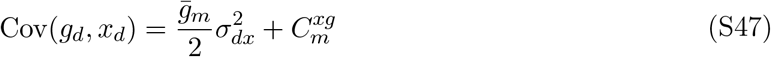

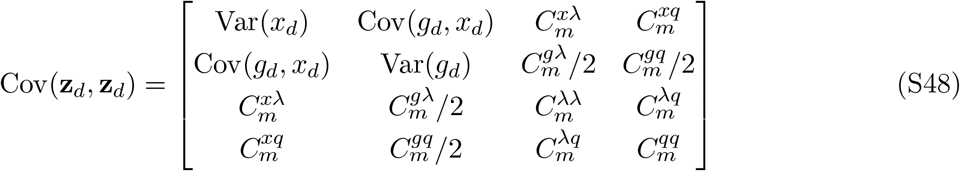

###### Backwards direction

The prior for the backward is calculated similarly to the forward direction, we have,

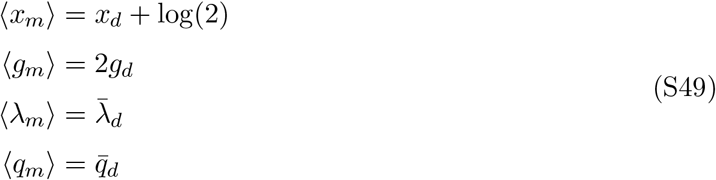

We also have ⟨*g*_*m*_⟩(*x*_*m*_) = 2*g*_*d*_*e*^*δx*^. First, we calculate the variance of *g*_*m*_ conditioned on *g*_*d*_, using a uniform prior we can write

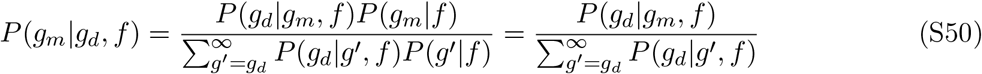

with the binomial distribution

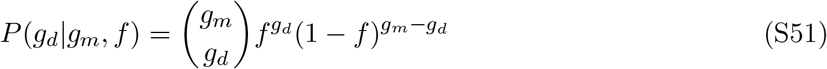

The sum in the denominator becomes

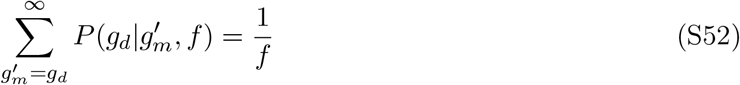

and thus

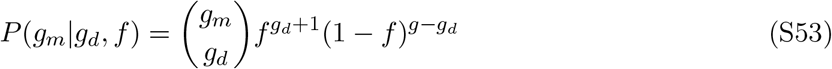

Thus the variance can be calculated as

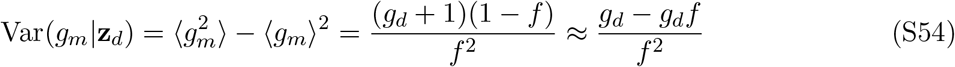

Again, accounting for the fact that the effective number of GFP molecules might differ, we rescale *g*_*d*_ with 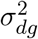 :

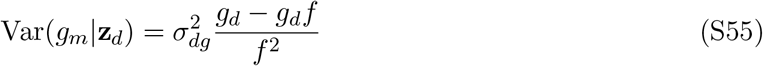

Then, the unconditioned variance using 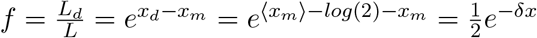

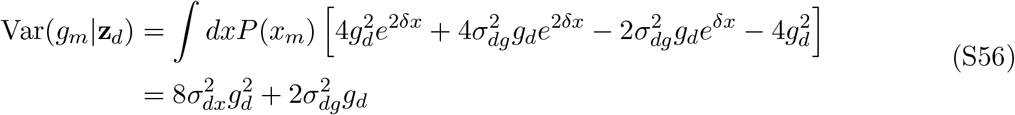

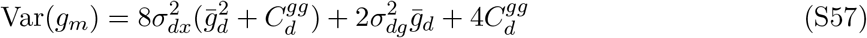

For the covariance elements we have,

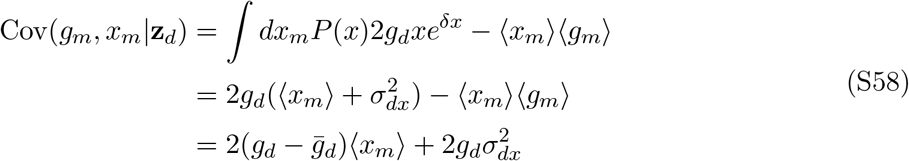

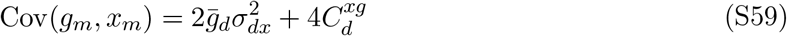

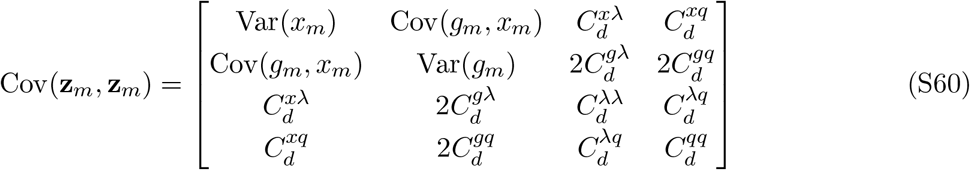

##### E.7.6 Calculation of the cross-covariance elements

To calculate joint posterior distributions, we calculate the cross-covariance terms of the prior,

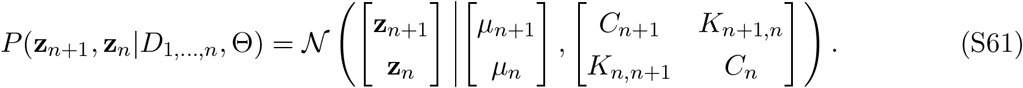

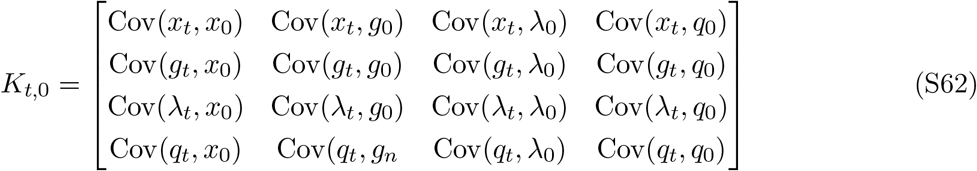

The procedure is similar to the previous calculation: We use Eq. (S28) to find an expression for the unconditioned cross-covariance terms and use Wick’s probability theorem, in case of the element Cov(*λ*_*t*_, *λ*_0_) we find,

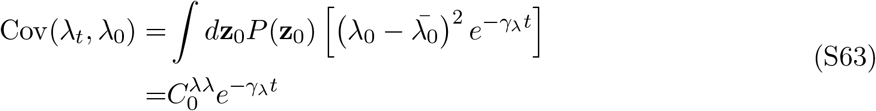

Similarly, we calculate the remaining elements. Again, we use a Mathematical script to aid the calculation of elements involving *g*_*t*_

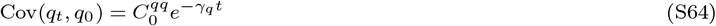

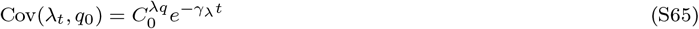

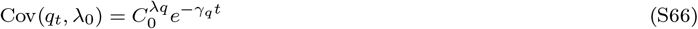

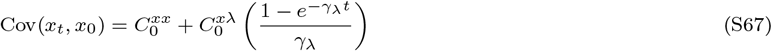

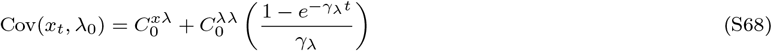

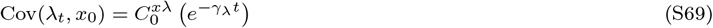

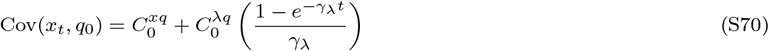

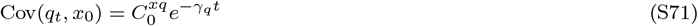

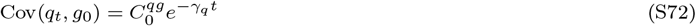

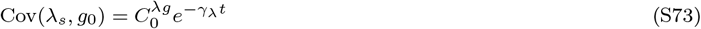

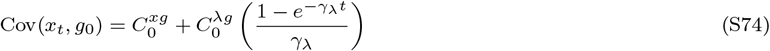

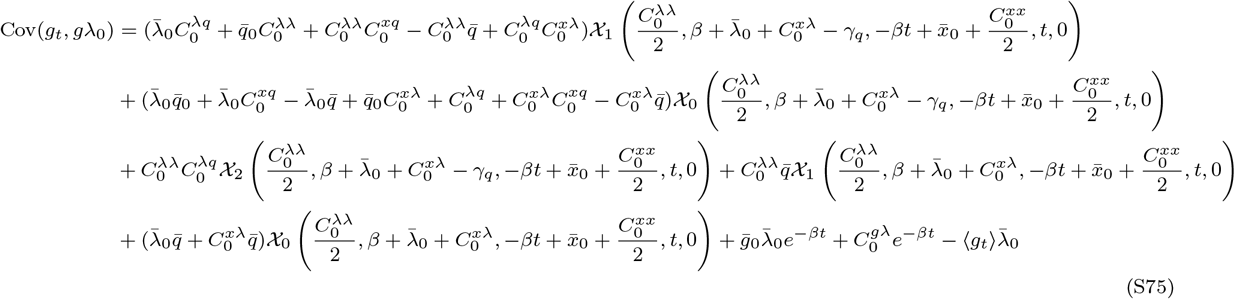

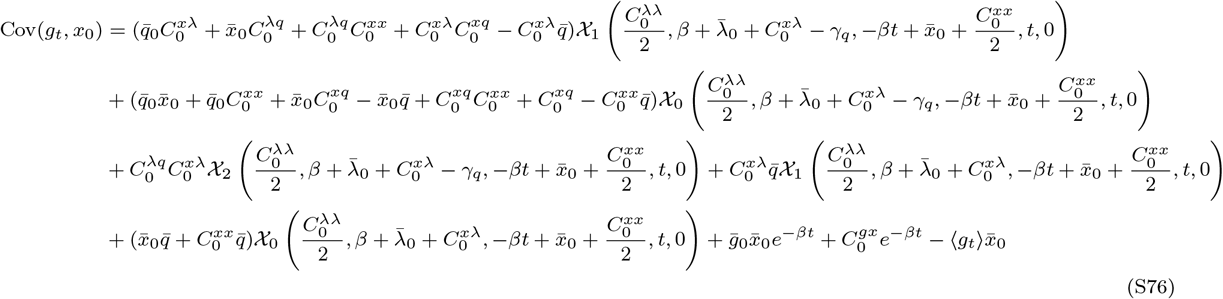

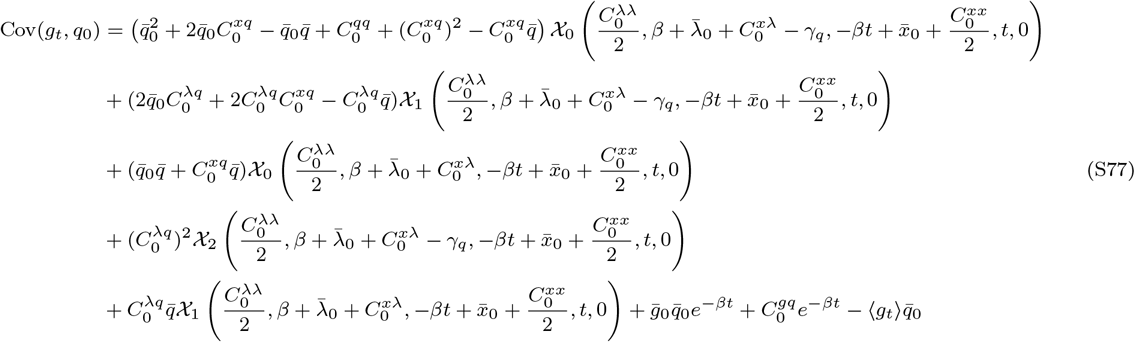

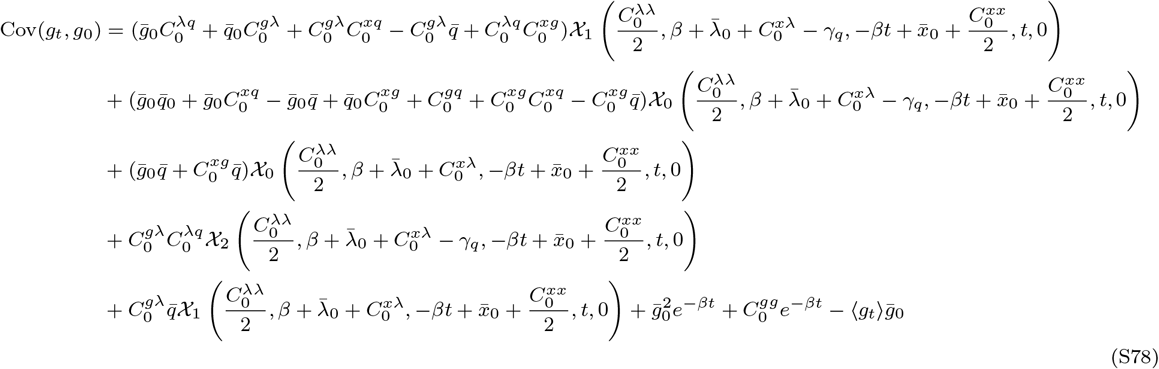

#### E.8 Calculation of the joint probability

Here we show how to calculate the joint posteriors of the states at two time points *n* and *n* + *m*, which can be decomposed analogous to the one-time point posterior distribution (see Sec. E.4),

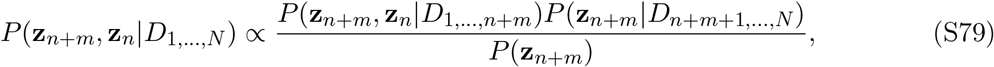

Thus, we first need to calculate the joint over two time points, where we include all measurements up to the later time point to obtain *P* (**z**_*n*+*m*_, **z**_*n*_|*D*_1,…,*n*+*m*_). Then, we include the remaining data points via the term *P* (**z**_*n*+*m*_|*D*_*n*+*m*+1,…,*N*_). The prior *P* (**z**_*n*+*m*_) is chosen as before for the one-time point posterior.

We aim to calculate the joint of non-consecutive time points by performing the integral

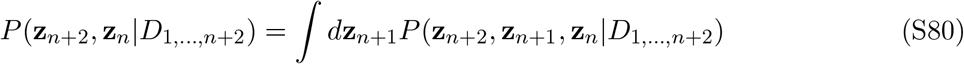

Thus, we calculate the joint probability over three time points, where the last two are consecutive, and integrate over the the intermediate time point, here *n* + 1. The integrand is given by

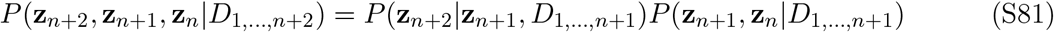

##### Joint probability

We start with the joint probability of **z**_*n*+1_ and **z**_*n*_. The cross terms *K* of the covariance matrix were calculated before in Sec. E.7.6,

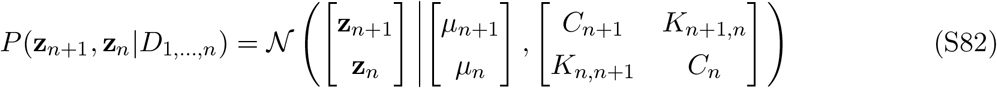

where mean and covariance are conditioned on the data up to *n*.

##### Incorporate measurement

Now, we incorporate **y**_*n*+1_ and calculate *P* (**z**_*n*+1_, **z**_*n*_|*D*_1,…,*n*+1_) via

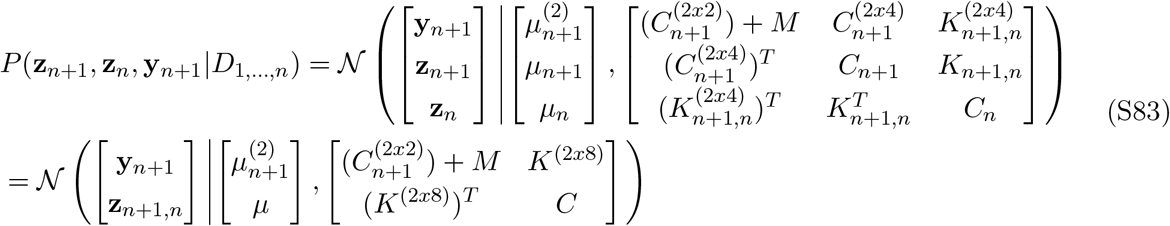

The upper index (2*x*4) indicates that we take the 2*x*4 sub-matrix from the full covariance matrix, i.e. the first two rows corresponding to *x* and *g*. Then, we can directly read off the conditional *P* (**z**_*n*+1_, **z**_*n*_|*D*_1,…,*n*_, **y**_*n*+1_) = *P* (**z**_*n*+1_, **z**_*n*_|*D*_1,…,*n*+1_), which updates the mean and the covariance of the joint over **z**_*n*+1_, **z**_*n*_ and we will drop the explicit conditioning of the mean and the covariance from now on, since we incorporated all the measurements for now. We obtain,

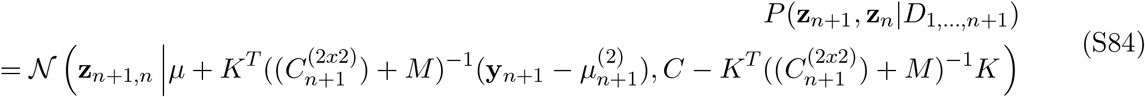

To solve the integral later, we separate the marginal and the conditional probability via

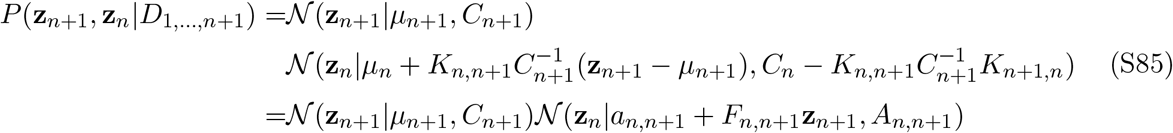

with *a*_*n,n*+1_, *F*_*n,n*+1_, *A*_*n,n*+1_ given by,

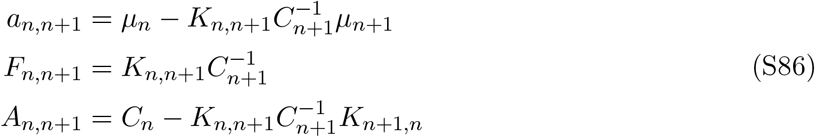

We will make use of these definitions again.

##### Conditional probability

We obtain the the conditional *P* (**z**_*n*+1_|**z**_*n*_, *D*_1,…,*n*_) by writing

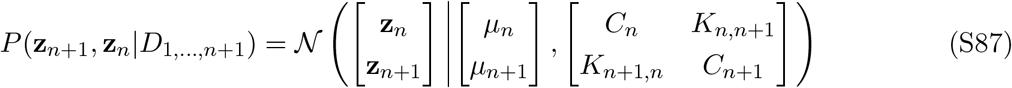

with the conditional

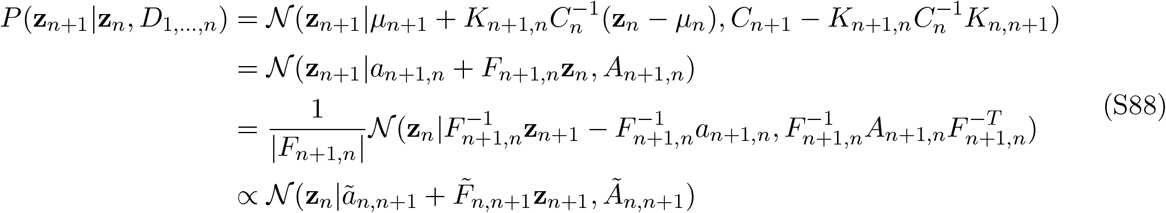

*a*_*n*+1,*n*_, *F*_*n*+1,*n*_ and *A*_*n*+1,*n*_ are defined as before, although the indices are reversed, and the transformation in the last step does not change the form of the Gaussian. It is important to note, that the means and covariances are not conditioned on the measurement **y**_*n*+1_ yet.

##### Joint distribution of three time points

To move forward we write down the joint over three time points and then integrate over the intermediate time point,

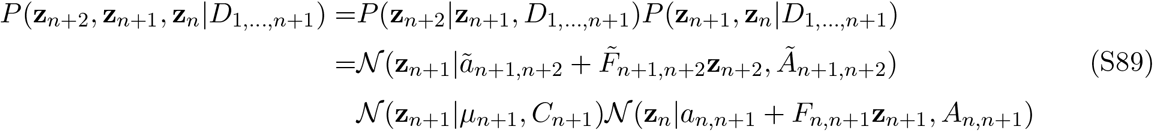

First, we multiply the two distributions over **z**_*n*+1_,

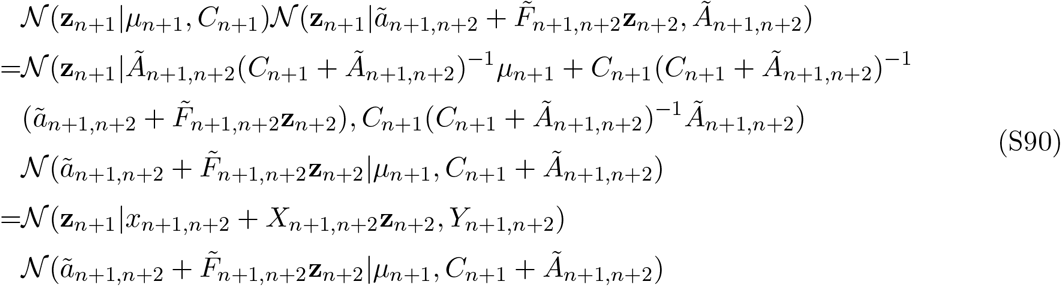

with

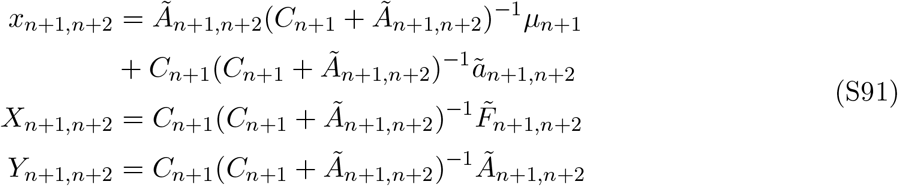

Thus, the joint over **z**_*n*+2_, **z**_*n*+1_, and **z**_*n*_ reads,

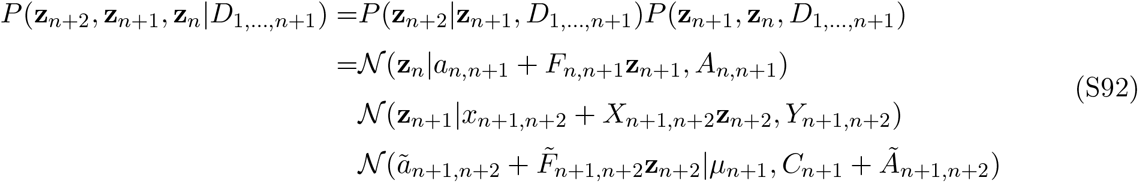

This expression has two Gaussians including **z**_*n*+1_, which can be integrated. The integral over **z**_*n*+1_ is solved by

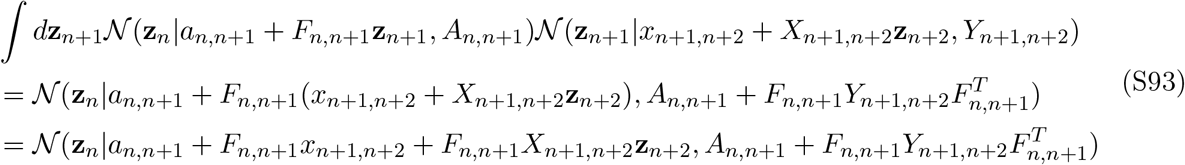

We also rewrite the Gaussian that only depends on **z**_*n*+2_

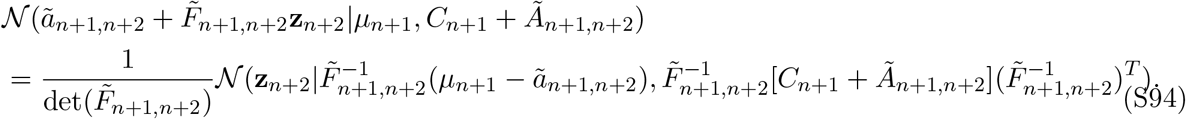

Together with the solution of the integral, we can write the entire expression as

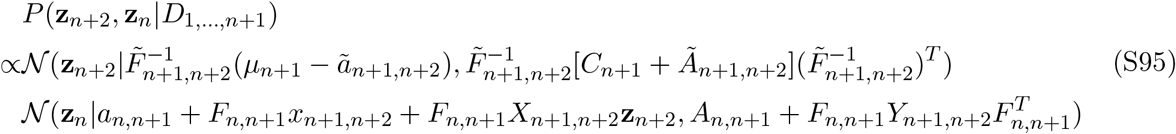

That expression can, of course, be rewritten as a single Gaussian.

Finally, we need to include **y**_*n*+2_ and thus move from *P* (**z**_*n*+2_, **z**_*n*_|*D*_1,…,*n*+1_) to *P* (**z**_*n*+2_, **z**_*n*_|*D*_1,…,*n*+2_). This calculation is identical to the calculation above (**Incorporate measurement**). Furthermore, we need to multiply by *P* (**z**_*n*+*m*_|*D*_*n*+*m*+1,….,*N*_), which was already calculated for the single time point posterior distributions.

#### E.9 Estimating means and variances from posterior distributions

We often want to calculate the mean and variance of some cell state variable *u* over some set of observations, e.g. the growth rate at a particular point in the cell cycle. To do this properly, we need to take into account the error bars on the estimates *u*_*i*_. For example, the estimated mean *µ*_*u*_ of a set of *N* observations *u*_*i*_ is given by

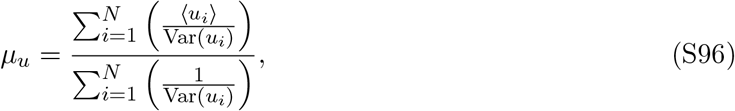

where ⟨*u*_*i*_⟩ is the mean of the posterior for *u*_*i*_ and Var(*u*_*i*_) its variance.

Similarly, the variance of the *N* observations *u*_*i*_ is estimated by

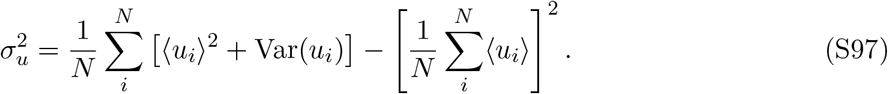

#### E.10 Maximum likelihood estimate for the auto-covariance function

We want to estimate correlations between cell state variables at different time points. For example, we might want to estimate the correlation between the current growth rate and the volumic production rate some time later, as in Fig. S11. More generally, we want to calculate the correlation between some state variable *u* and another state variable *v* at a certain time *t* in the future.

Let (*u*_*i*_, *v*_*j*_) denote pairs of observations that are a time *t* apart and assume that there are *a* different ‘ancestor’ observations *u*_*i*_ and that for observation *u*_*i*_ there are *d*_*i*_ ‘descendant’ observations a time *t* into its future. One might simply estimate the correlation between these pairs of observations by calculating the Pearson correlation function over all these pairs, but there are two problems with this approach. First, we do not know the exact values (*u*_*i*_, *v*_*j*_), but we only have joint posterior distributions for these pairs. Second, different ancestor observations *u*_*i*_ may have different numbers of descendants *v*_*j*_, and it is unclear how we should ‘weigh’ these in the calculation of the correlation.

We thus take the following approach, which, while still *ad hoc*, we believe is more accurate. We will assume that the pairs of observations (*u*_*i*_, *v*_*j*_) can be modeled as deriving from a multivariate Gaussian distribution. This distribution is described by the means *µ*_*u*_ and *µ*_*v*_, the variances 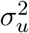 and 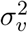, and finally the correlation *r*. That is, we write

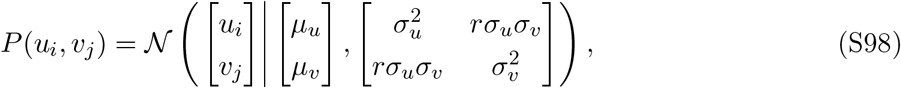

where *r* is the correlation between these variables.

As explained in section E.9, we can calculate the means *µ*_*u*_ and *µ*_*v*_, and the variances 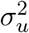 and 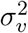 by averaging over all observations of the variables *u* and *v*. For example, if *u* is growth rate and *v* volumic production, we would estimate the mean and variance of both growth rate and volumic production by taking averages over all observations in the dataset.

The only unknown variable in our multivariate Gaussian of the pairs (*u*_*i*_, *v*_*j*_) is thus their correlation. We will now estimate this correlation by maximizing the log-likelihood *L*(*r*) of all observations (*u*_*i*_, *v*_*j*_) with respect to *r*.

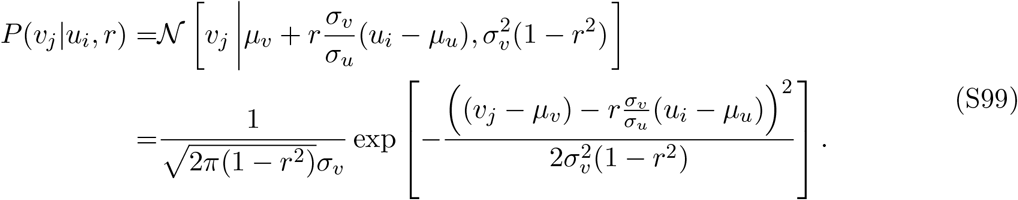

We can use this expression for the conditional probability to calculate the likelihood of all pairs (*u*_*i*_, *v*_*j*_) as a function of *r*

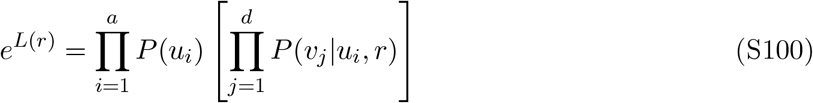

The log-likelihood *L*(*r*) then becomes

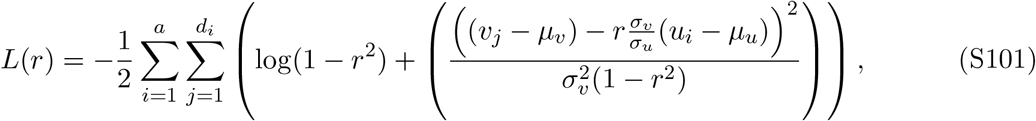

where we have only kept terms that depend on *r*.

Note that this expression still depends on the values (*u*_*i*_, *v*_*j*_) for which we have joint posterior distributions (as calculated above). We use these posterior distributions over the values of the pairs (*u*_*i*_, *v*_*j*_) to calculate the *expected* log-likelihood ⟨*L*(*r*)⟩.

The result can be expressed most easily by defining

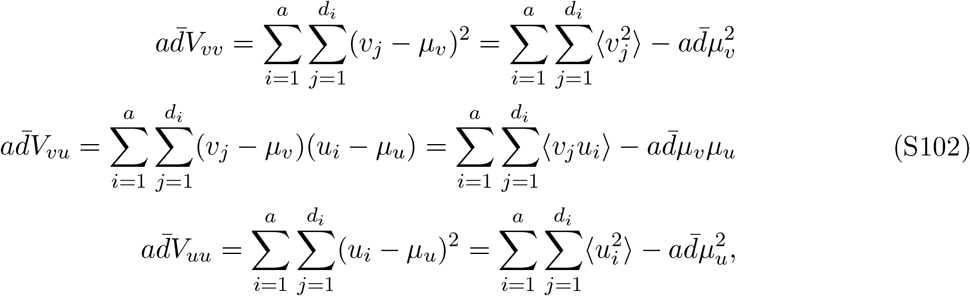

where 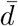 is the average number of descendants per ancestor. We then have

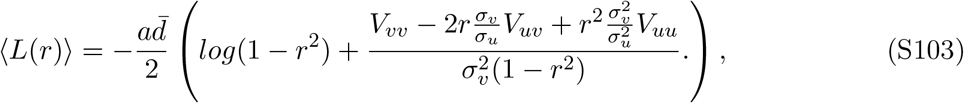

where we have again included only terms that depend on *r*. We then find the correlation *r* as the maximum of the likelihood function ⟨*L*(*r*)⟩. An error-bar on *r* can be calculated as the square root of the inverse of the Hessian of this log-likelihood, which reads,

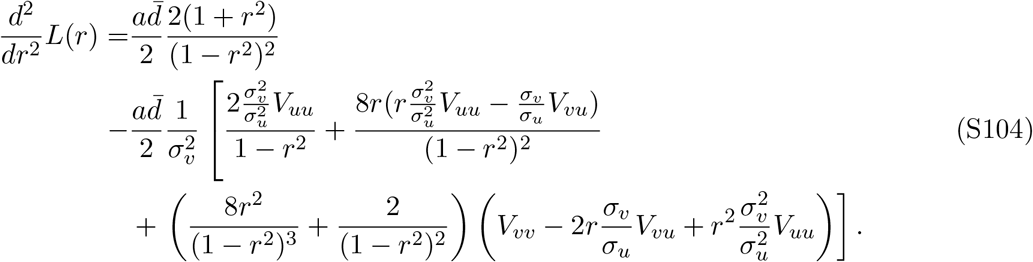

### F Promoter sequences

#### hi1

CAGTCAGTCCACGTGTGGCGGGAATCCCTTGGCGGCGGCCCTGCTT

CTGGTGTATACTGTTATGGAAACCAAATCTACGAGCCTACAGGATT

CCTATCCGGGATCCTCTGGATGTAAGAAGGAGATATACAT**atg**

#### hi3

CAGACGCTGGGTCGGCTGTCGGTCAGGTAGAGTCAGCTCTTACGAT

CTCTTGACATACTCAACCGGATCGTGTAATGGTTCAAAGTGCGTGA

TTTTAATGTTTTCGAGACGCGGAGGGTGCCTTCATAAGGGATCCTC

TGGATGTAAGAAGGAGATATACAT**atg**

#### rrnB

CGCCAGGAGCTGAACAATTATTGCCCGTTTTACAGCGTTACGGCTTCGA

AACGCTCGAAAAACTGGCAGTTTTAGGCTGATTTGGTTGAATGTTGCGC

GGTCAGAAAATTATTTTAAATTTCCTCTTGTCAGGCCGGAATAACTCCC

TATAATGCGCCACCACTGACACGGAACAACGGCAAACACGCCGCCGGGT

CAGCGGGGTTCTCCTGAGAACTCCGGCAGAGAAAGCAAAAATAAATGCT

TGACTCTGTAGCGGGAAGGCGTATTATGCACACCCCGCGCCGCTGAGAA

AAAGCGAAGCGGCACTGCTCTTTAACAATTTATCAGACAATCTGTGTGG

GCACTCGAAGATACGGATTCTTAACGTCGCAAGACGAAAAATGAATACC

AAGTCTCAAGAGTGAACACGTAATTCATTACGAAGTTTAATTCTTTGAG

CGTCAAACTTTTAAATTGAAGAGTTTGATCATGGCTCAGATTGAACGCT

GGCGGCAGGCCTAACACATGCAAGTCGAACGGTAACAGGAAGAAGCTTG

CTTCTTTGCTGACGGGATCCTCTAGATTTAAGAAGGAGATATACAT**atg**

#### rplN

CGTCCGCTGTCCAAGACTAAATCCTGGACGCTGGTTCGCGTTGTAGAGA

AAGCGGTTCTGTAATACAGTACACTCTCTCAATACGAATAAACGGCTCA

GAAATGAGCCGTTTATTTTTTCTACCCATATCCTTGAAGCGGTGTTATA

ATGCCGCGCCCTCGATATGGGGATTTTTAACGACCTGATTTTCGGGTCT

CAGTAGTAGTTGACATTAGCGGAGCACTAAAATGATCCAAGAACAGACT

ATGCTGAACGTCGCCGACAACTCCGGTGCACGTCGCGTAATGTGTATCA

AGGTTCTGGGTGGCTCTCGAGAGATCCTCTAGATTTAAGAAGGAGATAT

ACAT**atg**

## Notes

https://github.com/nimwegenLab/RealTrace

### Competing Interest Statement

The authors have declared no competing interest.

